# Ancient DNA Reveals Hominoid Evolution: Intermediate DNA Sequences and Advances in Molecular Paleontology

**DOI:** 10.1101/2025.04.27.650833

**Authors:** Li-Juan Zhao, Shu-Jie Zhang, Zhan-Yong Guo, Qing-Yang Zhong, Yong-Kai Zheng, Yu-Qin Cai, Chao-Dong Jia, Shu-Hui Zhang, Rui Mao, Cai-Yun Hong, Min-Zhi Wu, Yong-Kai Wang, Zhi-Fang Zheng, Yun Zhang, Yu-Xuan Jin, Wan-Qian Zhao

## Abstract

Fossils and ancient crude pottery vessels function as physical “DNA containers,” preserving oriDNA (original, *in situ* DNA) and accumulating eDNA (environmental DNA) over time. Ancient DNA (aDNA) serves as molecular fossils, chronicling evolutionary history. Using “nano-affinity bead technology”, we extracted and sequenced DNA from *Lycoptera* fossils in the Jehol region and a “round-bellied jar (Jar)” from the Erlitou period in Guangwu Town, yielding 236,545 primate sequences from the fossils and 86,908 from the pottery. We observed that the AFF value of DNA sequences negatively correlates with species divergence time^1^, offering a quantitative measure of DNA preservation and host divergence. Some fragments distinct from modern genomic sequences, termed “intermediate DNA sequences” (IDS), have been identified. Many IDS exhibit an upper age limit, preserving characteristics of the last common ancestor (LCA) of the Hominidae and offering molecular insights into “Darwin’s puzzle”. Among IDS, the SRRA subtype (Sequence Reversal and Rearrangement), identified in mRNA-coding exonic sequences, arises from the incorporation of a complementary antisense strand upstream of the sense strand, either adjacent to it or separated bya sequence interval. This introduces a novel post-transcriptional regulatory mechanism at the mRNA level, driven by SRRA, which presets hairpin structures and Indels in the UTR or CDS of exonic sequences, modulating gene variation. We propose: “SRRAs played a ‘critical trial-and-error’ role in early Hominidae evolution, facilitating adaptive genomic changes, with some SRRA sequences later excised from exons”, positioning SRRAas an evolutionary “genetic switch”. Additionally, seven species -specific fragments (SSFs) of non-human primates (NHPs) linked to Asian *Homo erectus* were identified in the fossils, and pottery DNA revealed sequences from tropical species (e.g., zebrass, oil palms), providing evidence of climate-driven local extinction and supporting paleo-ecological and paleo-environmental reconstruction. This method of analyzing aDNA from non-skeletal materials opens new avenues in paleontology, archaeology, and geology, guiding the tracing of ancient migration patterns and fossil searches. DNAfragments preserved within “DNA containers ” exhibit an “old-few, new-many” turnover pattern, with many aDNA fragments displaying non-deamination. This evidence challenges prevailing perspectives, the authenticity criteria for aDNA, and the capabilities and scope of the traditional research method, necessitating a thorough reevaluation of the relevant knowledge framework. Furthermore, this study opens new opportunities for frontier research.

## Introduction

Porosity of Fossils and DNA Dynamics: Many fossils, as lithic structures, are naturally porous, featuring cavities, pores, and tunnels that connect to the external environment, making them DNA containers. Vertebrate skeletal fossils, for example, possess a trabecular, spongy architecture filled with interstices and voids. *Lycoptera* fossils, formed in volcanic sedimentary rock approximately 120 million years ago (Ma), exhibit an internal volume ratio of 10.4–11.3%^1^. Similarly, Neolithic coarse pottery artifacts, such as the “Jar” from ∼3,700 years ago, display comparable porosity with an internal volume ratio of 10.5% (Materials and Methods). These containers experience cyclical moisture fluctuations in soil: water content diminishes during dry seasons and is replenished during rainfall. eDNA flows in and out with water, where mineral salts and lipids aid its deposition and preservation. Through repeated drought-precipitation cycles, fossil DNA undergoes exchange with external eDNA, resulting in a molecular profile dominated by more recent sequences. As oriDNAdiminishes, eDNAaccumulates, forming a Genome Accumulated over Time (GAT genome) that integrates both *in situ* and eDNA. Locally dominant species primarily contribute to the eDNA. In human settlements, burial practices sustain a subsurface influx of human DNA—a key eDNA component—alongside fragments from livestock, hunted animals, and crops, rendering the GAT genome a dynamic record of ecological shifts over time.

Elucidating Darwin’s Puzzle of Transitional Species through IDS: “Darwin’s puzzle,” namely the issue of transitional species, refers to the relative scarcity of intermediate forms in the fossil record. Intermediate species exhibit morphological and functional characteristics that lie between those of ancestral and derived taxa. While some fossil evidence suggests gradual changes in species traits, the evolution of certain species does not follow a linear or uniform trajectory; instead, it is characterized by prolonged periods of stability followed by rapid transformations. The “punctuated equilibrium” hypothesis, proposed by Eldredge and Gould in 1972, seeks to account for this phenomenon ^2^. Regardless of the rate of evolution, the evolutionary process fundamentally relies on genomic varia tion. Byanalyzing IDS that align with primate genomes and determining their formation timing, we can glean insights into the relationship between genetic mutations and evolution, adding more details to the divergence history of Hominidae species. The IDS identified in this study are absent from the genomes of extant species (AFF < 100%), serving as evidence of mutations that once occurred in ancient genomes. These accumulated mutations may influence the development of morphology and function by regulating morphogenetic genes. We suggest that, although some lineages lack morphologically intermediate species, their genomes are not deficient in IDS, which accumulate over time and ultimately drive evolutionary change. This study partially resolves “Darwin’s puzzle” at the molecular level.

DNA Extraction, Analysis, and Inferences on Primate Evolution: Utilizing nanoparticle affinity bead technology, we extracted and sequenced DNA from porous containers, recovering a total of 236,545 primate fragments from *Lycoptera* fossils and 86,908 human pieces from the “Jar” through the Best E-value mode analysis. By examining the types and quantities of IDS within the fossil GAT genome, we constructed an “association curve of species and corresponding AFF”^1^ (Materials and Methods). This curve delineates the temporal variation patterns of primate genome sequences accumulated in the environment, spanning from the formation of the container to its burial and preservation. Validation of these DNA fragments using the MS mode facilitated the identification of several fossil SSFs unique to NHPs and extinct species of mammals and plants preserved within the “Jar”. Based on the evidence, we infer that the fossil site region was once inhabited by Asian *Homo erectus*, also the SSFs reveal a distinct pattern of extinct species associated with the ecological context of the “Jar” region, providing insights into the historical biodiversity of the area. Additionally, SRRA, a subtype of IDS, is hypothesized to have played a “critical trial-and-error” role in the early evolution of great apes, conferring diverse post-transcriptional regulatory mechanisms on mRNA. These findings contribute to our understanding of primate evolution by providing molecular evidence of historical species distribution and genomic dynamics (Figure S1).

## Results

### I. Distribution of the Fossil DNA Sequences Across Primate Species

1. Species Related to Sequence Matches: Using the Best E-value mode, we identified 236,545 primate-related sequences in fossil DNA. Of these, 228,703 matched the modern human (*Homo sapiens*) genome, 6,727 aligned with the chimpanzee (*Pan troglodytes*) genome, 586 corresponded to the gorilla (*Gorilla gorilla*) genome, 208 matched the orangutan (*Pongo*) genome, 103 aligned with the gibbon (*Hylobates*) genome, 26 corresponded to the crested gibbon (*Nomascus*) genome, 58 matched the leaf monkey (*Trachypithecus*) genome, 96 aligned with the macaque (*Macaca*) genome, and 37 corresponded to the capuchin (*Sapajus*) genome (Figure 1A).
2. Variation in AFF Across Species Evolutionary Order: In *Homo sapiens*, *Pan troglodytes*, and *Gorilla gorilla*, DNAsequence proportions peak at AFF = 100%, followed by 99% ≤ AFF < 100%, then 95% ≤ AFF < 99%, with the lowest proportion at AFF < 60%. In *Pongo*, *Hylobates*, *Nomascus*, and *Trachypithecus*, the proportion of sequences with AFF = 100% decreases, while those with AFF < 60% increase significantly. In *Macaca* and *Sapajus*, sequences with AFF = 100% diminish further, with AFF < 60% predominating (Figure 1A). The AFF value, reflecting sequence variation, indicates that earlier-evolving species exhibit fewer high-AFF fragments.
3. Distribution of AFF Values for Primate-Related Sequences: DNA fragments are distributed across AFF values as follows: 170,245 at AFF = 100%, 36,860 between 99% and 100%, 21,291 between 95% and 99%, 2,984 between 90% and 95%, 4,070 between 60% and 90%, and 1,095 below 60% (Figure 1B). These data indicate a significant decrease in DNA amount associated with primates as the AFF declines, which supports an “old-few, new-many” pattern over time. This aligns with our hypothesis that fossils, as containers, enable a dynamic eDNA cycle of influx and efflux, progressively replacing older sequences with newer ones.
4. SSFs in NHPs: Using the MS mode (Materials and Methods), we identified seven SSFs unique to non-homo primate genomes: three from *Pan troglodytes*, one from *Pongo*, one from *Hylobates*, and two from *Macaca* (Figure 1A). All SSFs exhibit AFF values below 100%, reflecting significant divergence from modern genome sequences.

**Figure 1A.**
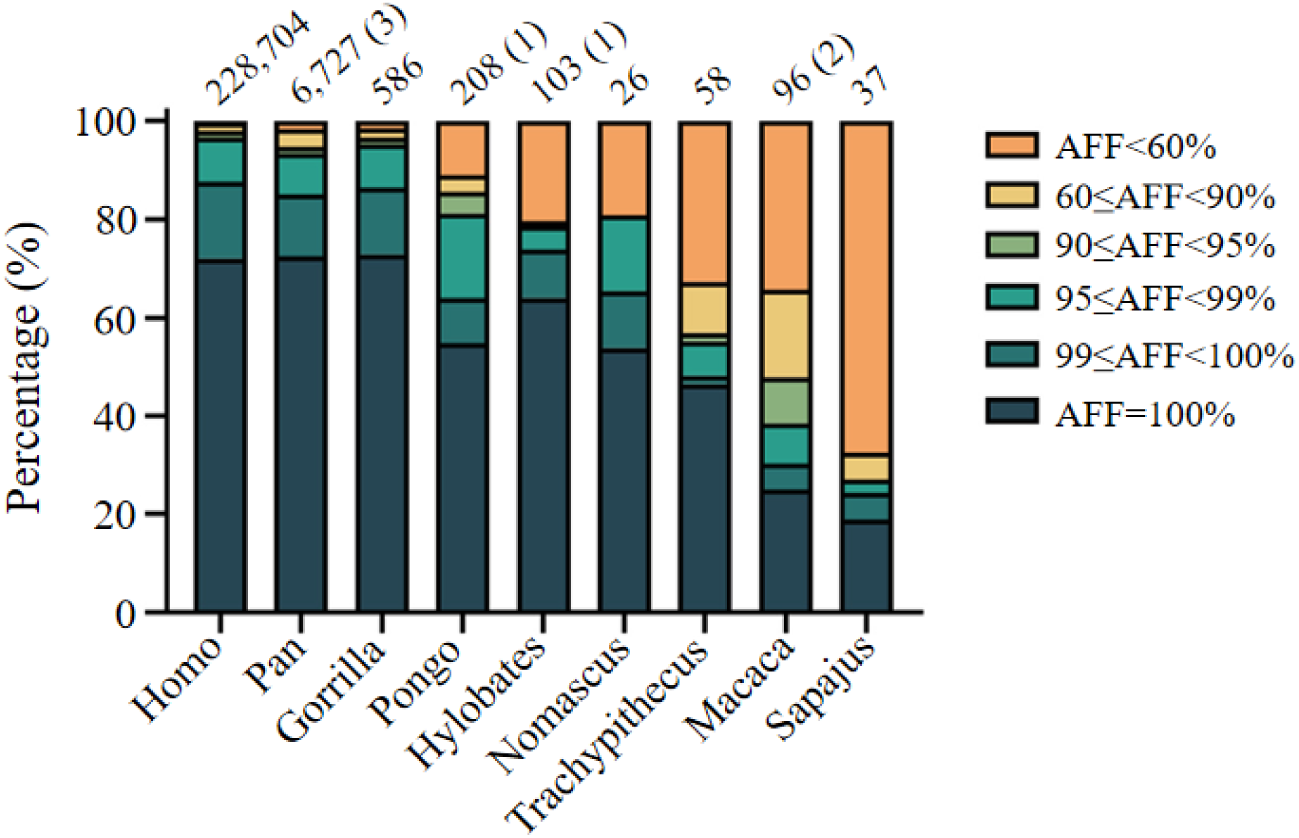
Distribution of AFF values for DNA fragments from *Lycoptera* fossils detected via the Best E-value mode and aligned w ith the genomes of primate species. Numbers indicate the total count of DNA fragments mapped to each species; numbers in parentheses represent the count of SSFs.

**Figure 1B.**
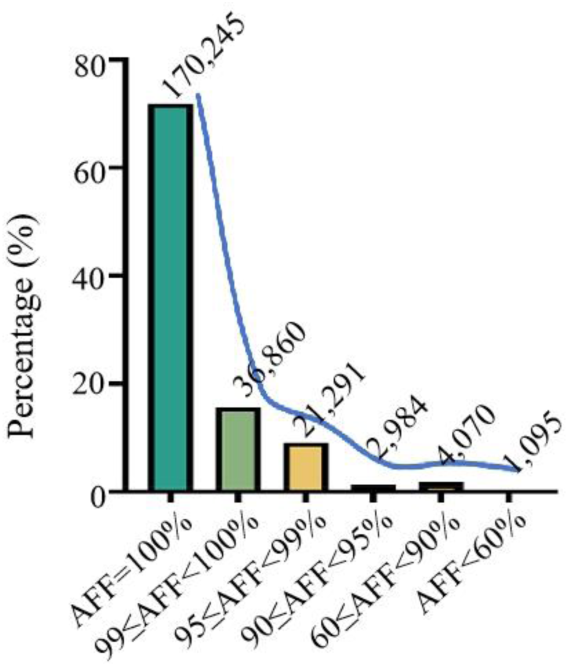
AFF descent rate curve of DNA fragment counts for primate species in *Lycoptera* fossils. Numbers represent the quantity of DNA fragments mapped to primate species w ithin corresponding AFF intervals, as detected by the Best E-value mode.

### II. IDS: Traces of Evolution in the LCA Genome

1. Classification of IDS: IDS fragments are class ified into three types: MNV (sequences with multi-nucleotide variations), Indels (fragments featuring nucleotide variations with insertions or deletions), and SRRA (fragments with partial sequence inversions and rearrangements). SSFs, matching only a single species, may internally harbor MNV, Indels, or SRRA.
2. Inferring the Upper Limit of IDS Emergence: We constructed a phylogenetic tree with evolutionary nodes for the genus Homo, Homininae, Hominidae, Hominoidea, Old World monkeys (Cercopithecoidea), and New World monkeys (Platyrrhini). Using Nucleotide Blast, we matched DNA sequences to species and integrated the results into the tree for analysis, enabling us to infer the upper age limit of IDS occurrence (Materials and Methods).
3. MNVs: This category comprises 18 fragments (Table S1). For instance, ID_014 primarily matches the Homininae, with equivalent alignments to *Homo sapiens*, *Pan troglodytes*, and *Gorilla gorilla* (E-value: 1e-62), followed by *Pongo abelii* (E-value: 3e-58). ID_014 diverges from human, chimpanzee, and gorilla genomes by four variant sites and from *Pongo abelii* by an additional two sites and a 3 bp deletion. Absent from modern genomes, ID_014’s upper age limit is estimated at 15.2 Ma based on phylogenetic relationships (Figure 2A).
4. Indels: This category comprises 155 fragments (Table S1). For example, ID_068 primarily matches the Homininae, with equivalent alignments to *Homo sapiens*, *Pan troglodytes*, and *Gorilla gorilla* (E-value: 2e-23), followed by *Pongo abelii* (E-value: 1e-21). Relative to human, chimpanzee, and gorilla genomes, ID_068 features a 14 bp insertion; compared to *Pongo abelii*, it includes an additional single-point mutation. Absent from modern genomes, ID_068’s upper age limit is estimated at 15.2 Ma based on phylogenetic relationships (Figure 2B).
5. SRRAs: Fossil aDNAcontains three SRRA fragments (Table S1).

**Figure 2A.**
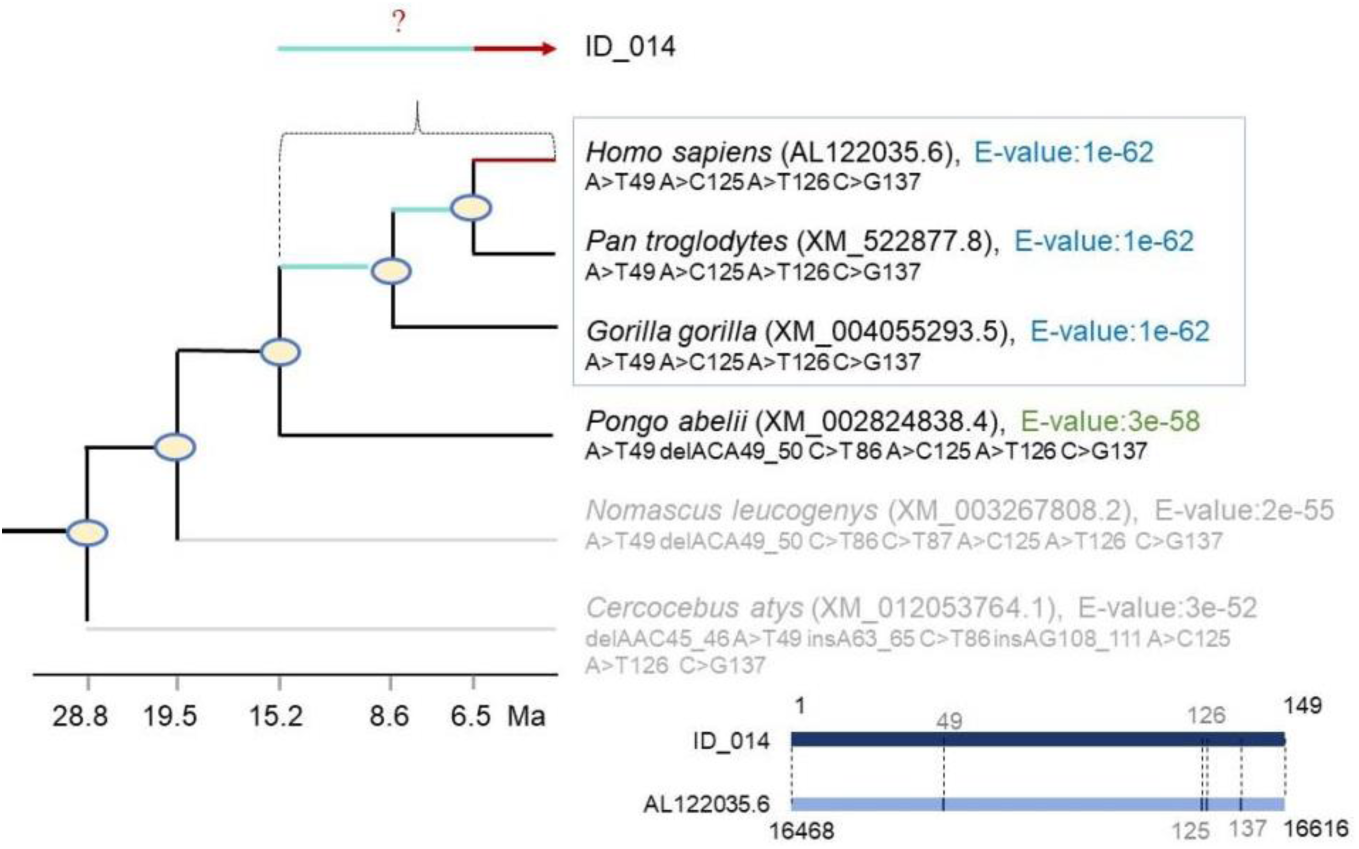
Illustration of multiple nucleotide variations (MNVs) across the mult-loci. The ID_014 sequence w as aligned w ith sequences from Hominoidea genomes, w ith corresponding results shown. Ellipses represent phylogenetic nodes. A>T49 indicates that the 49^th^ base “T” in the ID_014 DNA sequence corresponds to base “A” in the *Homo sapiens* sequence (AL122035.6).

**Figure 2B.**
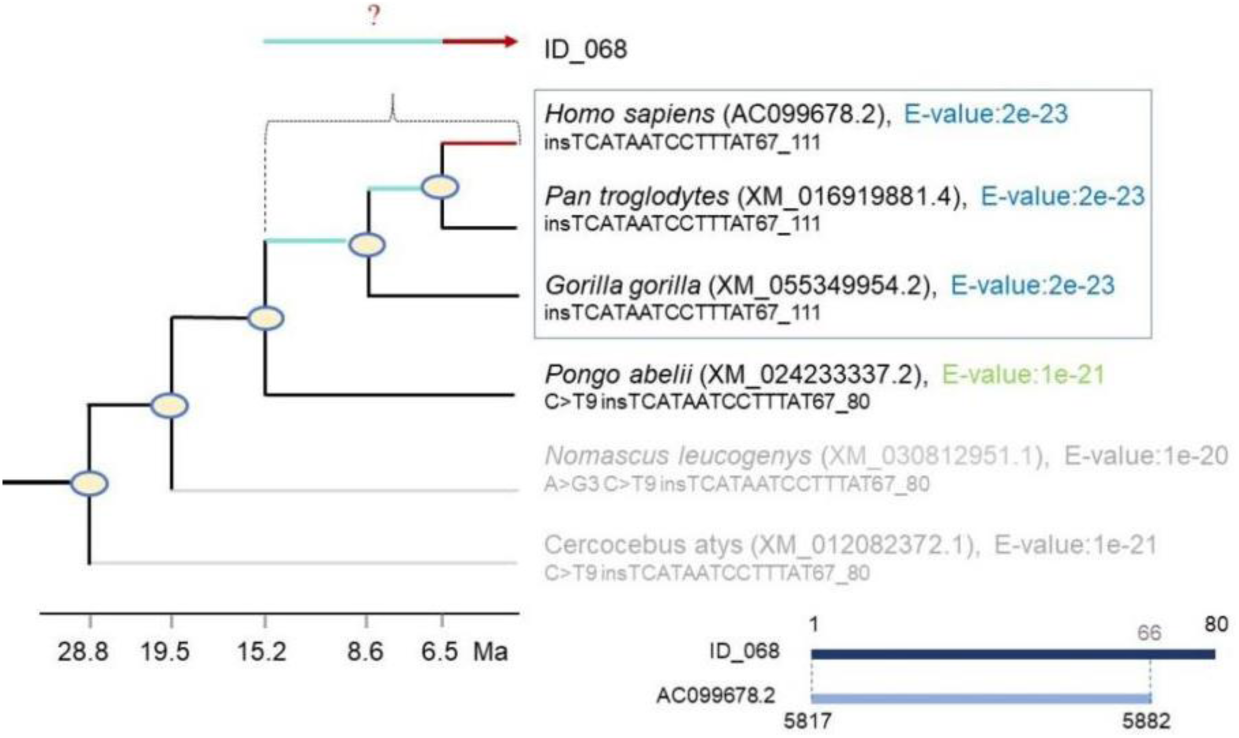
Illustration of a large Indels fragment in the ID_068 sequence, aligned w ith variations in genomic sequences across Homininae genomes. The aDNA sequence ID_068 corresponds to the genomes of *Homo sapiens*, *Pan troglodytes*, and *Gorilla gorilla*. Ellipses represent phylogenetic nodes. insTCATAATCCTTTAT67_111 indicates a short sequence insertion between the 67^th^ and 111th bases in the ID_068 DNA sequence, corresponding to the *Homo sapiens* sequence (AC099678.2).

The alignment results for the ID_180 sequence, sorted by E-value, are as follows: Homininae species (*Homo sapiens*, *Pan troglodytes*, *Gorilla gorilla*; E-value: 1e-49 to 1e-48) and *Nomascus leucogenys* (E-value: 1e-49) tie for the primary match, followed by *Cercocebus atys* (E-value: 1e-40) in second place, and *Pongo abelii* (E-value: 1e-38) in third. However, the E-values for *Pongo abelii* and *Nomascus leucogenys* misalign with their phylogenetic positions. Given that the E-value rankings of some species conflict with their evolutionary sequence, it prevents the estimation of the upper agelimit of ID_180 occurrence within the Hominoidea. ID_180 was assembled from two segments, including an antisense strand with a 5 bp overlap (Figure 2C), corresponding to the 3’-UTR of MCMDC2 mRNA. MCMDC2 plays a vital role in meiosis and DNA double-strand break repair, with mutations in mice linked to male infertility.

**Figure 2C.**
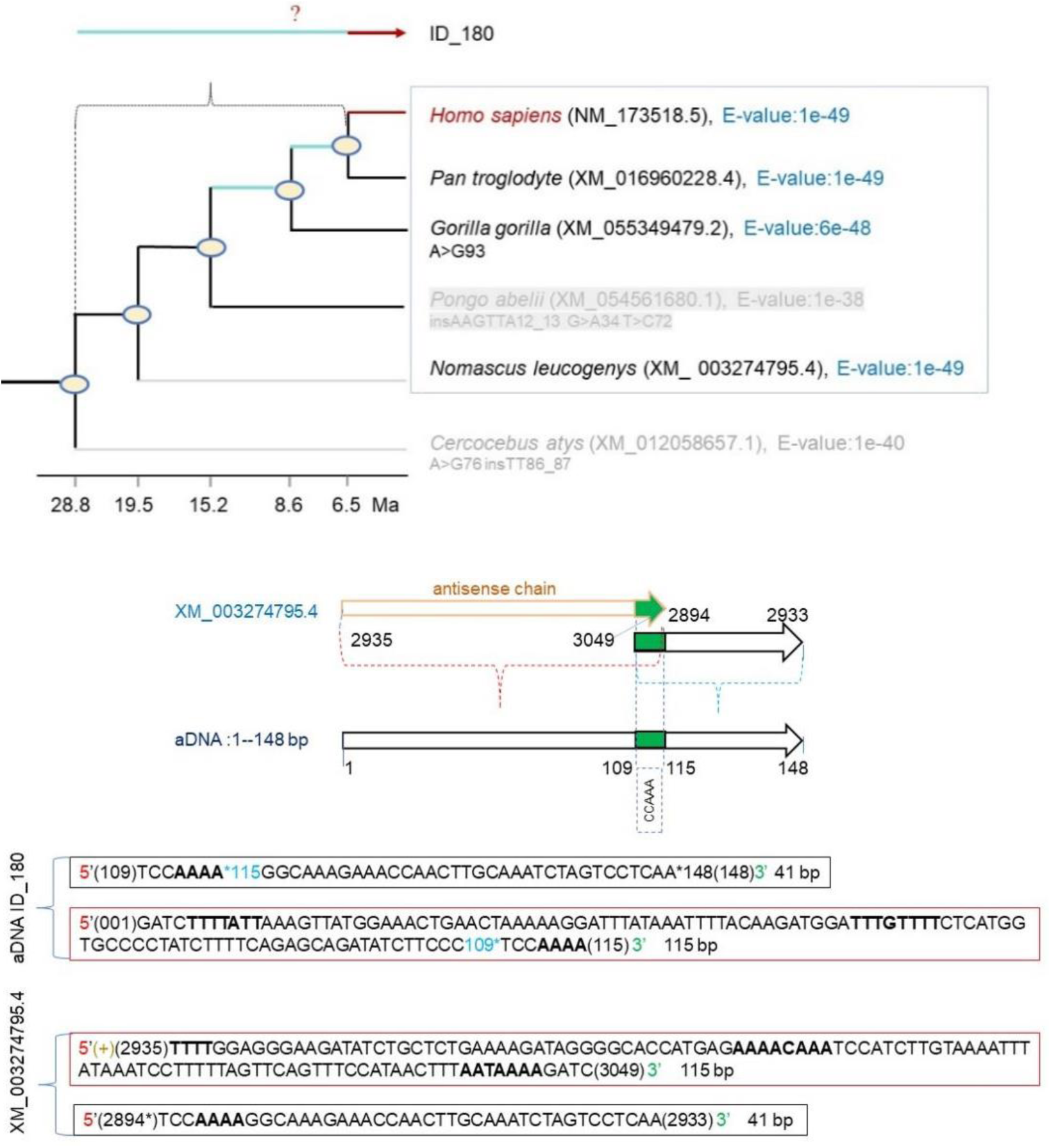
Illustration of an SRRA fragment in the aDNA sequence ID_180, depicting partial fragment inversion and sequence reassembly. Ellipses represent phylogenetic nodes. A>G93 indicates that the 93rd base “G” in the ID_180 DNA sequence corresponds to base “A” in the Gorilla genome (XM_55349479.2).

The alignment results for the ID_181 sequence, sorted by E-value, are as follows: all Homininae species (*Homo sapiens*, *Pan troglodytes*, *Gorilla gorilla*; E-value: 3e-46) and *Nomascus leucogenys* (E-value: 3e-46) tie for the primary match, followed by the non-primate European rabbit (*Oryctolagus cuniculus*; XM_070075356.1, E-value: 4e-43) in second place, with *Cercocebus atys* (E-value: 1e-42) and *Pongo abelii* (E-value: 1e-42) tying for third (Figure 2D). The E-values for *Pongo abelii* and *Oryctolagus cuniculus* deviate from their expected phylogenetic positions. Since the E-value rankings of some species conflict with their evolutionary order, we cannot estimate the upper age limit of ID_180 occurrence within the Hominoidea.

**Figure 2D.**
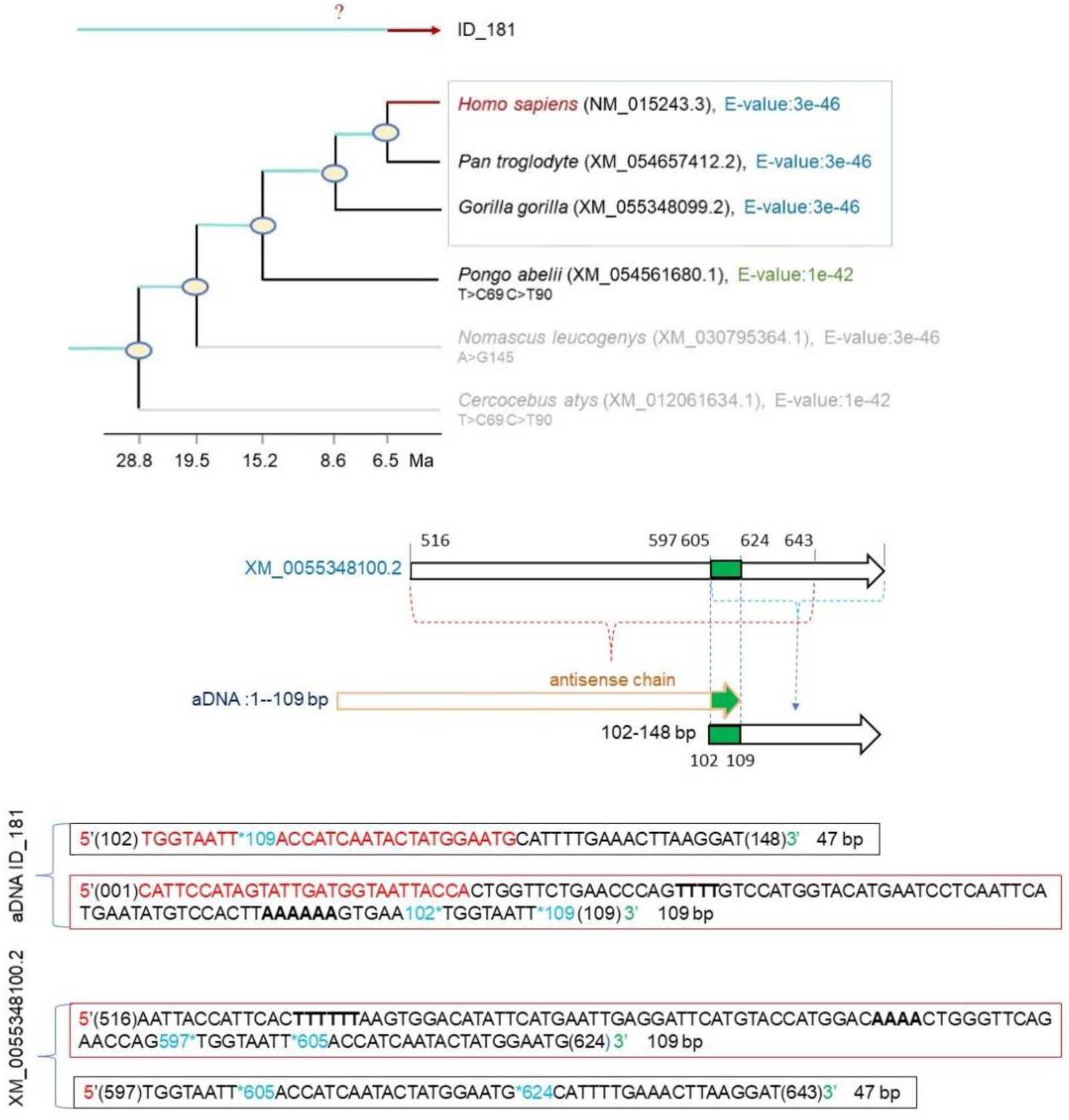
Illustration of an SRRA fragment (sequence reversal and rearrangement) in the aDNA sequence ID_181, depicting partial fragment inversion and sequence reassembly. Ellipses represent phylogenetic nodes. T>C69 indicates that the 69^th^ base “C” in the ID_181 DNA sequence corresponds to base “T” in the Pongo genome (XM_054561080.1).

This SRRA fragment lies within the CDS of VPS13B mRNA, a highly conserved region with minimal mammalian divergence. ID_181 was assembled from two segments of XM_055348100.2, including an antisense strand forming complementary strands within the sequence (marked in red, Figure 2D). This creates an insertion and hairpin structure in the CDS, potentially producing a new polypeptide or termination signal, thus altering amino acid composition and protein 3D structure. VPS13B variants may impair intracellular vesicle transport, Golgi function, and inter-organelle lipid transfer, contributing to Cohen syndrome, an autosomal recessive disorder characterized by developmental delay, intellectual disability, ocular anomalies, and related features ^3^.

**Figure 2E.**
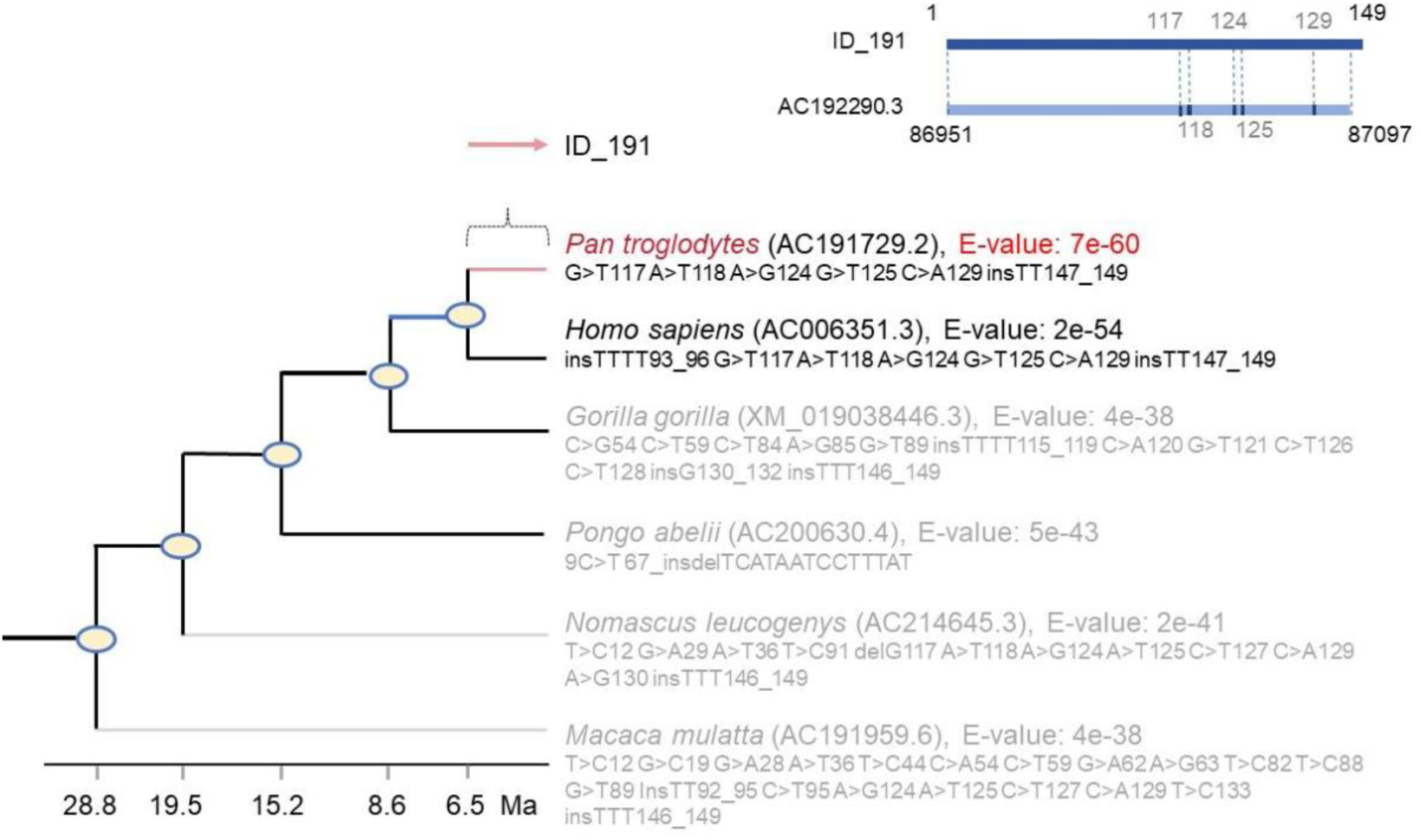
Demonstration of DNA fragments aligning closely w ith the genome of *Pan troglodytes* (species-specific). Ellipses represent phylogenetic nodes. G>T117 indicates that the 117^th^ base “T” in the ID_191 DNA sequence corresponds to base “G” in the *Pan troglodyte’s* genome (AC191729.2).

(4) SSFs: We cataloged 712 SSFs, including seven from NHPs (Table S1). The species -specificity of ID_191 aligns with *Pan troglodytes* (E-value: 7e-60), with an estimated upper age limit of 6.5 Ma for its host (Figure 2E).

2. Variant Types and Frequency: Variants include MNVs, Indels, and the most severe type, SRRAs (with a very low probability of reversion). The higher the complexity of a variant, the lower its frequency of occurrence. We found that SRRAs occur with the lowest frequency, with only three instances identified in fossils. In IDS fragments, the NHP SSF count is just seven. These sequences have an AFF < 100%, indicating they are not derived from modern humans. East Asia once hosted *Homo erectus*, but NHP evolution did not occur there. Their ancestors originated far away in Africa and the South Asian peninsula, geographically isolated from the fossil sites. Additionally, given the “old-few, new-many” characteristic of DNA in containers and the likelihood that non-human SSFs originate from the closest related species, we infer that these fragments likely derive from late Asian *Homo erectus* rather than earlier species (Table S1).

### III. DNA in the “Jar” Vessel

We excavated a Neolithic site, dated to 3,700 years ago, in Guangwu Town, Zhengzhou City, P.R. China, within the Yellow River Basin (Xia state territory).Among the artifacts was the “Jar,” a coarse pottery cooking vessel prone to cracking when dry-fired. The region’s elevation, spanning 109–253.1 meters, reduces water’s boiling point below 100°C, preserving food DNA during cooking by preventing complete degradation. The jar’s internal channels, cavities, and interstices readily trap sediment or lipids, protecting the DNA from loss or degradation caused by external factors.

1. Analysis of the “Jar” DNA

We extracted 86,908 primate-related DNA fragments from the jar’s body, spanning a temporal range from 3,700 years ago to just before excavation (Figure 3). No SSFs from NHPs were detected; all fragments originated from modern humans. Genetic variation analysis revealed fragments with MNVs, Indels, and SRRAs among the 86,908 fragments (Table S2).

**Figure 3.**
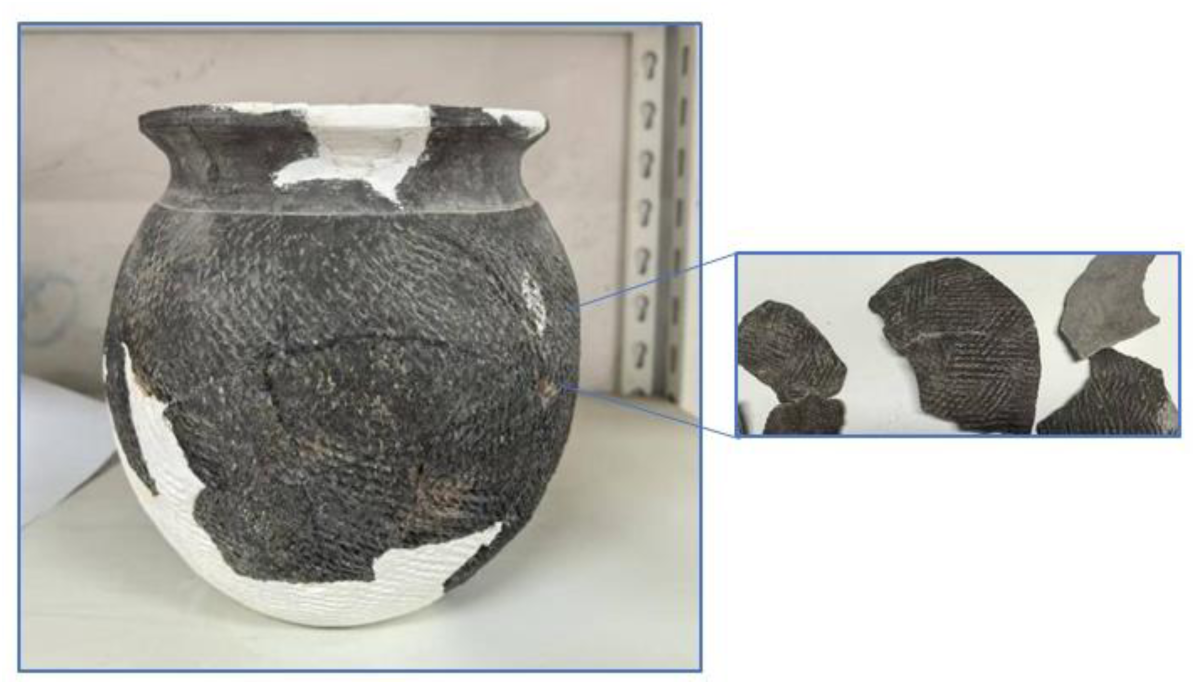
Sample H1156① w as excavated from a Neolithic (Erlitou) site in Guangw u Tow n, Zhengzhou City, China (Latitude: 34.8813597, Longitude: 113.4472668). Dated to the Xia Dynasty, ∼3,700 years ago. Photographed on October 23, 2024.

Seq_1389922 is a 148 bp sequence with an exonic structure similar to ID_180 ’s (Figure 4, indicated by red letters), corresponding to the 3’-UTR region of the modern human OLFM1 mRNA. OLFM1 is expressed in the brain, retina, and ganglion cells, and its mutation is linked to autosomal recessive disorders such as glaucoma, Alzheimer’s disease, and cancer^4^. The Seq_1389922 variant is undocumented in modern medical literature. We propose its host’s upper age limit as 8.6 Ma, tracing back to the LCA of humans and chimpanzees (*Pan troglodytes*).

1. 2. The SSFs as Evidence of Ecological and Climatic Variation

**Figure 4.**
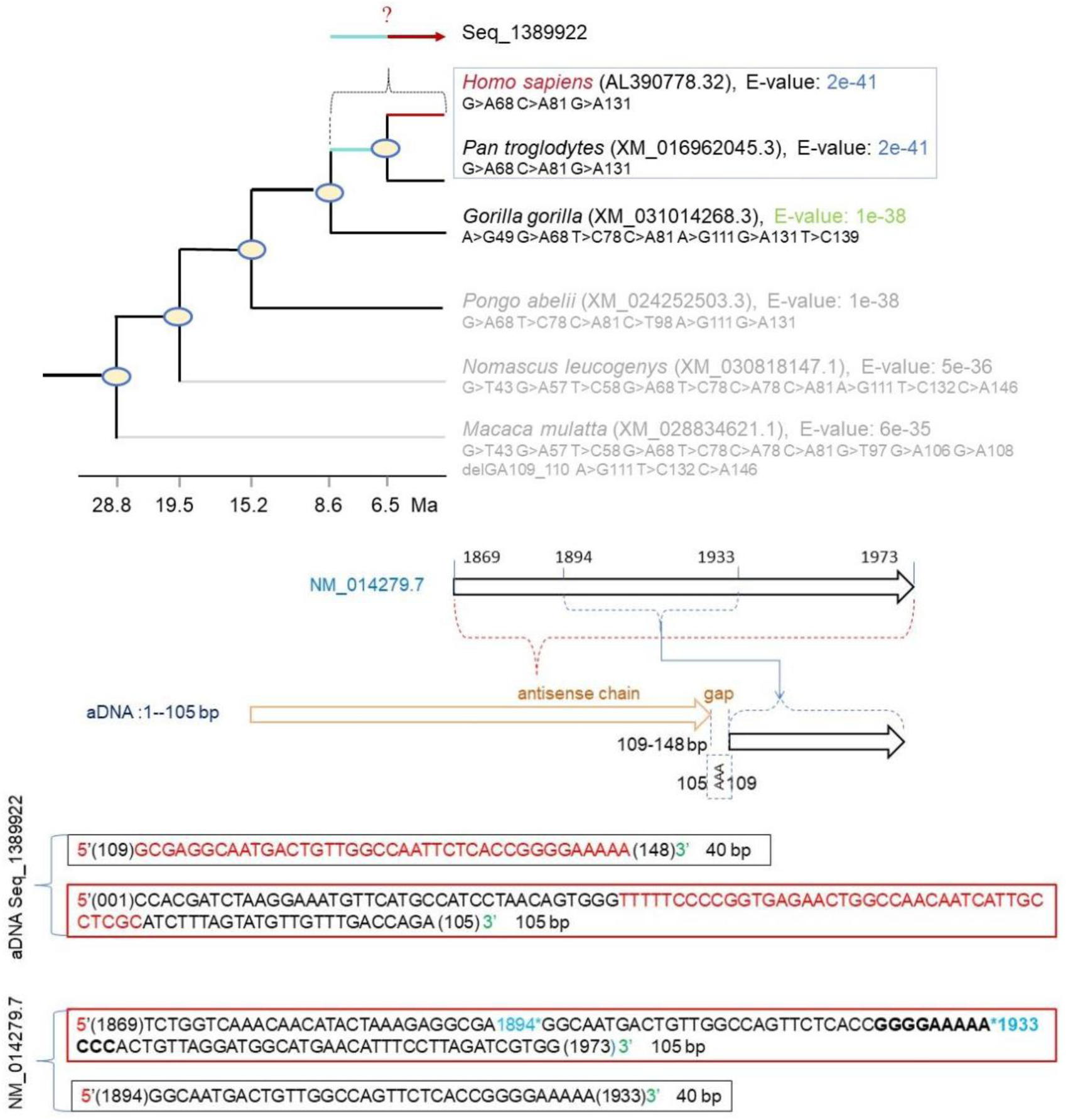
The illustration of an SRRA mutation in the aDNA sequence. Seq_1389922 depicts partial fragment inversion and sequence reassembly. Ellipses represent phylogenetic nodes. G>A68 indicates that the 68^th^ base “A” in the Seq_1389922 DNA sequence corresponds to base “G” in the *Homo sapiens* genome (AL390778.32).

We identified one SSF corresponding to the extinct *Equus quagga* (Table 2), a species with no prior record of existence in the local region^5^. Zebras form a monophyletic group and do not share this ancestor with horses or donkeys. Consequently, this fragment is unlikely to have originated from hybridization with local equid species^6^. Two further SSFs align with the tropical oil palm (*Elaeis guineensis*), believed to have originated in West and Central Africa, with a natural range limited to Africa and cultivation centered in Southeast Asia, never native to this locality. Guangwu Town, located along the Yellow River’s southern bank in a water-rich network, borders the Songshan mountain range to the west. Archaeological evidence indicates a warm, humid climate 3,700 years ago, typical of tropical conditions ^7^.

### IV. Molecular Characteristics and Biological Significance of SRRA

1. **The SRRA** mutation is exceedingly rare and has neither been a focus of attention nor evaluated for its functional value or evolutionarysignificance byscientists. SRRAs predominantly occur within exonic sequences encoding mRNA. In the sense strand, a segment of the antisense strand emerges and is inserted, in the 5’→3’ direction, upstream of a complementary segment of the sense strand. This insertion occurs immediately adjacent to or separated by a sequence interval from the complem entary region, accompanied by MNVs and Indels, significantly altering the original gene sequence.
2. **Possible Formation Mechanisms of SRRA:** SRRA is a complex gene variant, and its formation mechanism likely involves a process in which mRNA is reverse -transcribed into DNAand inserted into the genome, resembling retro-transposition mediated by LINE elements (long interspersed nuclear elements, LINEs). However, given the numerous unresolved details in SRRA’s formation process, this mechanism cannot fully account for SRRA’s formation. Other genomic variation mechanisms also fail to offer a satisfactory explanation, including replication slippage-recombination, non-homologous end joining (NHEJ), DNA double-strand break repair, and genomic rearrangements (e.g., translocations, inversions, and other large-scale genomic changes).
3. **Distribution of SRRAs in Primate Genomes:** Three SRRA fragments were identified in fossils, with the following distribution: ID_179, ID_180, and ID_181 map to exonic sequences of primate mRNA-encoding genes. The two SRRA fragments in the “Jar” also map to exonic sequences of primate mRNA-encoding genes (Figure 5).
4. **Role of SRRA in Adaptive Evolution:** SRRAs occur within exonic sequences of mRNA-encoding genes and can influence post-transcriptional regulation of mRNA in the following ways. (1) Altering mRNA translation efficiencies, such as through hairpin structures formed in the 3’-UTR or 5’-UTR with Indels (e.g., Seq_1389922). (2) Directly affecting protein expression profiles, as exemplified by ID_181, which forms an insertion sequence within the coding sequence (CDS) of VPS13B, potentially generating hairpin structures, additional coding sequences, or stop codons. (3) During evolution, SRRAs may separate from the exons they reside within, ceasing to appear in mRNA sequences. For example, four sequences with SRRA-like hairpin structures internally (SRRA-related: ID_174, ID_175, ID_177, ID_178) map to non-coding mRNA sequences in *Homo sapiens* and *Pan troglodytes*, with no corresponding regions in other primates. We hypothesize that these represent “SRRA relics ” retained in the human and chimpanzee genomes, whereby the genomes have repurposed SRRA-associated exonic sequences into stable non-coding mRNAsequences; this specific process requires further investigation (Figure 5).

**Furthermore**, the SRRA variation regulatory mechanism shares similarities with the transposase generation mechanism, “coding region sliding replication and recombination ”, recently identified in the *Lycoptera* genome^1^. Both processes utilize the host gene sequence as a template, increasing their total number of bases, sequence diversity, and genomic complexity without introducing external DNA sequences. Elucidating these mechanisms could contribute a new chapter to molecular genetics.

**Figure 5.**
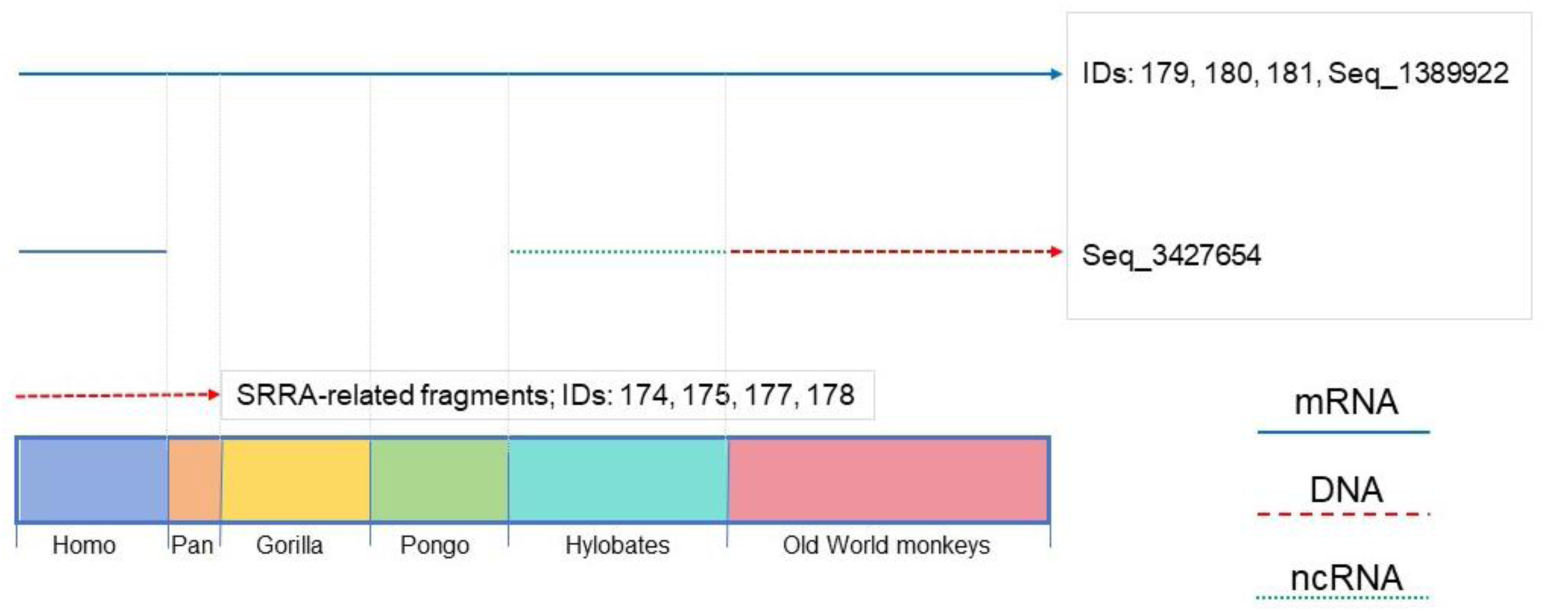
Distribution of SRRA-Associated Sequences in Primate Genomes.

## Discussion

### I. Possible Origins of Primate SSFs

Within the Jehol Biota *Lycoptera* fossils, we identified NHP SSFs, including three from *Pan troglodytes* (plus one highly likely sequence), one from *Pongo abelii* (with two highly likely sequences), one from *Hylobates moloch* (with two highly likely sequences), and two from *Macaca mulatta*. These sequences diverge significantly from modern genomes (AFF < 100%), confirming they do not stem from extant species. The habitats of these living primate species are geographically remote from the Jehol region, ensuring isolation that precludes genomic contamination from modern populations. We propose that these NHP SSFs originated from extinct late Asian *Homo erectus*, whose prolonged habitation in the region allowed their DNAto enter the environment and become preserved in the fossils.

### II. Our Reasoning

East Asia is not a hub of NHP evolution; modern chimpanzees (*Pan troglodytes*) and their ancestors emerged in Africa, as did the Hominidae LCA, with the arrival of its descendants in East Asia still undated. Our methodology offers technical support for locating Asian *Homo erectus* fossils and reconstructing their migration paths. Human evolution followed a linear trajectory from African *Homo erectus* through *Homo heidelbergensis* to modern humans. Asian *Homo erectus*, spanning approximately 2 million to 100,000 years ago, dispersed from Africa to Asia (e.g., Peking Man, Java Man), retaining conservative skeletal traits—such as thick skulls and low foreheads—akin to early African counterparts. Conventionally, their extinction is attributed to geographic isolation and limited gene flow, marking them as an “evolutionary dead end”^8–11^. Evolution hinges on DNA variation, and the conservatism of Asian *Homo erectus* may reflect a genome shaped predominantly by genetic drift. Modern human chromosome 2 resulted from the fusion of ancestral chromosomes 2Aand 2B, homologous to those in chimpanzees. We hypothesize that Asian *Homo erectus* maintained 24 chromosome pairs (like chimpanzees), potentially fostering reproductive isolation from incoming modern populations. Future detection of “non-fusion evidence” in DNA from their remains or artifacts, linked to chromosomes 2A and 2B, could verify this karyotype. Although some studies suggest minor gene flow with Denisovans^12, 13^, we challenge these claims and the reliability of their methods ^14^.

### III. Characteristics of IDS and Their Role in Primate Divergence

IDS variations are ubiquitous in modern humans (e.g., contemporary individuals from Guangwu) and ancient human populations. This raises an intriguing question: Why do many IDS, particularly SRRAs, observed in aDNA appear absent from modern genomes? Two potential pathways may explain this phenomenon: host extinction induced by SRRA or genomic correction. First, Seq_1389922_OLFM1 is associated with an autosomal recessive genetic disorder, leading to increased mortality and reduced family size compared to the general population—a cost of SRRA mutations. Second, during a phase of rapid primate evolution, SRRA may have provided value by offering multiple options for mRNA translation and protein patterns without completely disrupting the function of the affected gene (e.g., SRRA can occur in the mRNA 3’-UTR). At this stage, the organism could still tolerate the changes induced by SRRA. However, once this phase concluded and the genome underwent remodeling (including genetic changes beyond SRRA mutations), the functional role of SRRA-containing regions became redundant, leading to their eventual removal from exonic sequences. Thus, SRRA likely serves as an intermediate molecular mediator of rapid genomic evolution. Additionally, studies suggest gene deletion may drive evolution by simplifying the genome, conferring a selective advantage during adaptation to new environments or stressors^15, 16^. Whether SRRA can induce wholesale gene deletion remains unknown and worthy of further investigation.

### IV. Contrasting Effects of MNV and Indel Mutations with SRRA

Compared to the SRRA, the mechanisms of MNV and Indel mutations are considerably more direct. When occurring within CDS, these mutations can have pronounced effects, potentially leading to functional loss, no discernible change, or functional enhancement i n very few cases. For instance, *Anab arilius grahami,* an endemic fish species of the Xenocyprididae family, is found exclusively in Fu Xian Lake, located in southeastern China. This species is characterized by a small bodysize (10–15 cm) and is geographically isolated from the *Lycoptera* fossil site. Temporally, it is separated from the fossils by 120 Ma. Comparative analysis of the Growth arrest–specific 2 protein (GAS2) sequences reveals that, uniquely among known fish species, the GAS2 protein in *A. grahami* contains an additional segment derived from the CDS of a transposase (Tcb2, positions 114–194, ROL47475.1). This indicates that a segment of the Tcb2 CDS was inserted upstream of the C-terminal region of the CDS. These findings suggest two key implications: first, gene flow likely occurred between *Lycoptera* and the ancestral lineage of Xenocyprididae; second, this CDS Insertion was retained by the host genome, probably enhancing GAS2 function (Fig. 3 in Reference 1). In contrast, SRRA represents a newly identified mutation type. Its ability to generate diverse mRNA patterns appears to provide the genome with a “buffering” period—even within exonic CDS—potentially mitigating the immediate impacts of genetic changes. Consequently, we hypothesize that SRRA may exert diverse effects on host function, phenotype, morphology, and adaptive evolution. However, these potential impacts require further investigation to be fully elucidated.

### V. Genetic Information Loss in Fossil Restoration

Fossil restoration typically employs fine tools to remove sediment layers, deposits, and impurities from bone surfaces and contours, aiming to reconstruct the original skeletal shape. However, soft tissues—including muscles, organs, and skin—are replaced by mineral deposits, often yellow or black, formed through processes such as the Maillard reaction or interactions between organic acids from the organism and environmental heavy metal salts, yielding colored precipitates. These deposits, frequently found between bones, are rich in oriDNA. Regrettably, conventional restoration techniques discard these materials as waste, squandering valuable scientific resources. Our method enables the extraction of aDNA from such deposits. We urge researchers to preserve these materials during restoration to safeguard critical genetic information.

### VI. Critique of Traditional Molecular Paleontology Method

**We propose the novel concept** of a “DNA container”: since the burial of skeletal fossils in soil, they not only carry oriDNA but also continuously adsorb eDNA, including genetic material from extinct humans, modern humans, and their associated species, which is retained internally. The fossil skeletons of Neanderthals and Denisovans also function as “DNAcontainers ”, containing oriDNAthat diminishes over time and accumulates eDNA, much of which is of ancient origin. These findings directly challenge the initial assumptions of the traditional method established by Dr. Pääbo and his team in the early 2000s.

**Fundamental Logic of the Traditional Method**: (1) Initial assumption: aDNA in skeletal fossils originates solely from the corresponding individual, i.e., oriDNA. (2) Contamination control: Fossil DNAis mixed with modern contamination DNA(from excavation and handling), which can be removed through strict anti-contamination measures and high-grade sterile laboratories. (3) Screening criteria: DNAmust exhibit nucleic acid degradation, with deamination as a prerequisite. Fragments showing deamination in the terminal 25% of bases at the 3’ end are identified as oriDNA, excluding modern DNA contamination. (4) Genome construction: Screened fragments (typically 40–60 bp) are aligned via BLAST against a predefined target genome (e.g., human or great ape) and assembled to construct an ancient human genome.

**Fundamental Flaws of the Traditional Method:** (1) Neglect of exogenous aDNA: the aDNA of skeletal fossils is not limited to oriDNA but includes substantial environmental aDNA (e.g., from ancient modern humans, animals, plants, and microbes) and its “old-few, new-many” pattern. The traditional method fails to account for this complexity. (2) Limitations in removing contaminant DNA: Even under the strictest sterile conditions, exogenous aDNA sealed within the “DNA container” cannot be removed, inevitably mixing non-target genetic material into the sample (Figure S2). (3) Misconceptions in using deamination to screen aDNA: Deamination occurs not only in skeletal DNA but also in environmental aDNA, leading to the erroneous selection of non-oriDNA; meanwhile, non-deaminated fragments (e.g., those preserved by minerals or lipids), which carry more complete information, are overlooked, such as SSF and SRRA sequences in this study. (4) Unreliability of alignment methods: Using a predefined genome as a BLAST reference, a large amount of exogenous aDNA (e.g., soil microbial DNA) is misclassified into the ancient human genome. This misclassification is particularly pronounced with short fragments (40–60 bp) and high E-values. We analyzed the only publicly available sequence from their paper (via online BLAST with the nr/nt database) and found it matches multiple kingdoms, including soil microbes ^12, 14^, highlighting the methodological flaws. (5) Negative Control: We performed BLAST alignments of DNA fragments from the GAT genomes (derived from petroleum and *Lycoptera* fossils) using a reference genome limited to the primate mitogenome database. Under these conditions, a substantial number of fragments matched the Neanderthal genome. However, the petroleum and fossil sites are not regions historically inhabited by Neanderthals. This indicates that selecting an appropriate BLAST reference database is critical, and artificially restricting the dataset can lead to misleading results.

**Conclusion:** The traditional method has distorted a wealth of research data generated by this approach, leading to erroneous conclusions. Over the past decade, studies with these flawed conclusions have been widely published in top-tier journals. Regrettably, this represents not a triumph of scientific progress but a predetermined academic catastrophe.

### VI. Biological Significance of This Study

**Molecular Paleontology and Study Objectives**: Molecular paleontology aims to analyze ancient aDNA to elucidate species relationships, substantiate evolutionary theories, and illuminate the history, mechanisms, and future trajectories of life’s evolution. A central obstacle in this field is the lack of an effective and reliable method for extracting and analyzing aDNA, particularly from samples of abundant lithic fossils and non-skeletal materials, such as crude pottery, petroleum, and sedimentary rock s. This study tackles this challenge by developing innovative methods, rectifying prevailing misconceptions, and enhancing the efficacy and scope of aDNA analysis.

**Implications of the Findings:** The results of this study challenge conventional perspectives and undermine the foundational assumptions of the traditional method, thereby destabilizing the established knowledge framework. This necessitates a critical reevaluation of numerous conclusions, potentially jeopardizing the integrity of the entire framework. For example, interpretations concerning the evolution of late *Homo sapiens*, gene flow between extinct hominins and modern humans, and the origins of disease susceptibility genes—derived from the genomic analyses of Neanderthal and Denisovan data—may contain fundamental inaccuracies. These findings expose a sobering reality: most natural fossils cannot be fully isolated from eDNA, rendering it impossible to exclude the possibility of ancient contamination. As a result, such fossils are deemed unsuitable for reliable genomic reconstruction. Consequently, the knowledge framework of molecular paleontology requires substantial revision, necessitating the development of new theoretical paradigms and methodological approaches.

**Contribution to Evolutionary Theory:** This study advances evolutionary theory by establishing a method to estimate the upper temporal boundary of the origin of IDS, enabling the examination of the temporal relationship between the emergence of IDS and the evolution of primate species. Furthermore, it identifies two distinct subclasses of IDS: SSF and SRRA. SSF facilitates tracking extinct species, while SRRA contributes to a multifaceted post-transcriptional regulatory mechanism. This mechanism enhances the genome’s capacity to adapt rapidly to environmental changes, potentially functioning as an evolutionary “gene switch”. Following the remodeling of the host genome, SRRA may be excised from exons. By elucidating these processes at the molecular level, this study partially resolves Darwin’s conundrum regarding the paucity of transitional species in the fossil record.

**Summary and Outlook:** A major challenge in primate aDNA study is the scarcity of fossils, exacerbated by contamination with exogenous DNA, including that from modern humans. Our approach, the most robust solution to date, is grounded in a foundational logic that prevents the conflation of DNA from diverse sources. This method extends its utility by extracting aDNA from non-skeletal materials and screening for IDS, including SRRAs and SSFs, offering an effective tool for investigating primate molecular evolution. It also underscores the possibility and urgency of extracting aDNA from an integrated trinity of fossils, source rocks, and petroleum to establish a global aDNA database. By classifying species within this database, we can integrate factors such as fragment age limits (evolutionary timelines), host survival duration (stratigraphic context), species distribution patterns (including extinct taxa), paleoclimatic and geographic shifts, paleo-ecological changes, and unique sequence characteristics. Using enhanced algorithms and artificial intelligence, this approach elucidates molecular mechanisms driving pri mate evolution or extinction, particularly sequence alterations under environmental stress, revealing tangible “gene switch mechanisms”. These findings reference humanity’s future adaptation to environmental changes and survival in space.

Future applications of this method encompass leveraging aDNA to inform the discovery of morphological fossils in natural environments; extracting aDNA from fossils, sedimentary rocks, and petroleum to construct novel aDNA databases; and fostering interdisciplinary advancements by facilitating the integration of molecular paleontology, archaeology, and geology. Since the unsuccessful attempts to purify dinosaur DNA in the 1990s^17^, humanity has relentlessly pursued the reconstruction of extinct species ’ genomes and the revival of ancient life. We propose a critical concept for this investigation: the “ideal fossil.” Such remains should originate from a live burial, encapsulated in materials that isolate environmental macromolecules, and undergo rapid dehydration, salting (with mineral salt ions permeating tissues to preserve nucleic acids), and moderate heating (e.g., 55–100°C) to inactivate microorganisms and prevent DNA degradation while minimizing bacterial proliferation. Grouped SSFs serve as effective biomarkers for monitoring the origins and abundance of environmental molecules. A genomic assembly can be initiated once the level of eDNA is deemed acceptable. The concepts and methodologies introduced in this study establish a robust framework to support this objective.

## Materials and Methods

### I. Wet Laboratory Experimental Procedures

1. Processing of *Lycoptera* Fossil Material: Conducted as described in reference 1.
2. **2. Processing of “Jar” Material:** Pottery fragments (total weight 460 g, consisting of smaller pieces) were selected, rinsed repeatedly with purified water, and air-dried; this step was repeated once. Sample surfaces were wiped twice with a wet cloth, followed by a single wipe with an alcohol -soaked paper towel. Further processing followed reference 1.
3. **3. Evaluation of Internal Volume Ratio**: Materials were cleaned, dried at 56°C for 72 hours, and weighed (W1). Dried materials were immersed in purified water, subjected to ultrasonic treatment (10 W, 2 hours), then quickly wiped to remove surface water and weighed again (W2). Wet materials were placed in a graduated container and submerged in water, and the total volume (V2) was measured. Internal volume (V1) was calculated as follows: 

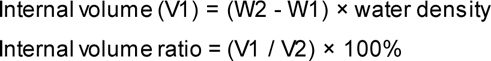
4. **DNA Extraction, Library Construction, and Sequencing:** Detailed in reference 1.

### II. Dry Laboratory Experimental Procedures

**1. AFF**: We defined AFF to quantify the similarity between a sequence and its matched genome. The calculation involves multiplying the Identity and Cover values obtained from NCBI BLAST and converting the result to a percentage (AFF = Identity × Cover × 100%). aDNA typically exhibits significant sequence divergence from modern genomes. A high AFF value indicates a high similarity to modern genomes, whereas a low value suggests significant divergence.
**2. SSF Identification**

**Rationale**: DNA sequences exhibit widespread similarity across species, from prokaryotes to higher primates. For example, humans and chimpanzees share over 98% sequence identity in most fragments, while similarity with bacteria can exceed 20%. To detect human DNA contamination in a chimpanzee’s sample, the mega screen method is applied to assess specificity^1^. For a given fragment, E-values are calculated against the chimpanzee and human genomes and compared, when the difference between the E-value of the former and the latter spans two orders of magnitude (specifically, when the quotient of the former E-value divided by the latter E-value is ≤ 0.01, expressed in scientific notation as ≤ 1e-02), indicating a significant disparity, the sequence is identified as chimpanzees -derived and termed a chimpanzees SSF. SSFs constitute a minimal fraction of the genome, and their detection in a s ample is sufficient to infer the presence of the species.

#### Specific Procedures

1. Determining BLAST Result Confidence Threshold Using the Best E-value Mode Rationale: First, the GAT genome comprises eDNA fragments from diverse sources, precluding the use of predefined reference genomes; thus, we utilized the NCBI nucleotide sequence database (e.g., Version 5), encompassing all species. Second, BLAST analysis of the GAT genome risks misaligning sequences due to the variety of environmental species involved, particularly with short fragments (50–60 bp). Hence, a threshold (E-value ≤ 1e-07) is set to exclude such ones.
2. Comparison of Taxonomic Units Exhibiting Significant E-value Disparities Using the MS Mode Initially, we establish taxonomic “units” for species classification, such as “genus” or “family,” to serve as the basis for comparison. The first species represents all species within its designated taxonomic unit (e.g., the same genus or family), whereas the second species represents all species with in a distinct second taxonomic unit. In the online BLAST analysis, the species with the highest E-value in the first taxonomic unit was selected, followed by the selection of the species with the lowest E-value in the second taxonomic unit. Second, significant differences are assessed: if the quotient of the former E-value divided by the latter E-value is ≤ 0.01 (1e-02), the fragment is deemed specific to the taxonomic unit of the first species, confirming its “specificity”; if the quotient distance ranges from 0.1 to 0.01 (1e-02 ~1e-01) specificity is highly likely. The taxonomic units represented by the first and second species may differ in level; for example, the first species may represent the Homininae, while the second represents the genus *Pongo*.
3. Ensuring Kinship Similarity in Top Three Species Rationale: “Kinship similarity” in DNA sequences is a hallmark of molecular evolution. In this study, the top three species ranked by E-value should belong to Primates. If the first is *Homo sapiens*, the second should be a non-human ape or at least a primate. If the second is bacterial and the first’s E-value exceeds 1e-15, the sequence lacks “specificity.”

**Note:** E-values should be obtained from BLAST results under identical conditions (parameters).

### 3. Estimating the Upper Age Limit of Sequence

**Rationale**: The DNA fragment undergoes BLAST analysis, and matched species are ranked according to their respective E-values. These species are then integrated into a phylogenetic tree. If the evolutionary hierarchy of the observed species remains consistent, the node connecting the primary and secondary species indicates the upper age limit for the sequence.

**Specific Procedures:**

Phylogenetic tree nodes were employed as unit boundaries, and the E-values of species within the closest unit were compared. If the E-values within a given node are identical or their quotient (the former E-value divided by the latter) differs by more than 1e-02, the species within that node are classified as co-ranked first. Subsequently, comparisons are extended to species in the adjacent node. Should the quotient be less than 1e-02, co-ranking is excluded, and the species from the next node is designated as the second species. Specifically, when the quotient of the E-values between the first and second species is less than 1e-02, the sequence is inferred to originate from the first species. By integrating this approach with phylogenetic orders, the upper temporal limit of the sequence ’s (and thus the host’s) emergence can be estimated.

**Note**: Discrepancies may arise between traditional morphological and molecular classifications. For instance, if the E-value rank of sequence-matched species in the constructed phylogenetic tree conflicts with the inferred evolutionary relationship, a reliable upper age limit cannot be accurately estimated.

### 4. Software Applications

Sequence alignment is conducted using the NCBI Nucleotide BLAST online tool, starting with the Core_nt database and transitioning to the nr/nt database depending on the i nitial results. DNAsequence homologyanalysis is conducted with MEGA11 software.

---

**Figure S1.**
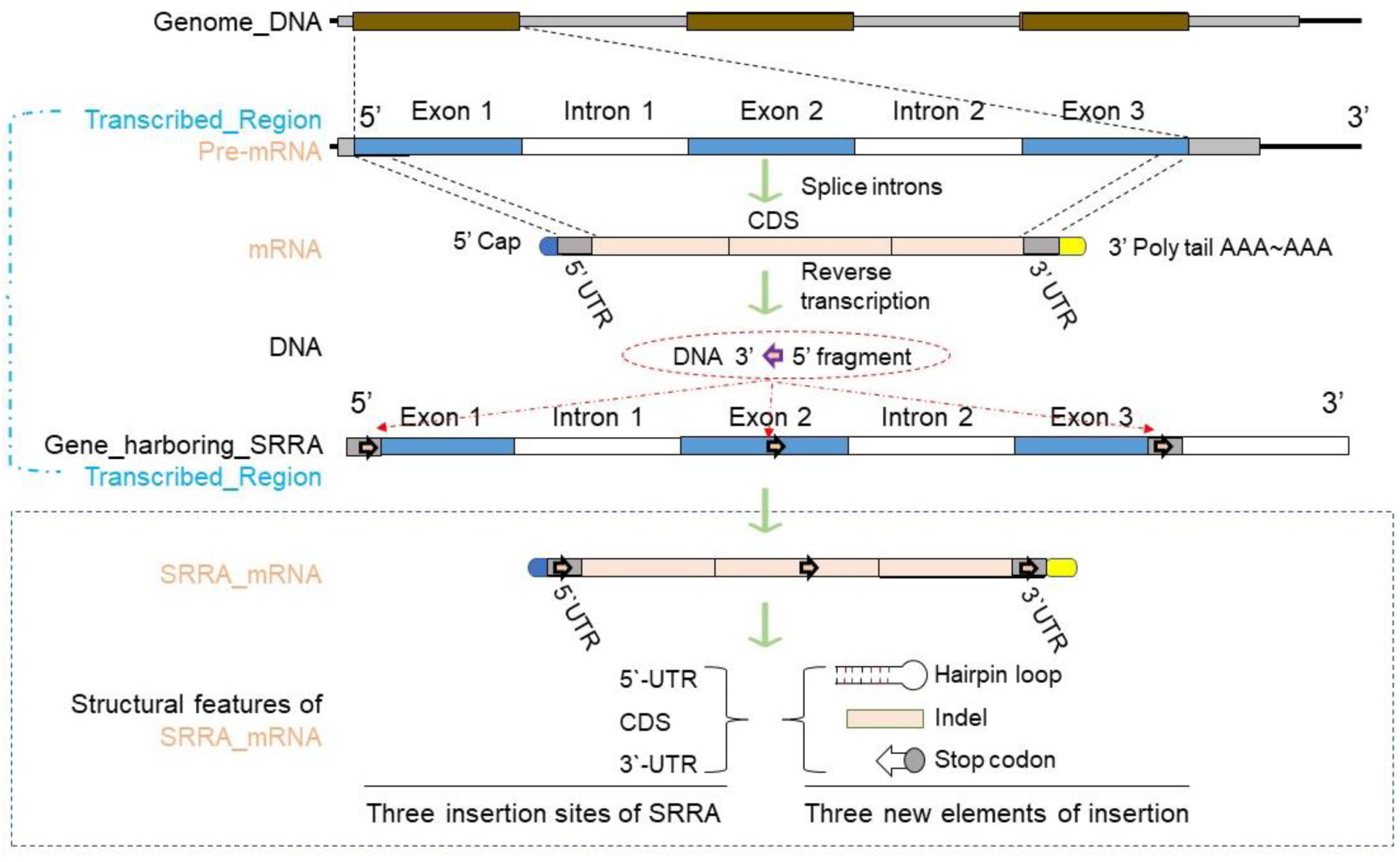
Formation of SRRAs and Their Effects on m RNA Molecular Structures

**Figure S2.**
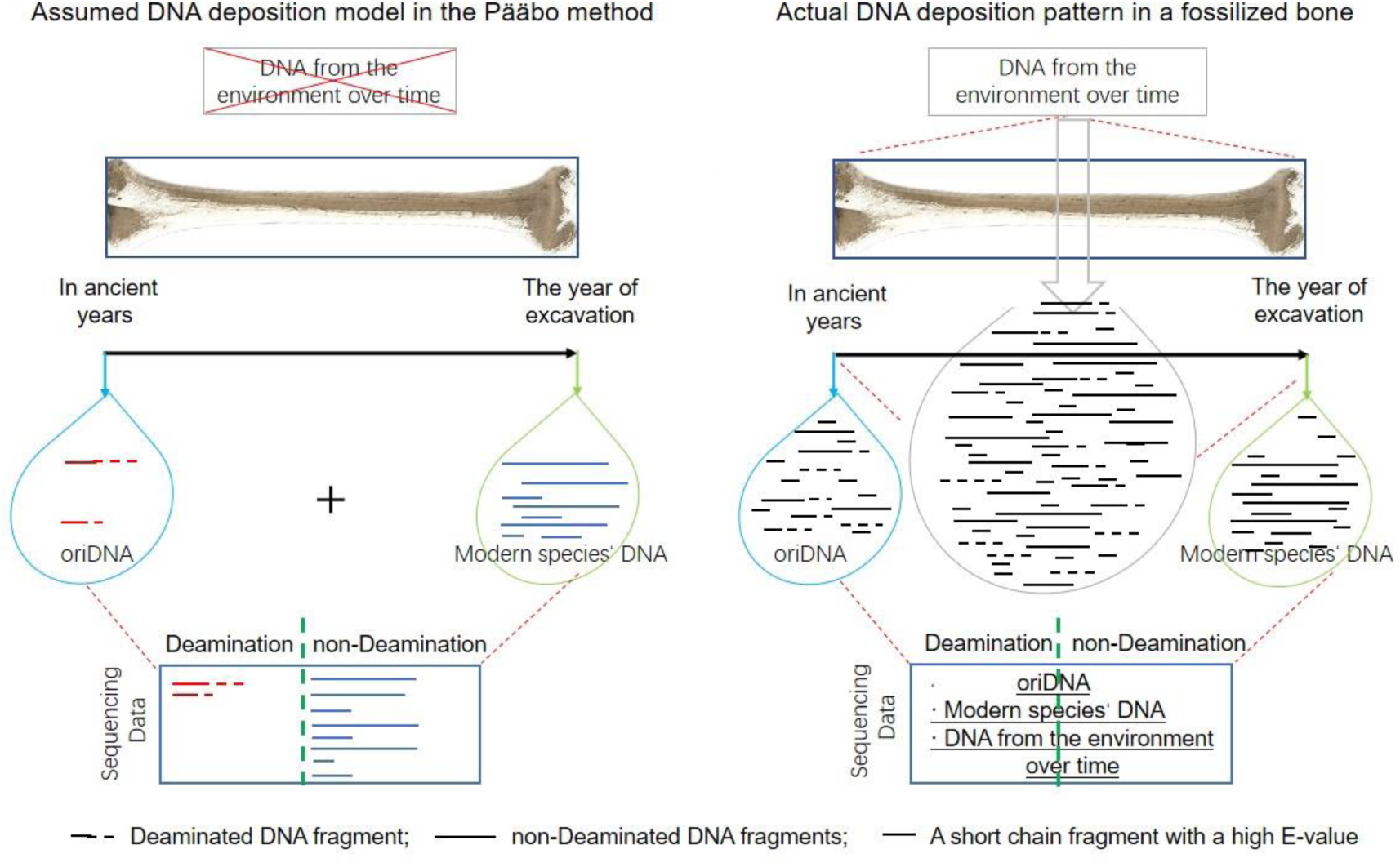
This figure illustrates the traditional approach, w hich posits that a skeletal fossil primarily consists of oriDNA before excavation, with modern macromolecules added during the excavation (left panel). This study shows that eDNA fragments can infiltrate and accumulate w ithin bones, from burial until excavation, ultimately becoming a significant component of the fossil (right panel).

**Table S1.**
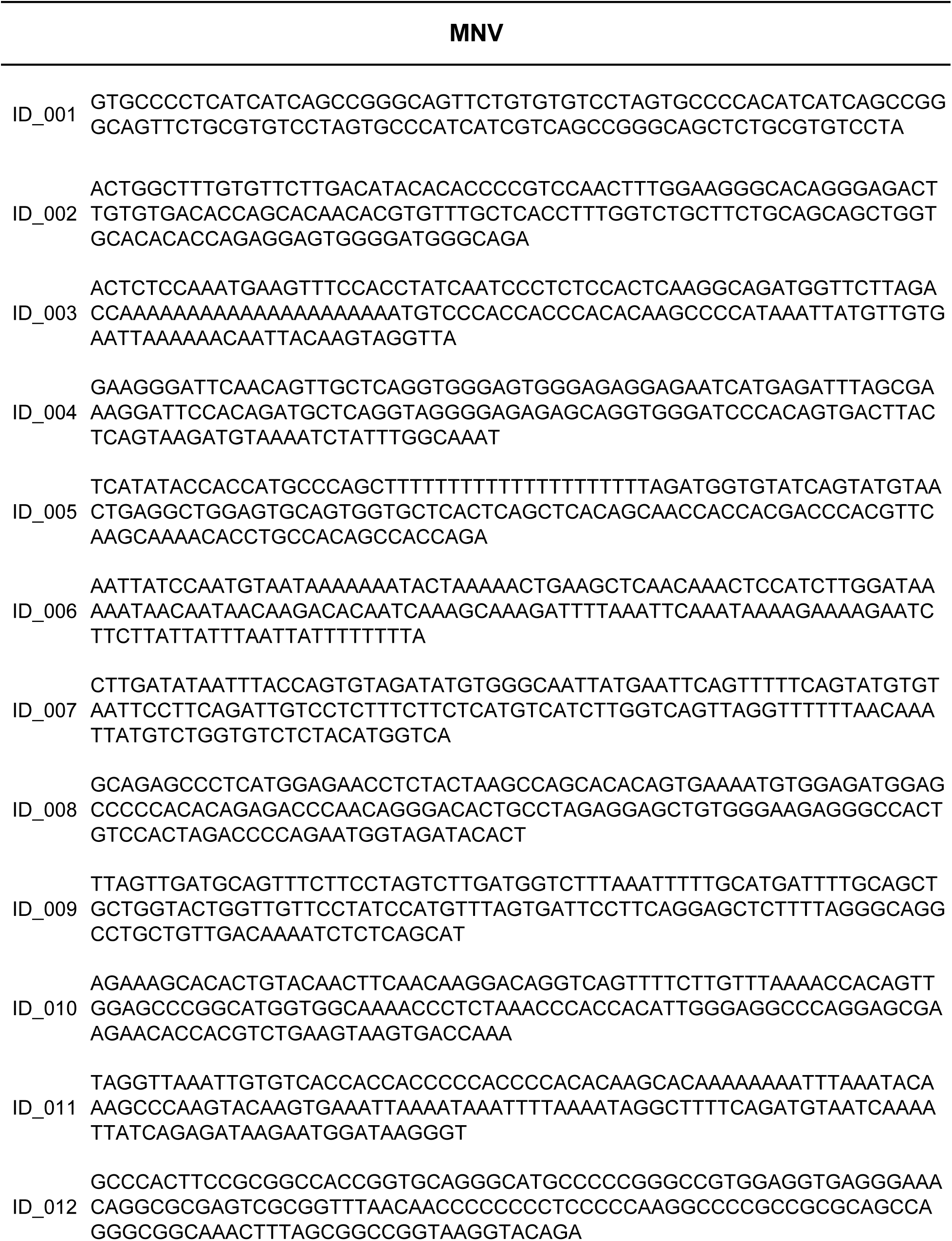
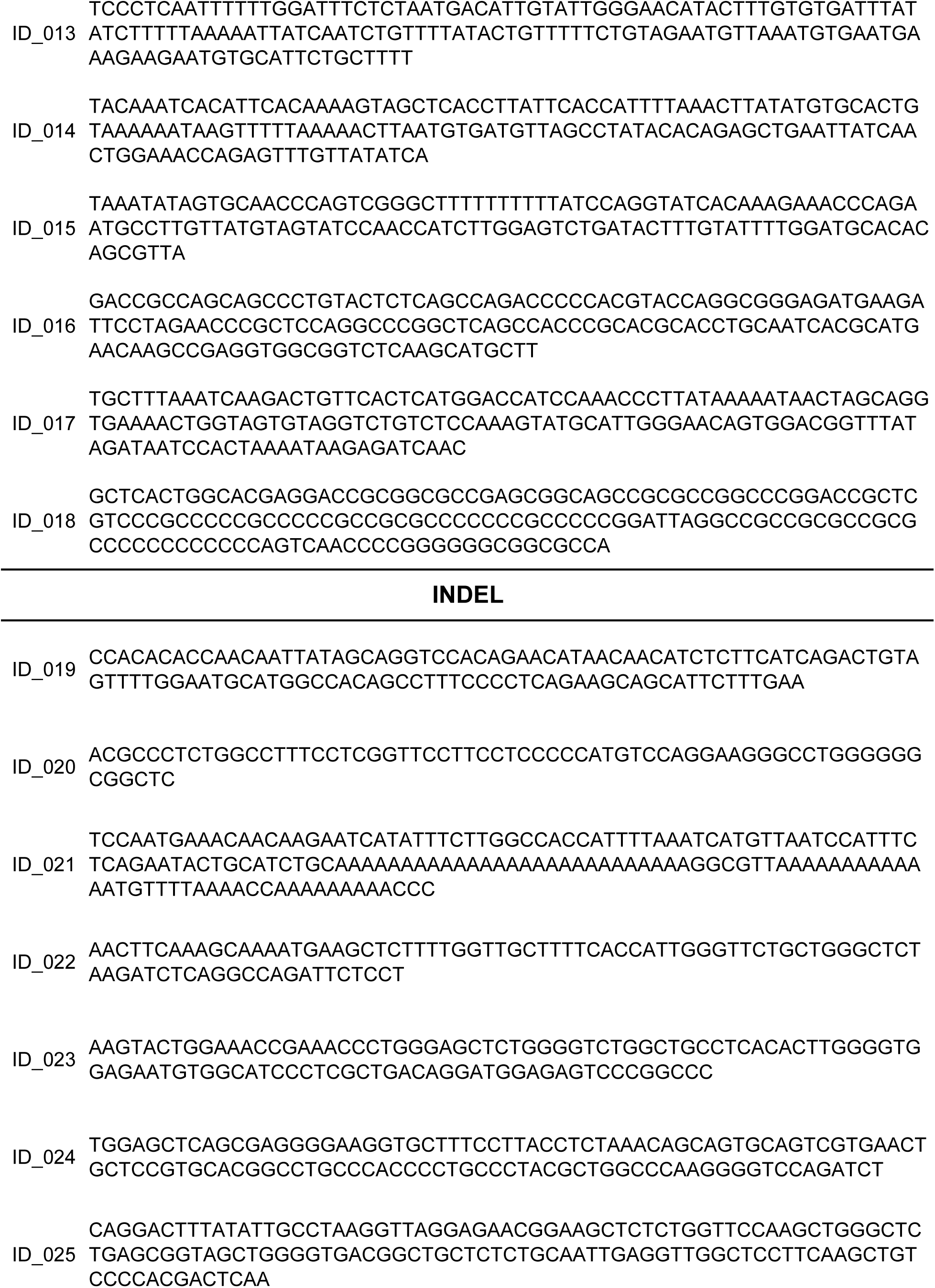
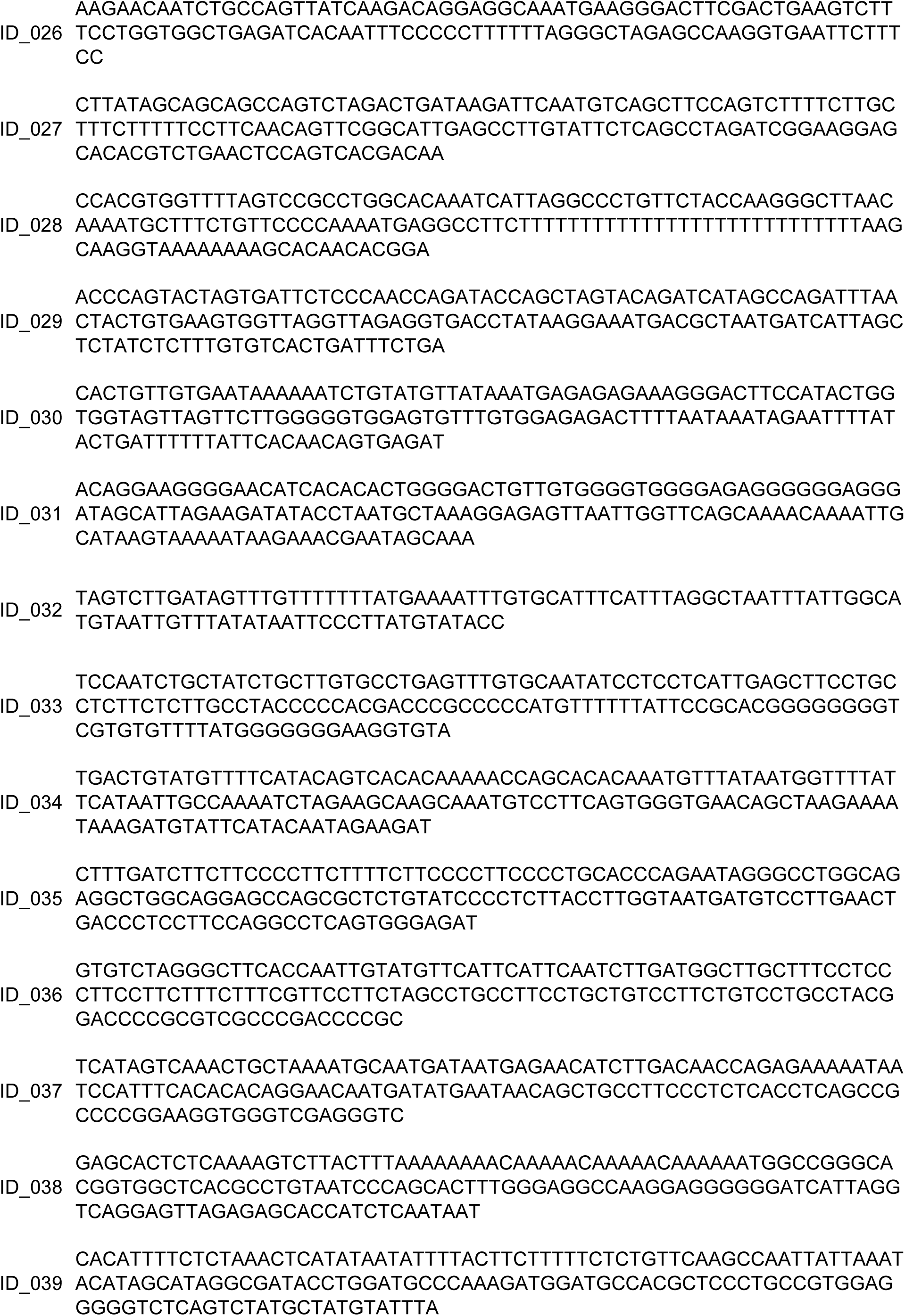
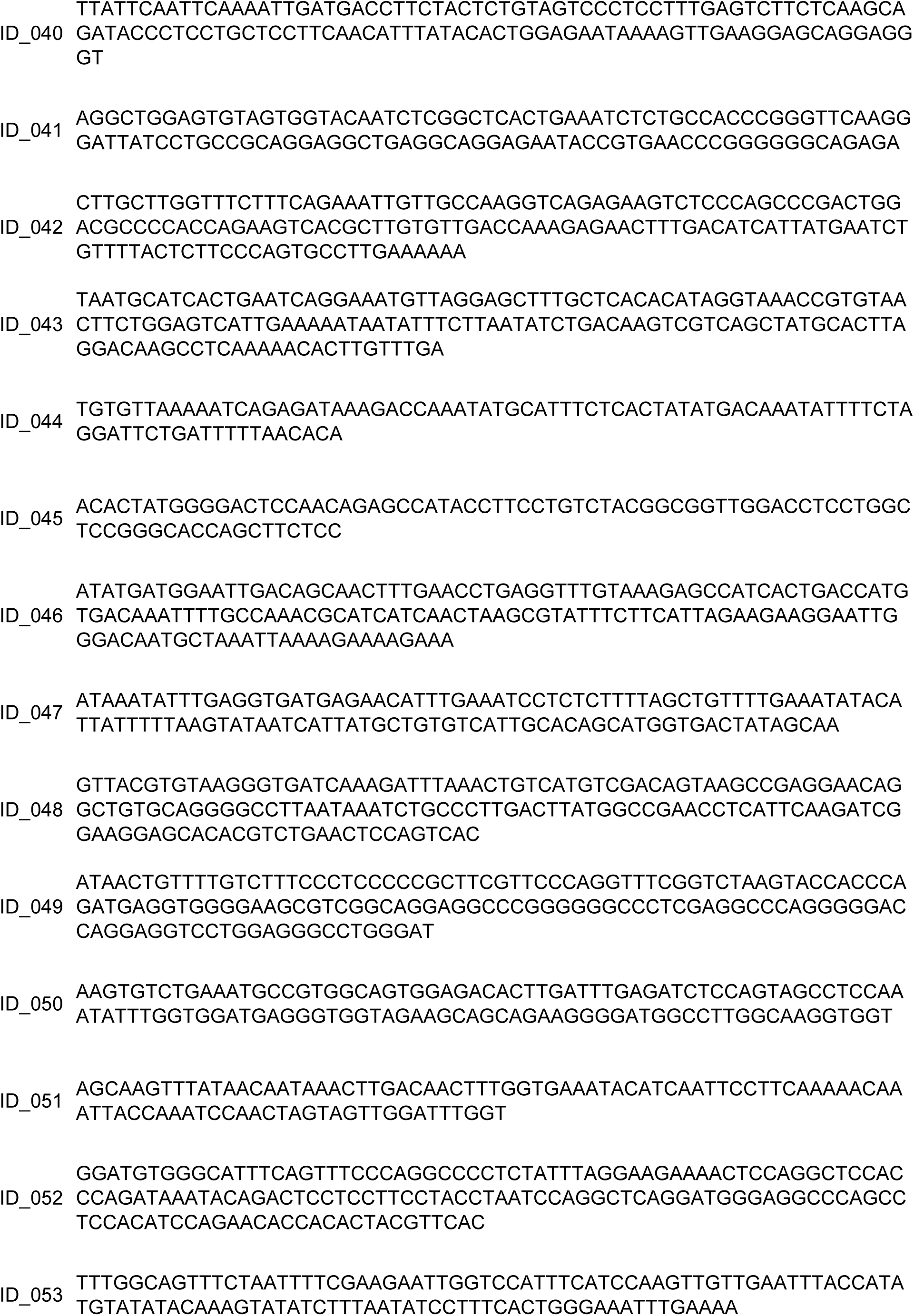
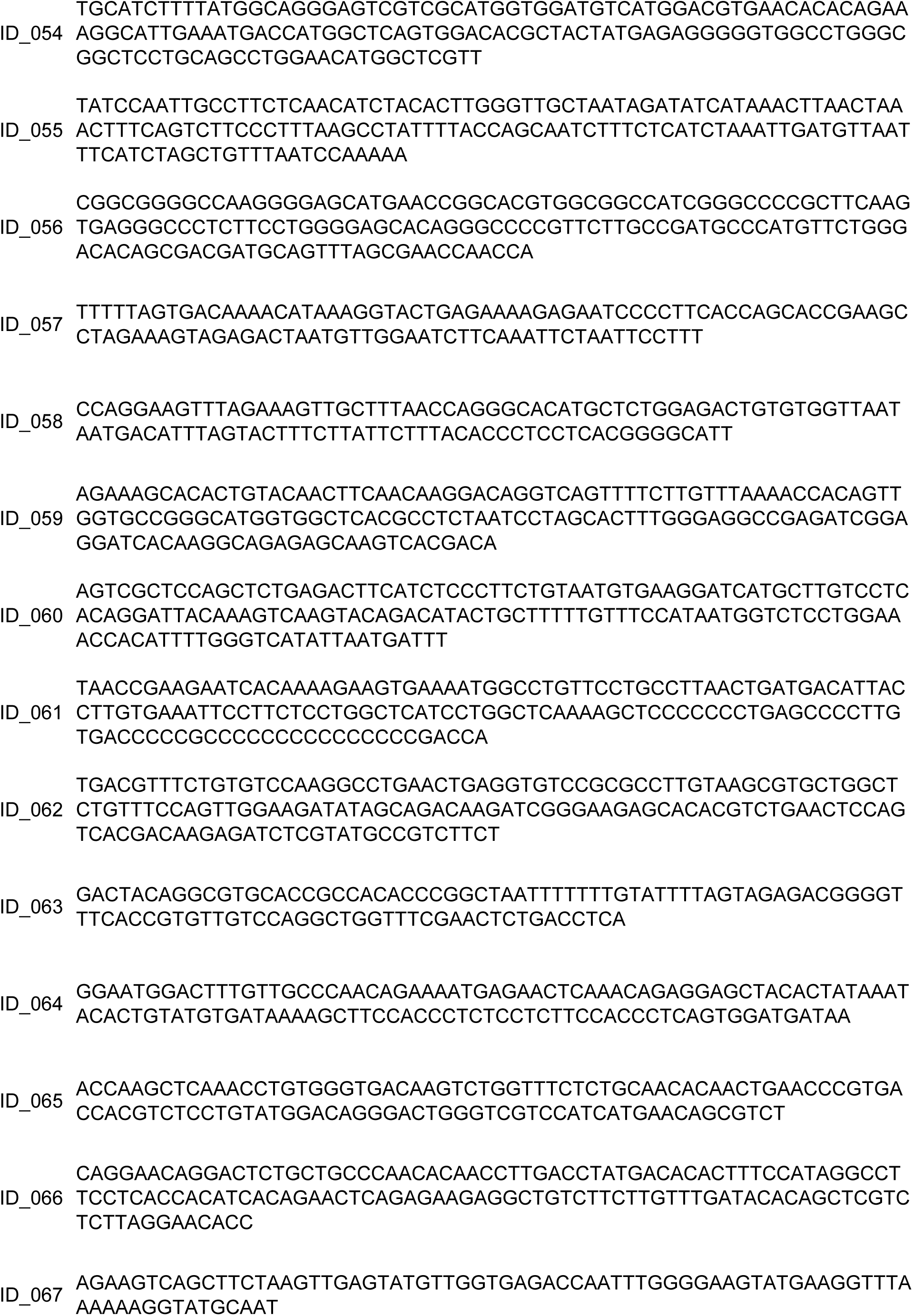
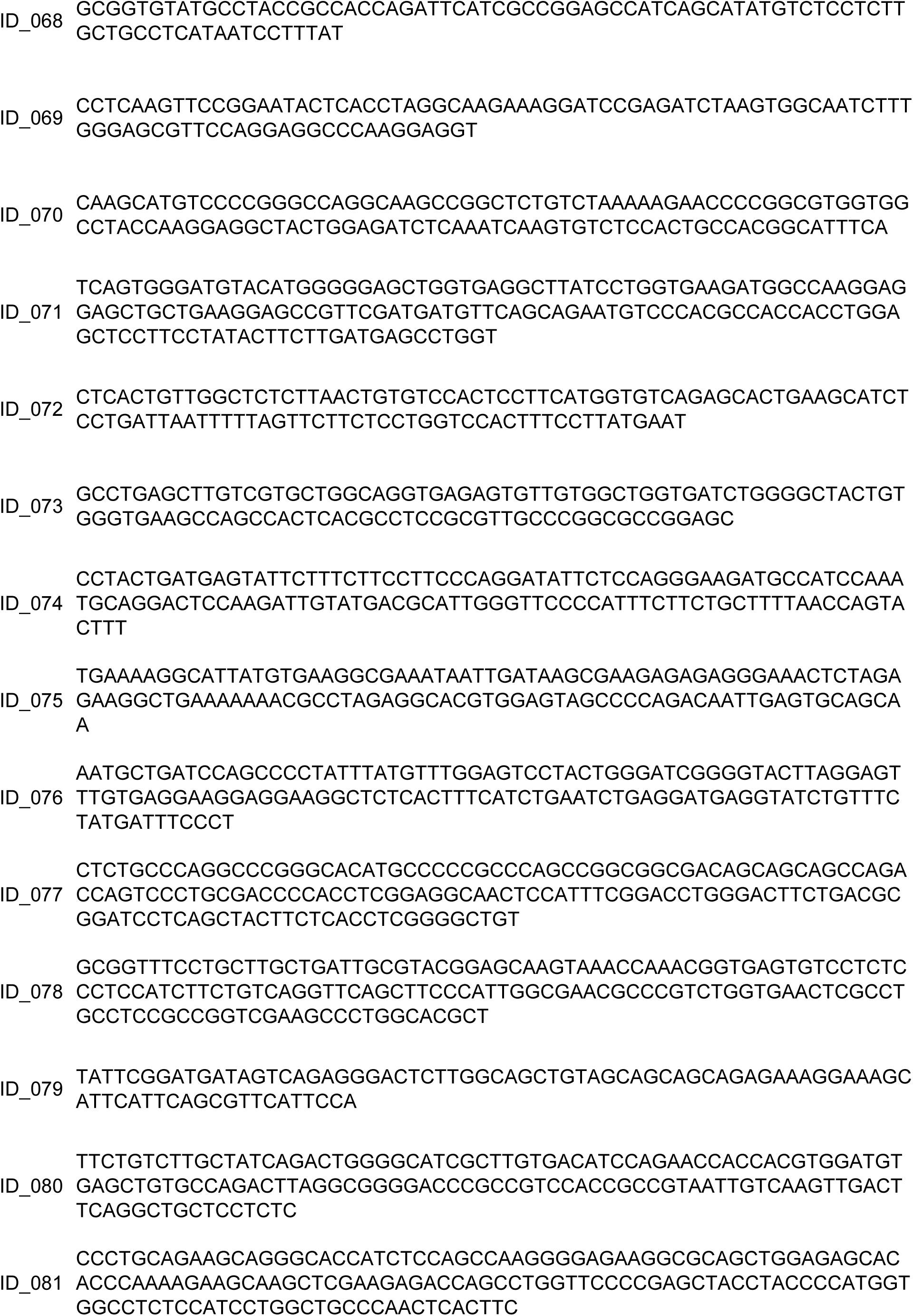
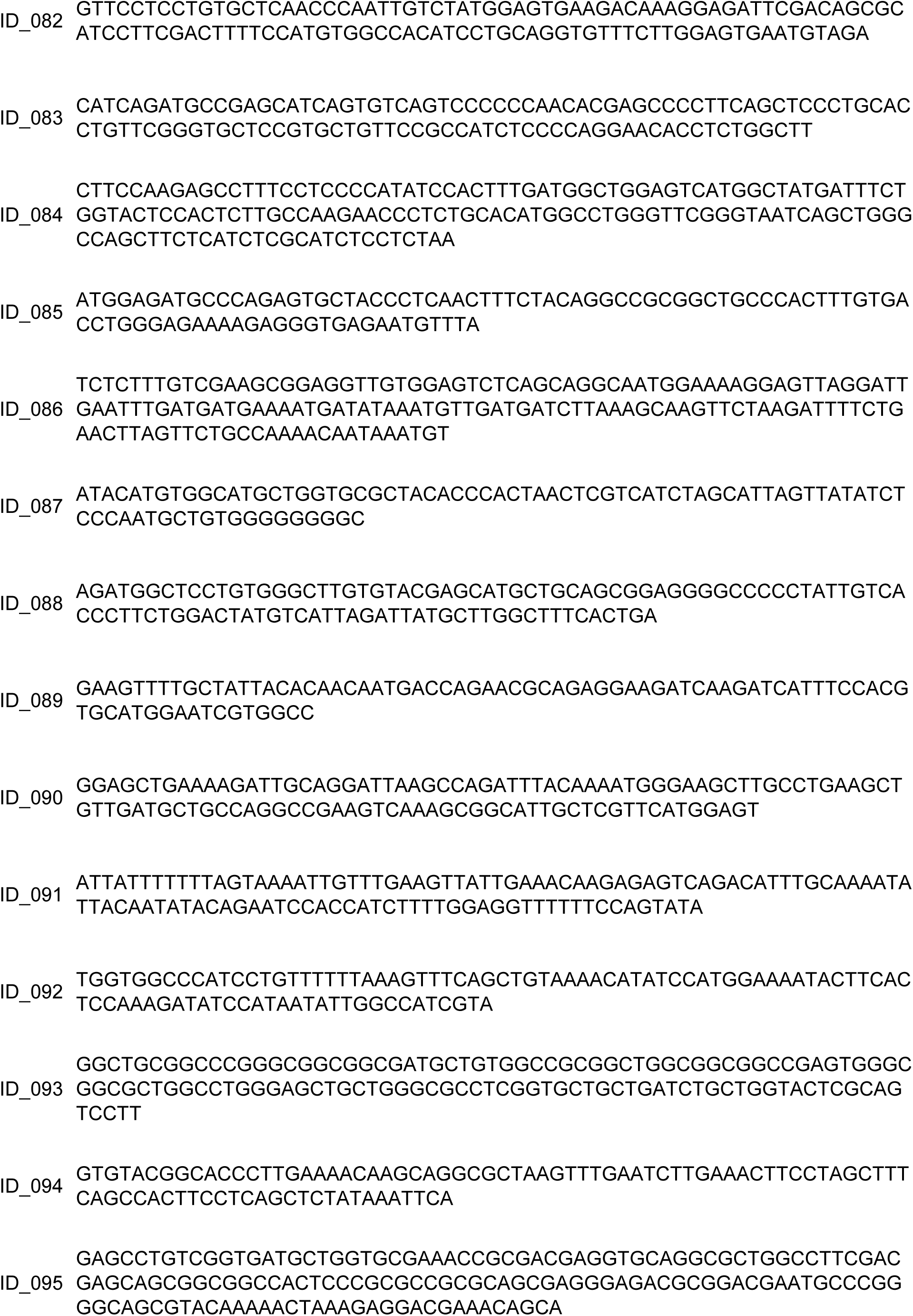
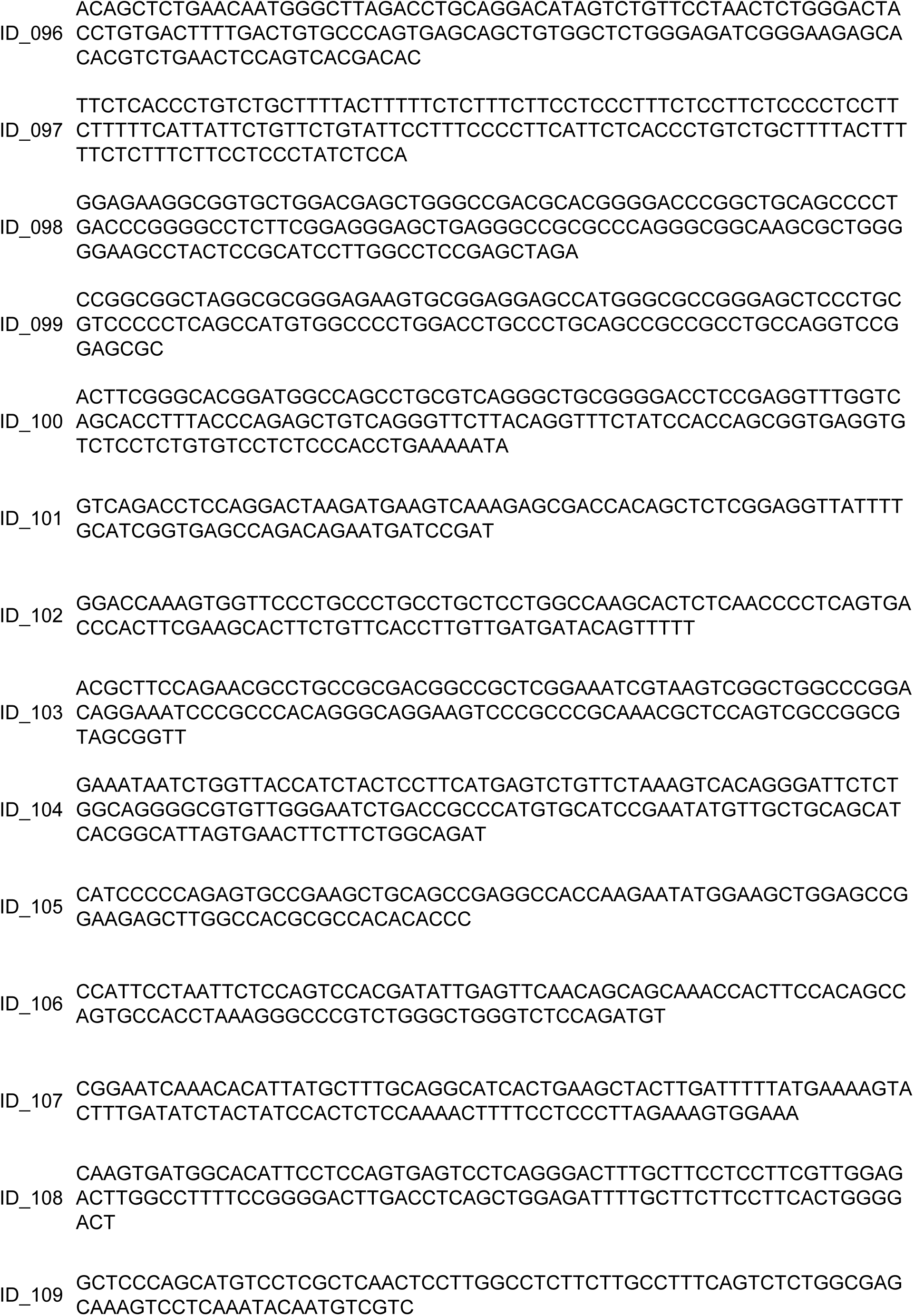
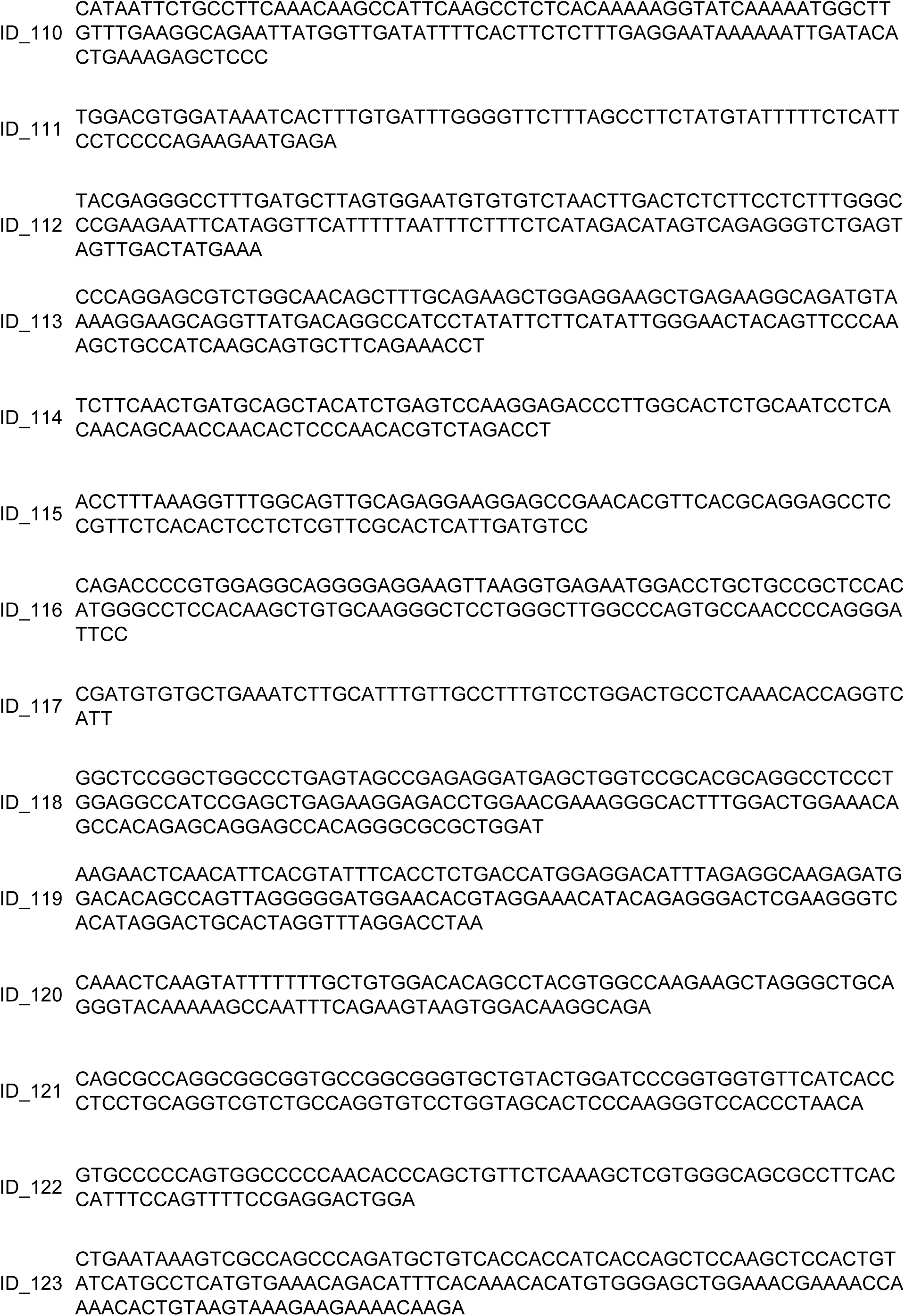
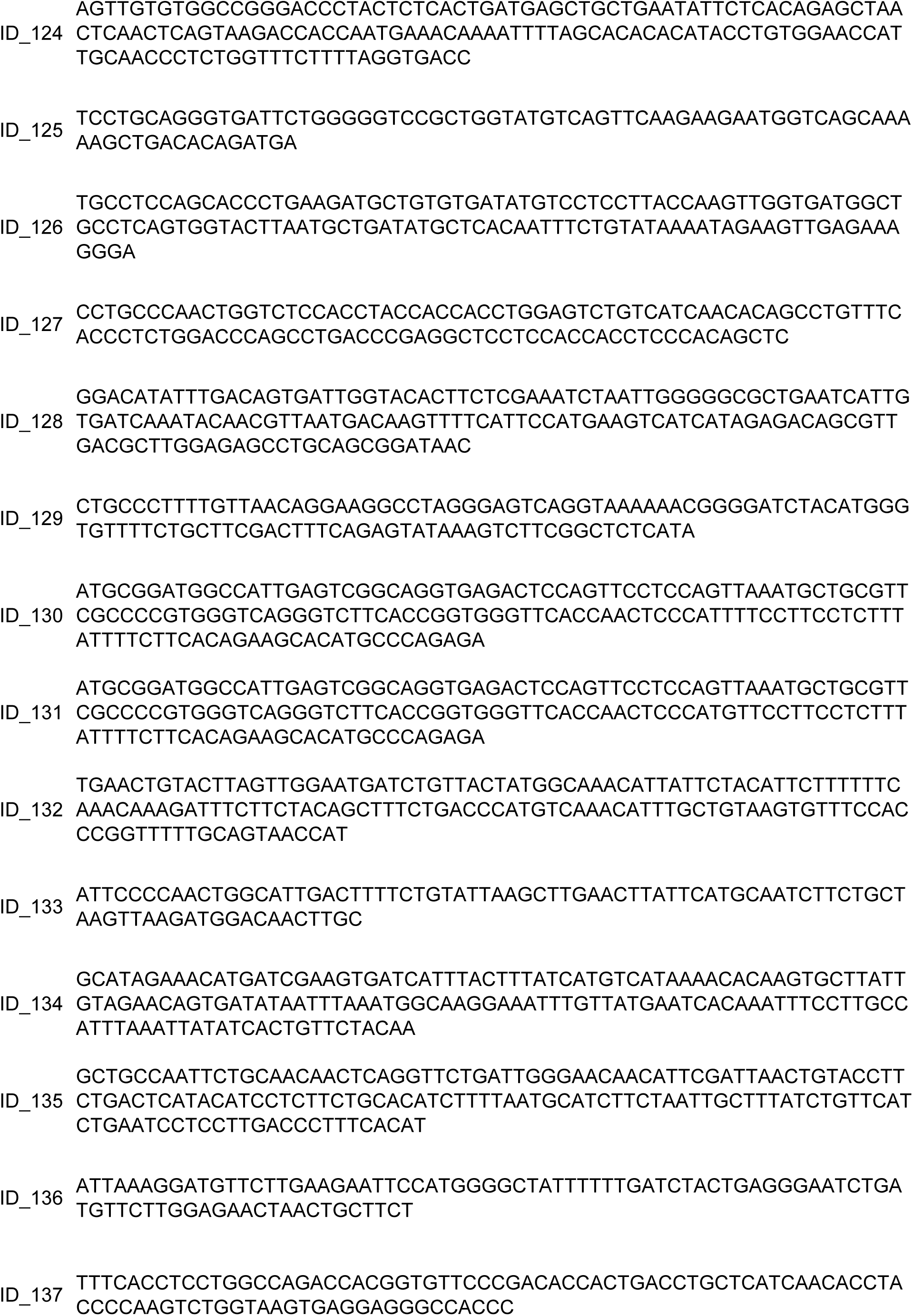
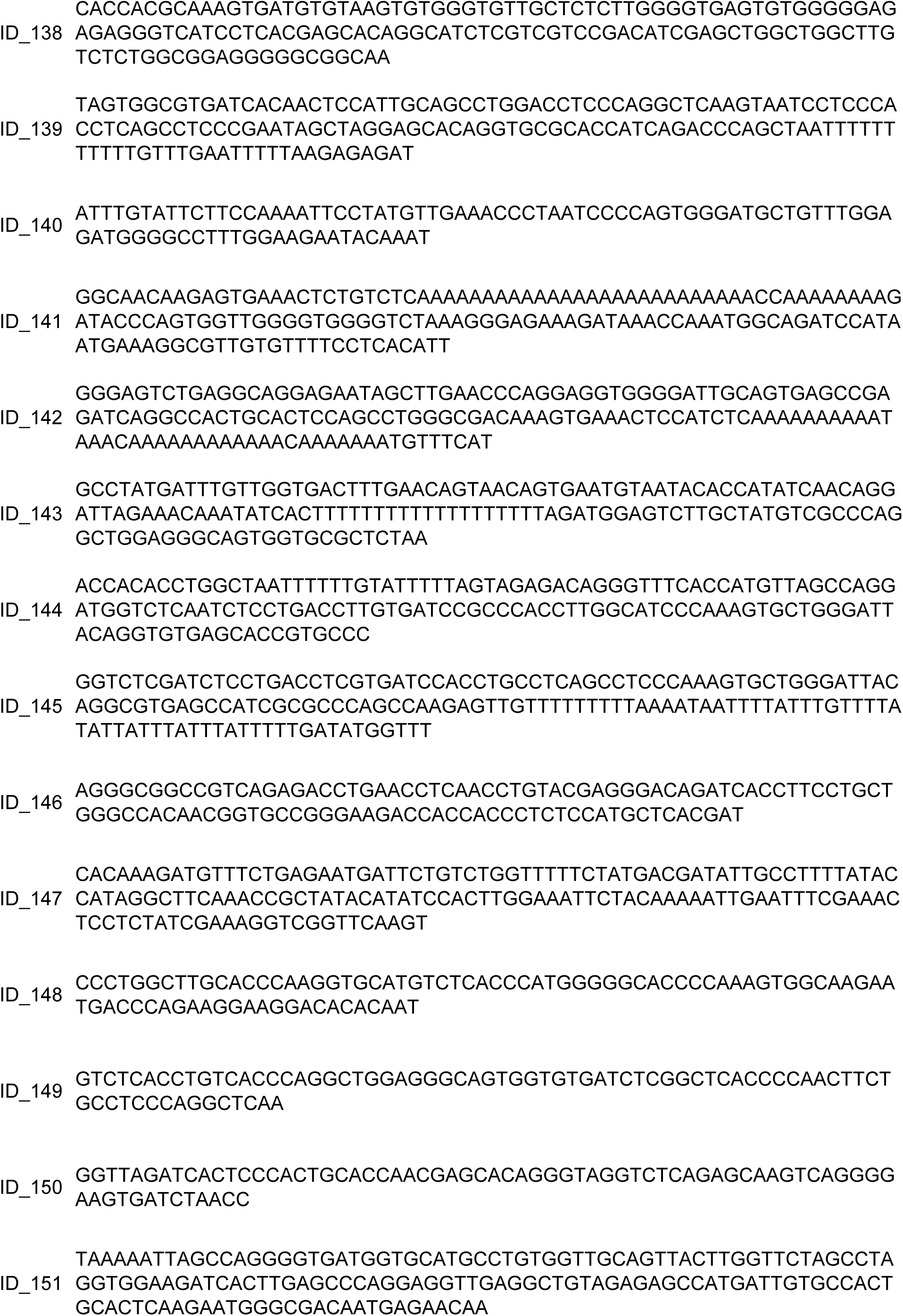
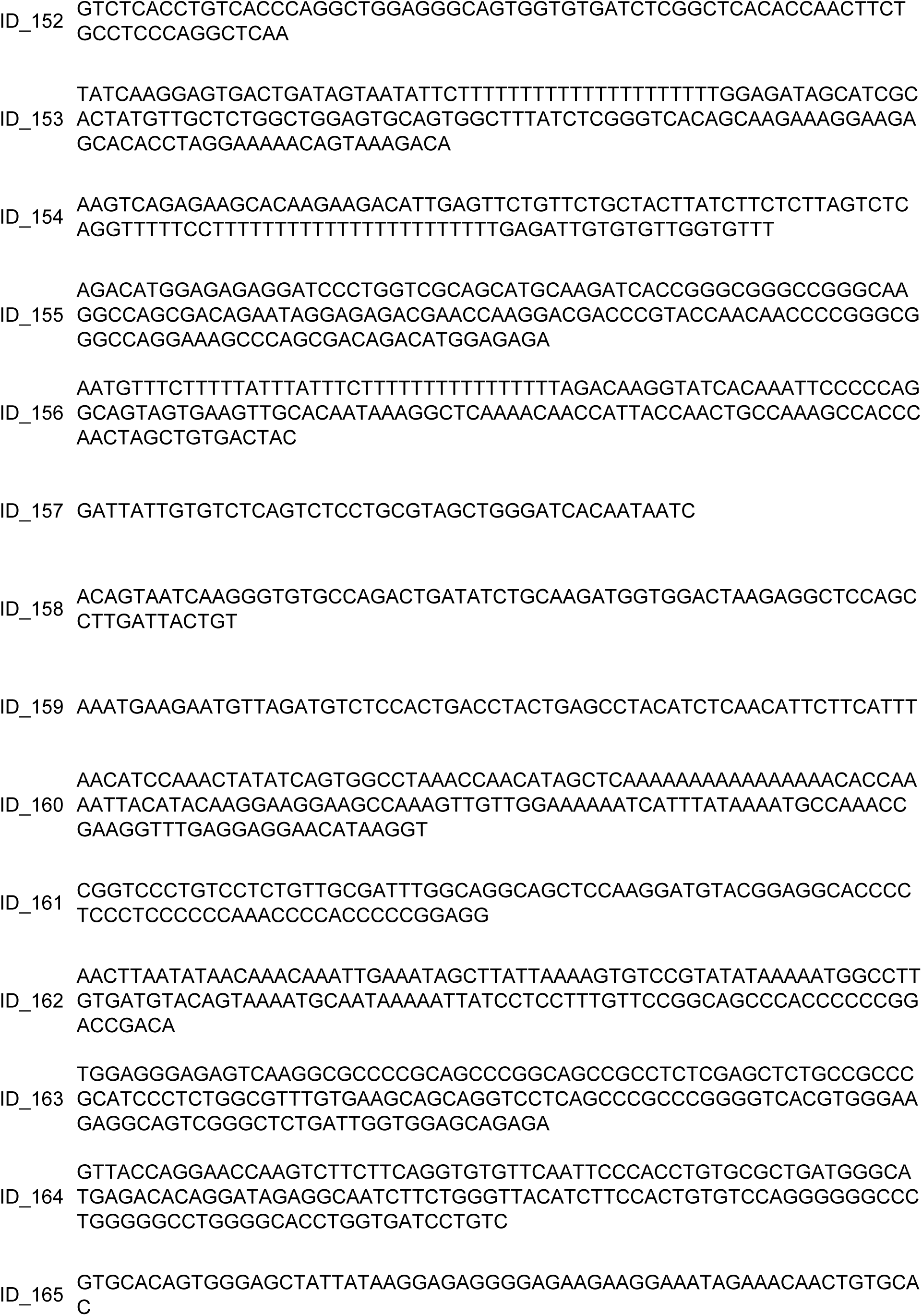
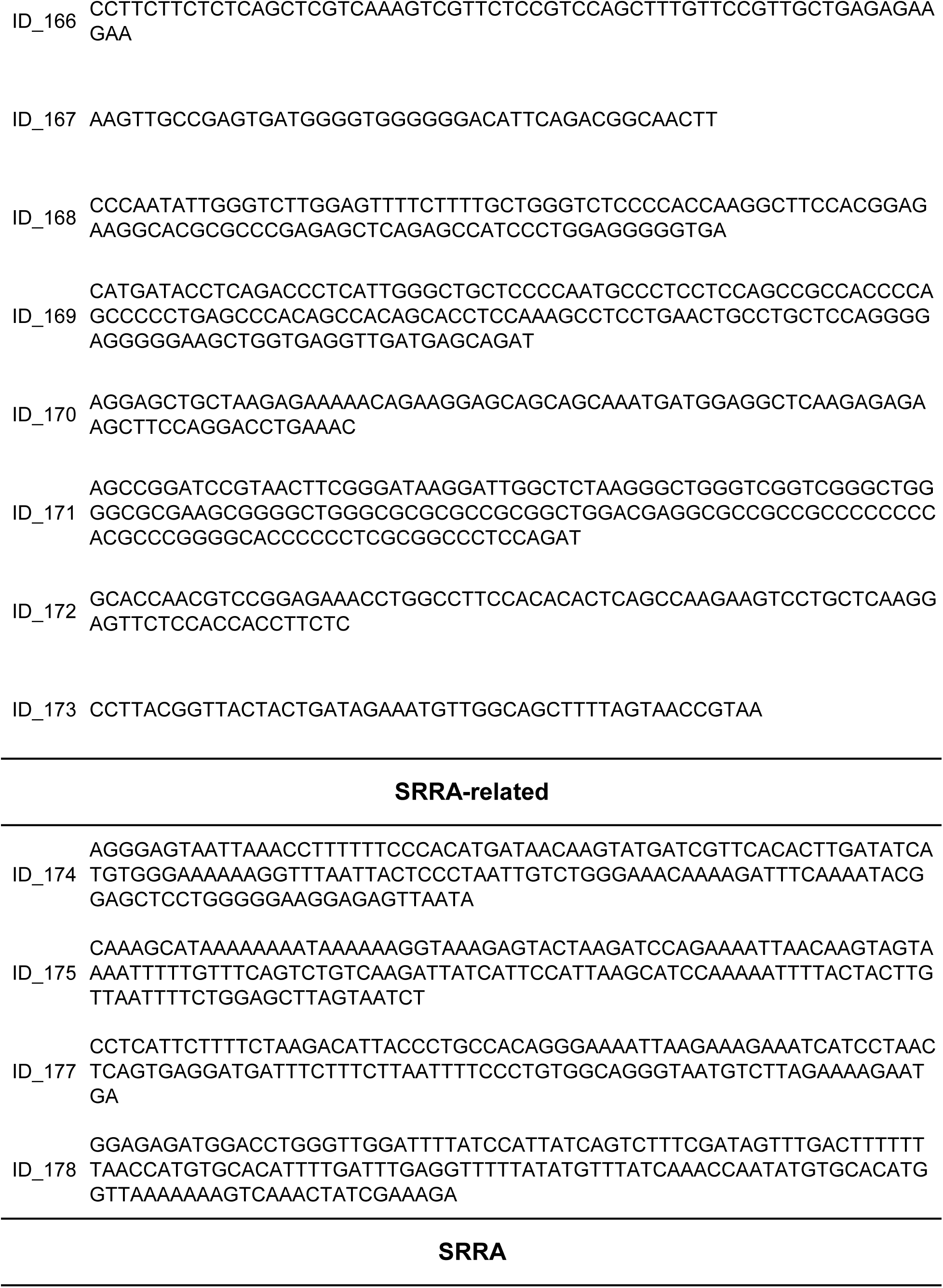
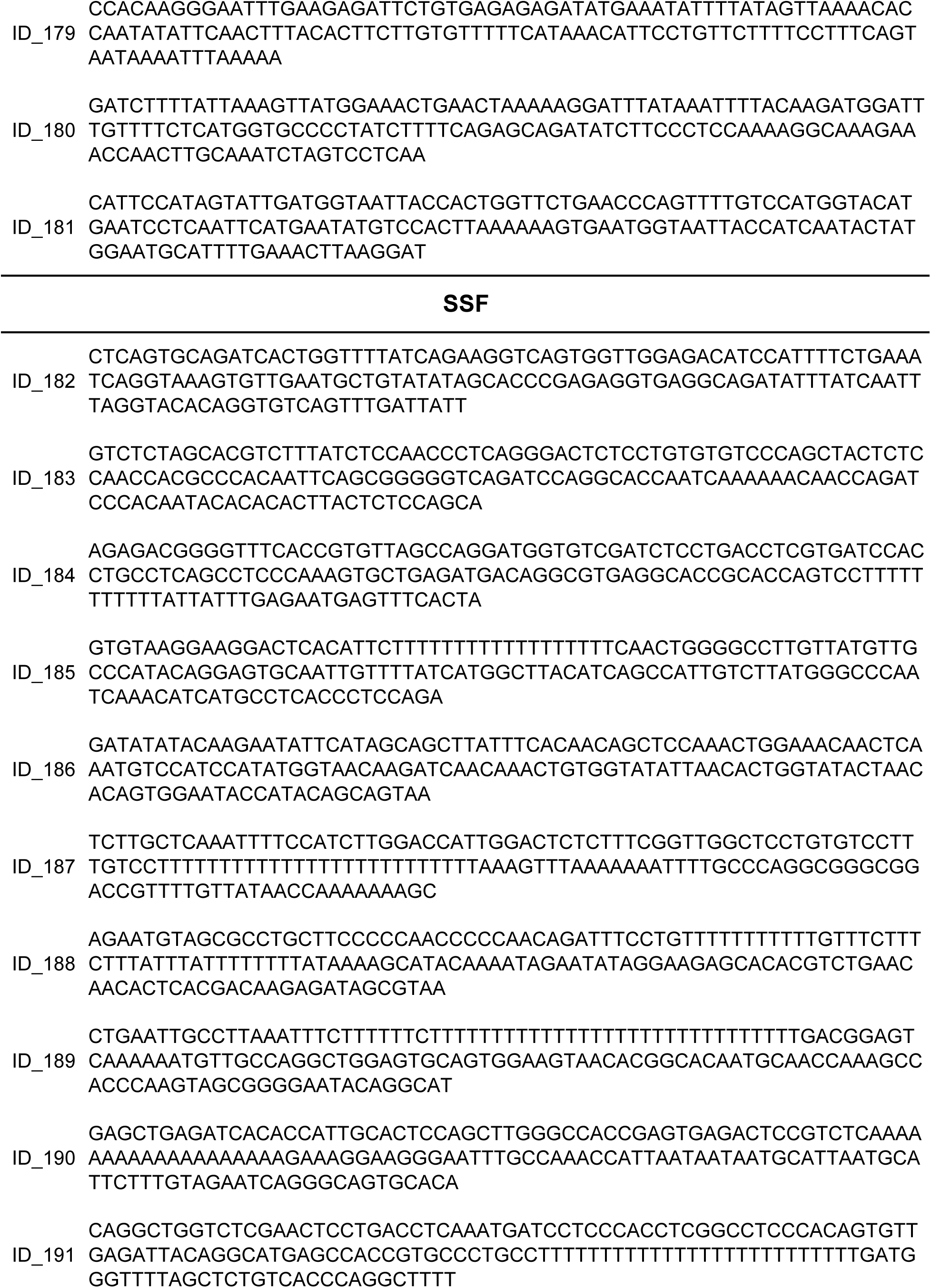
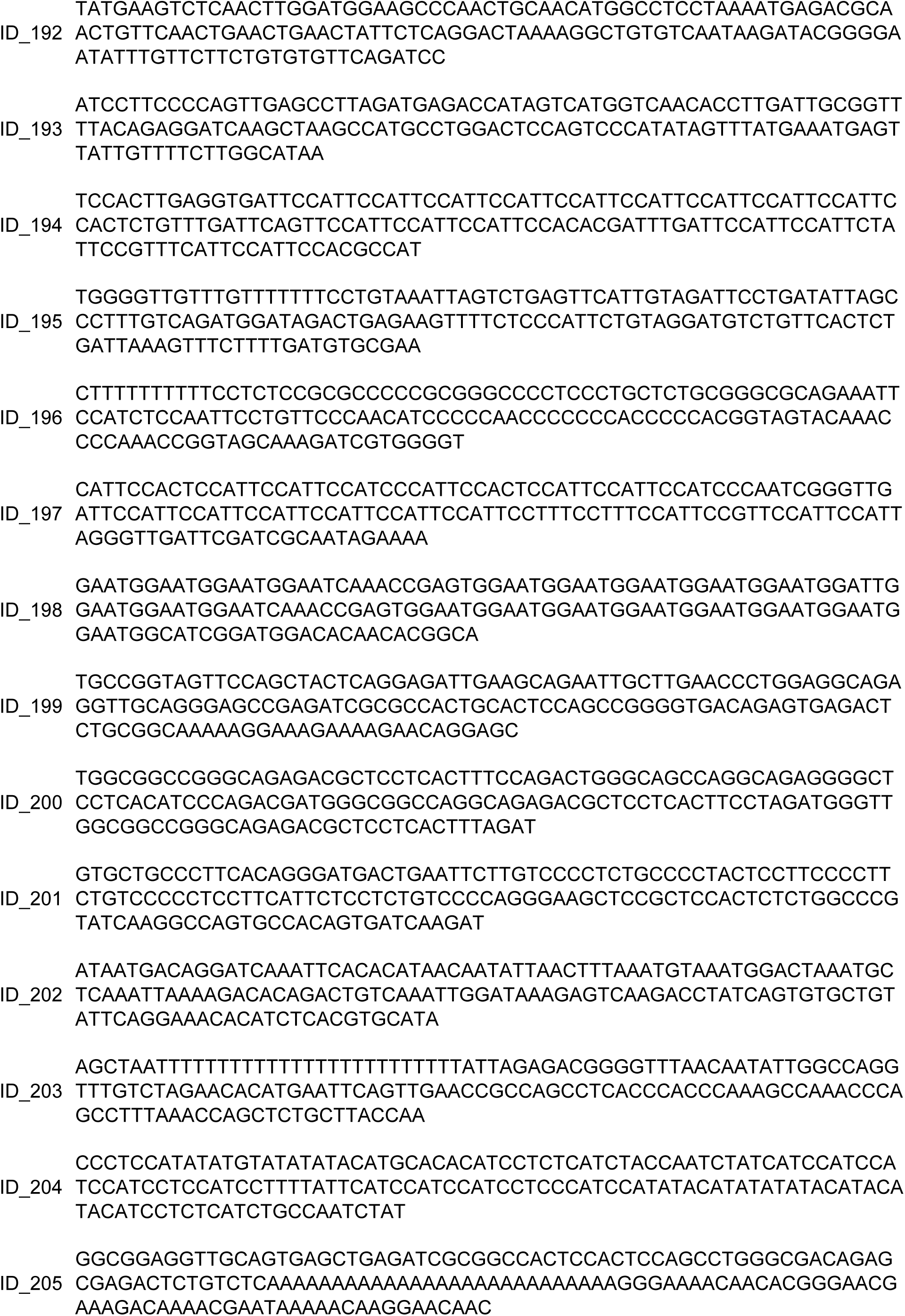
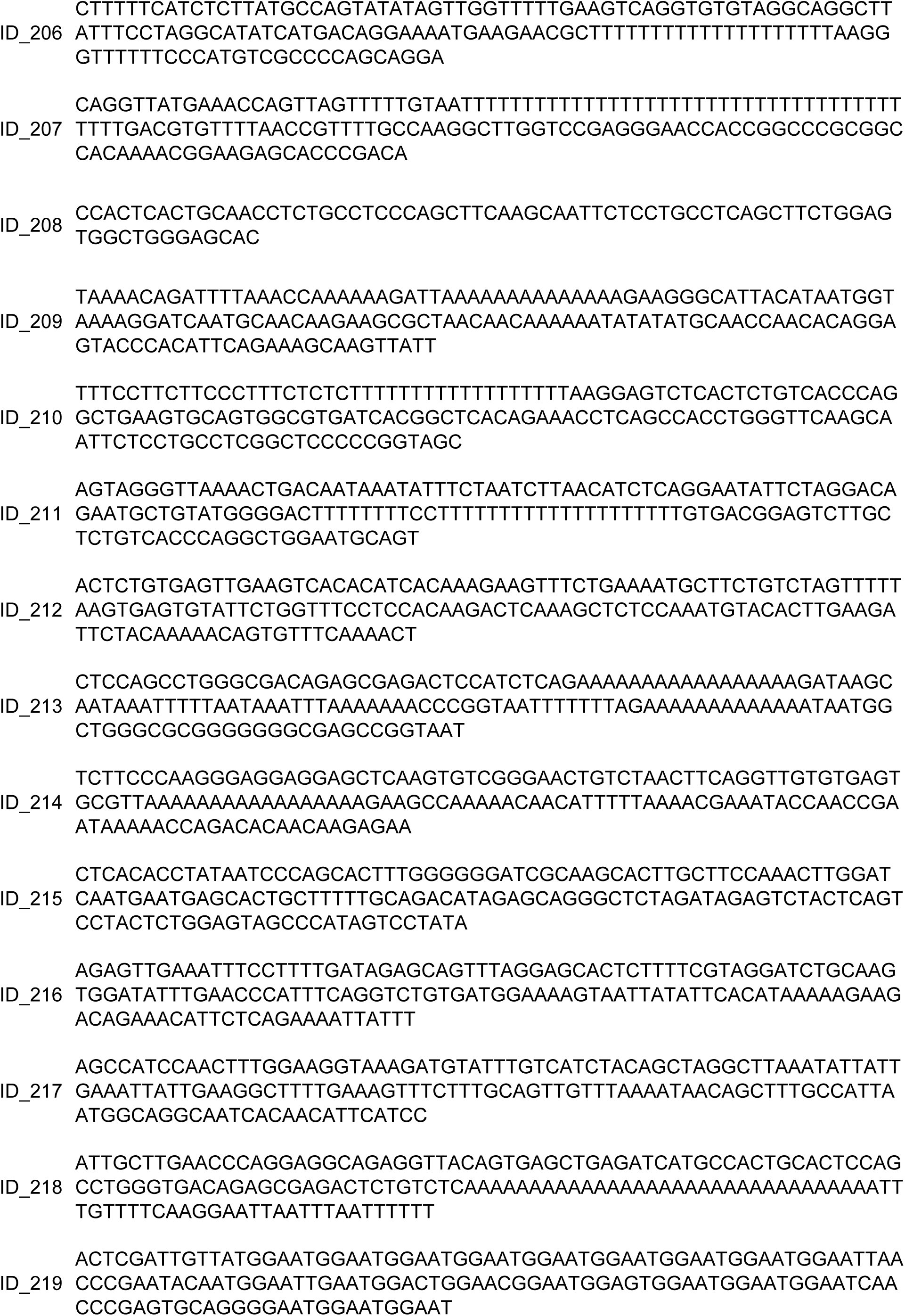
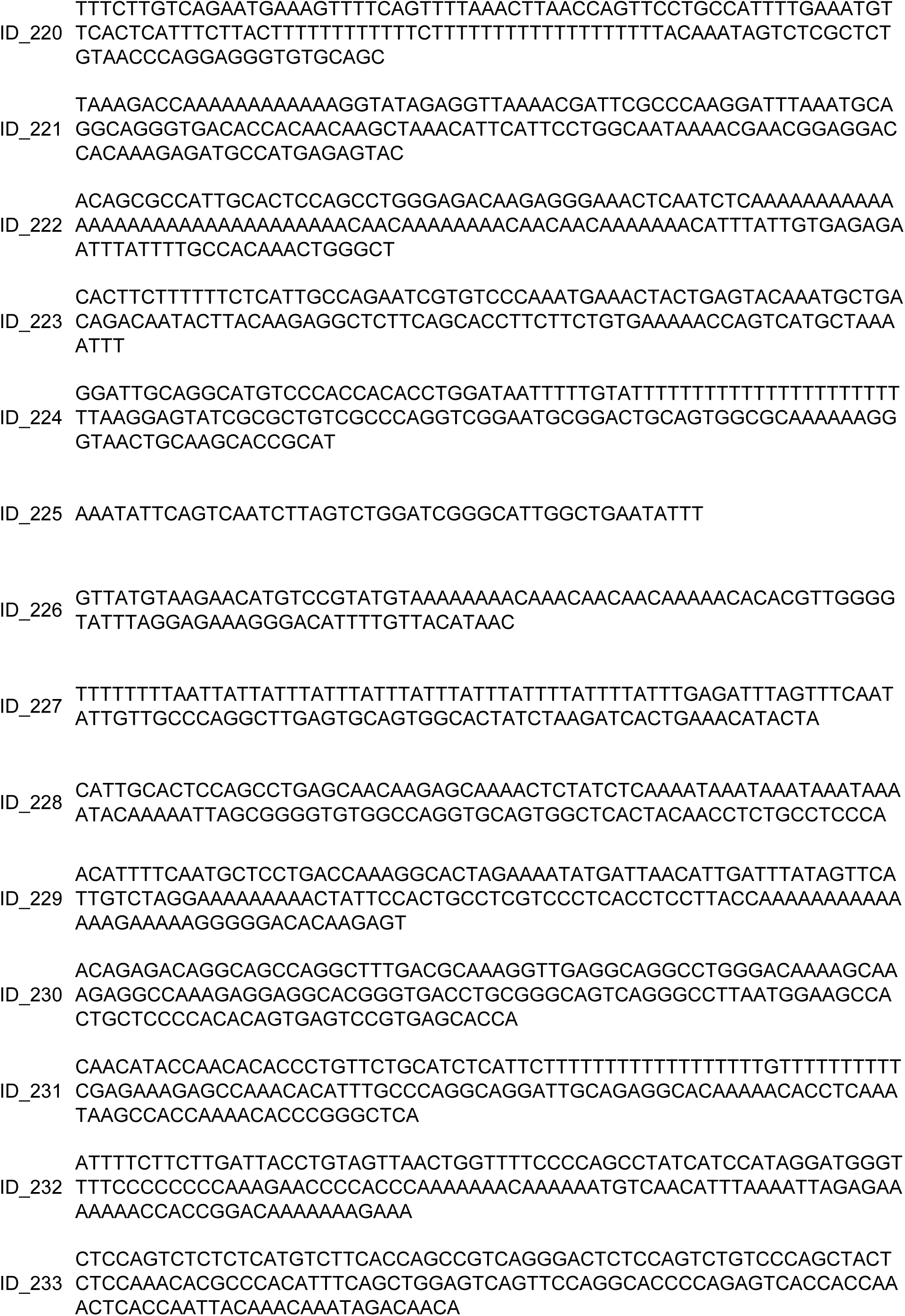
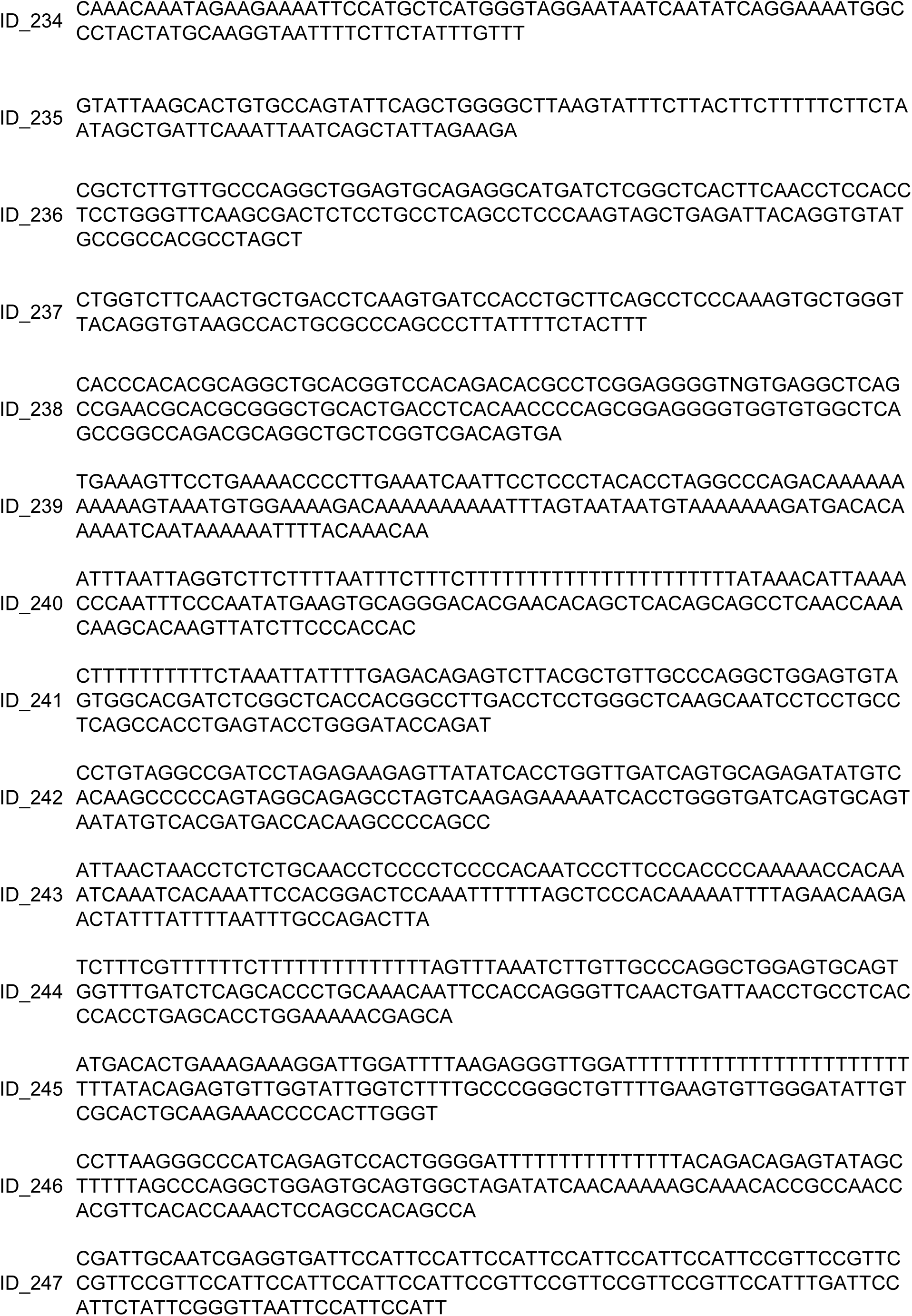
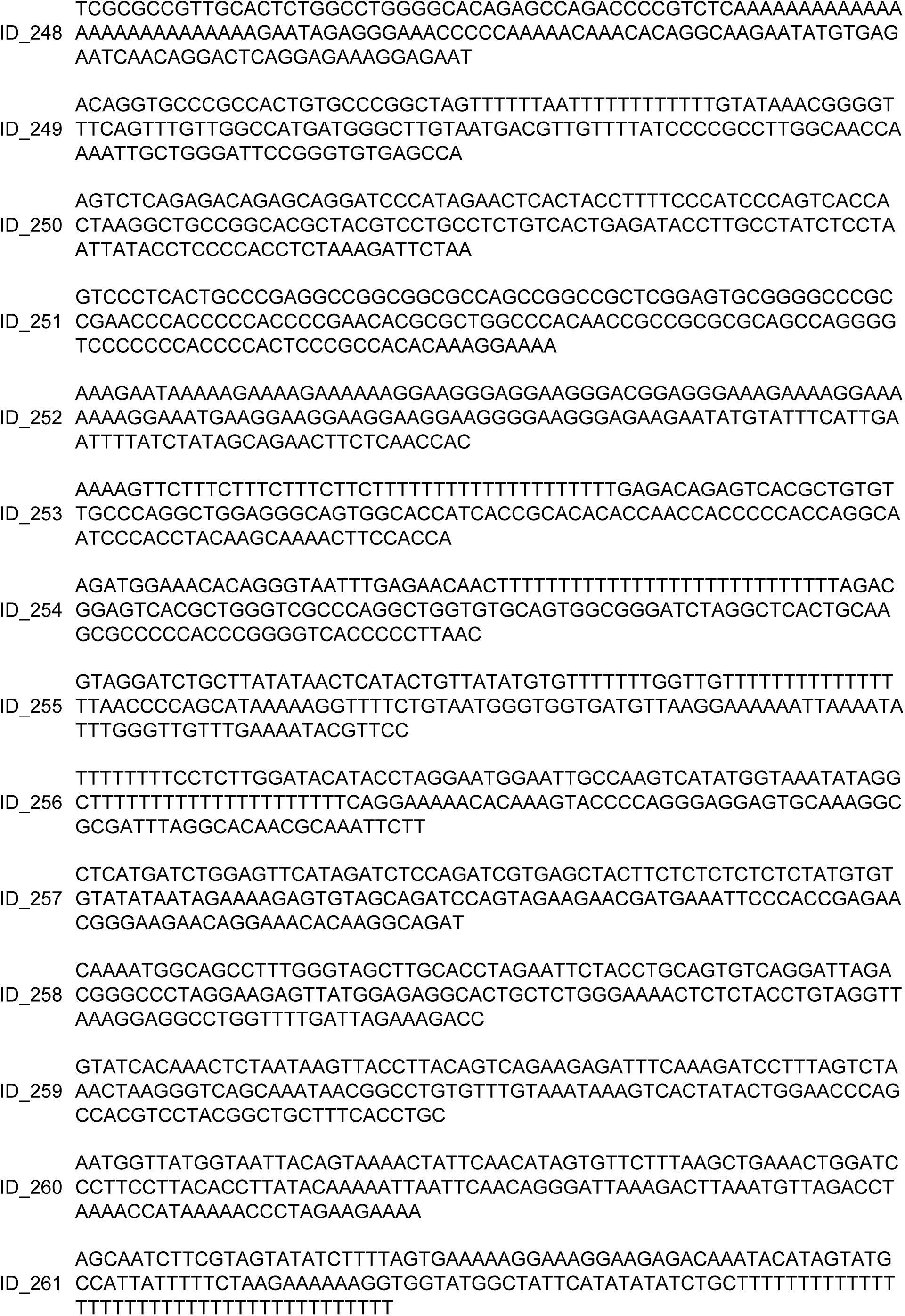
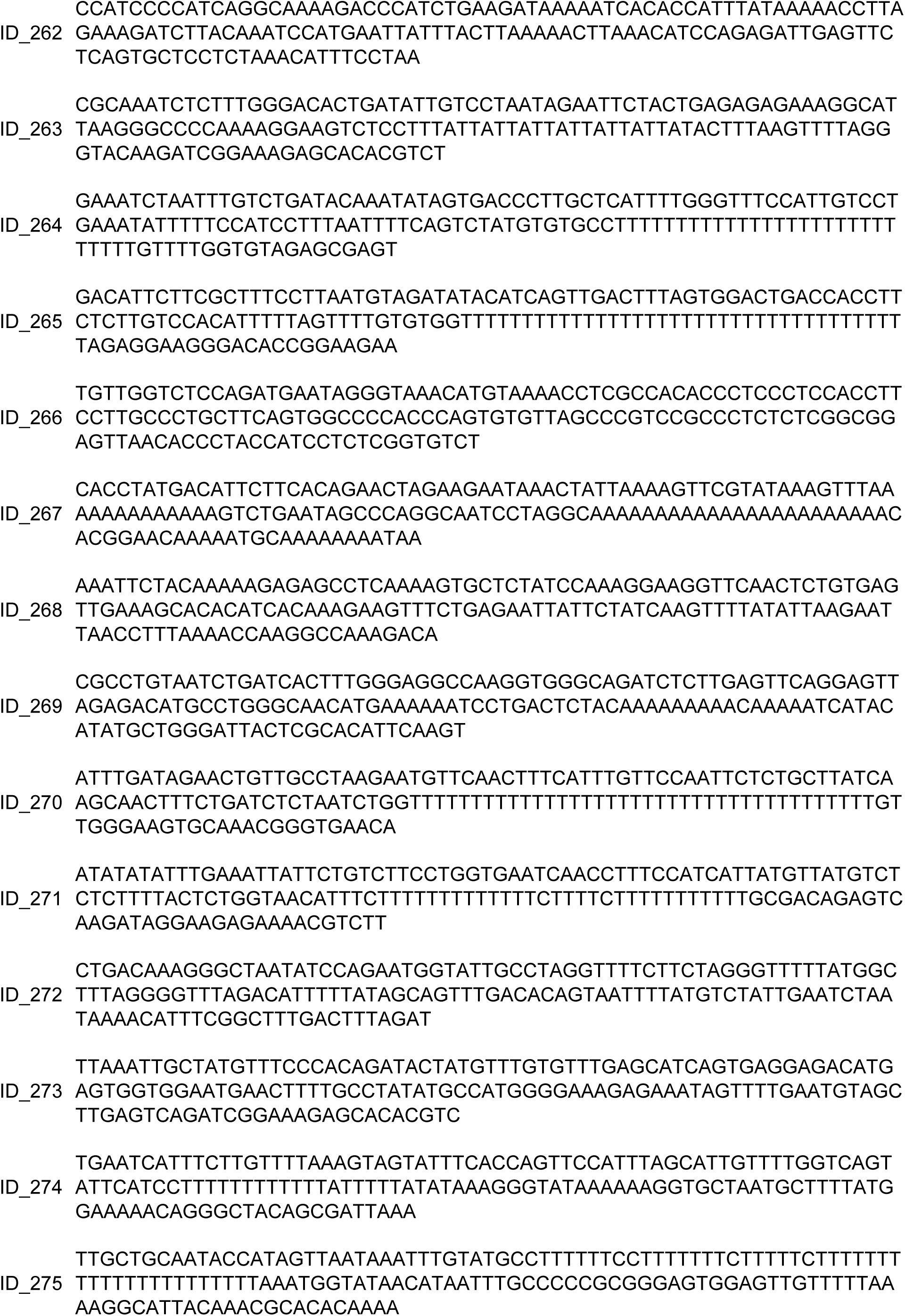
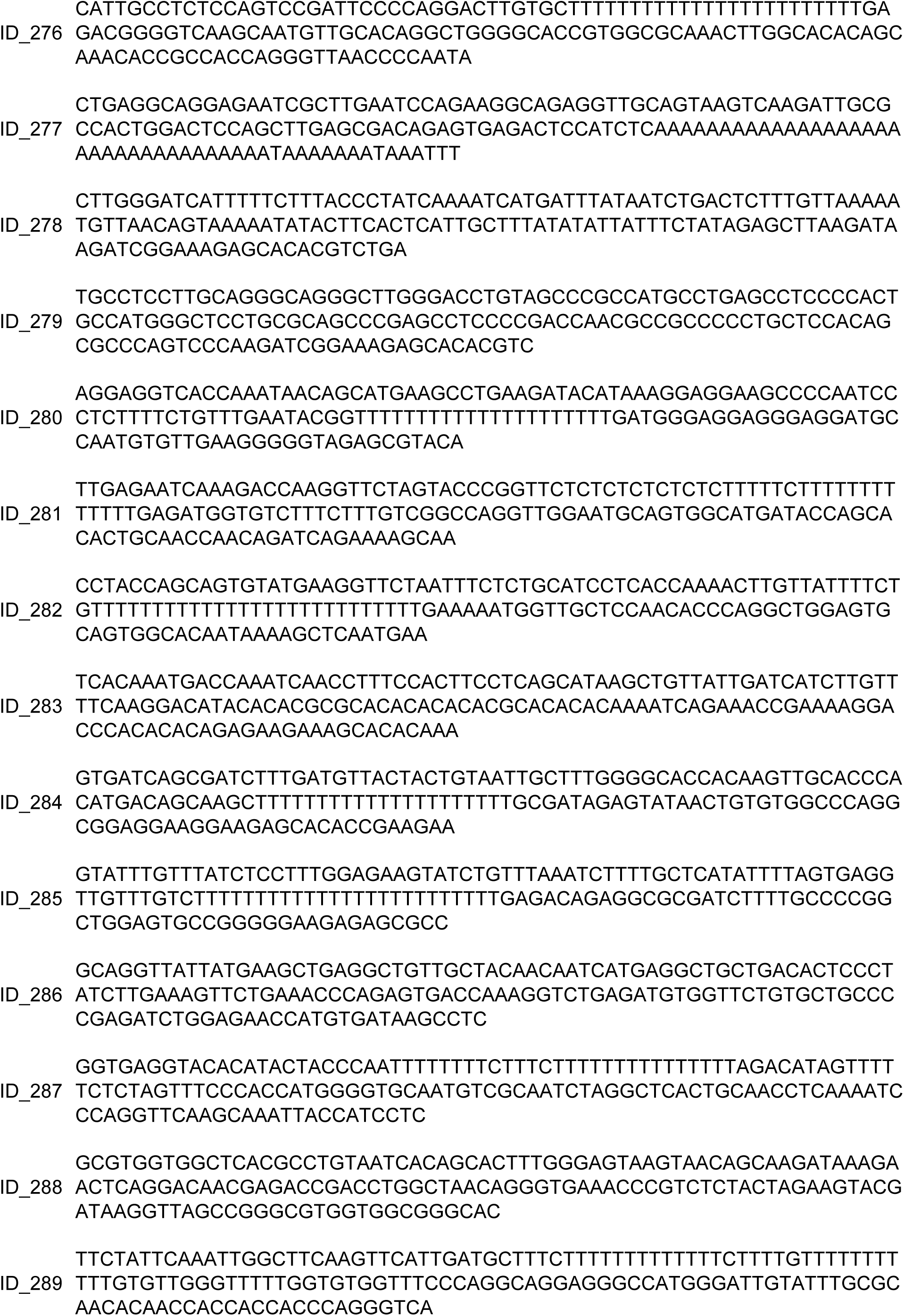
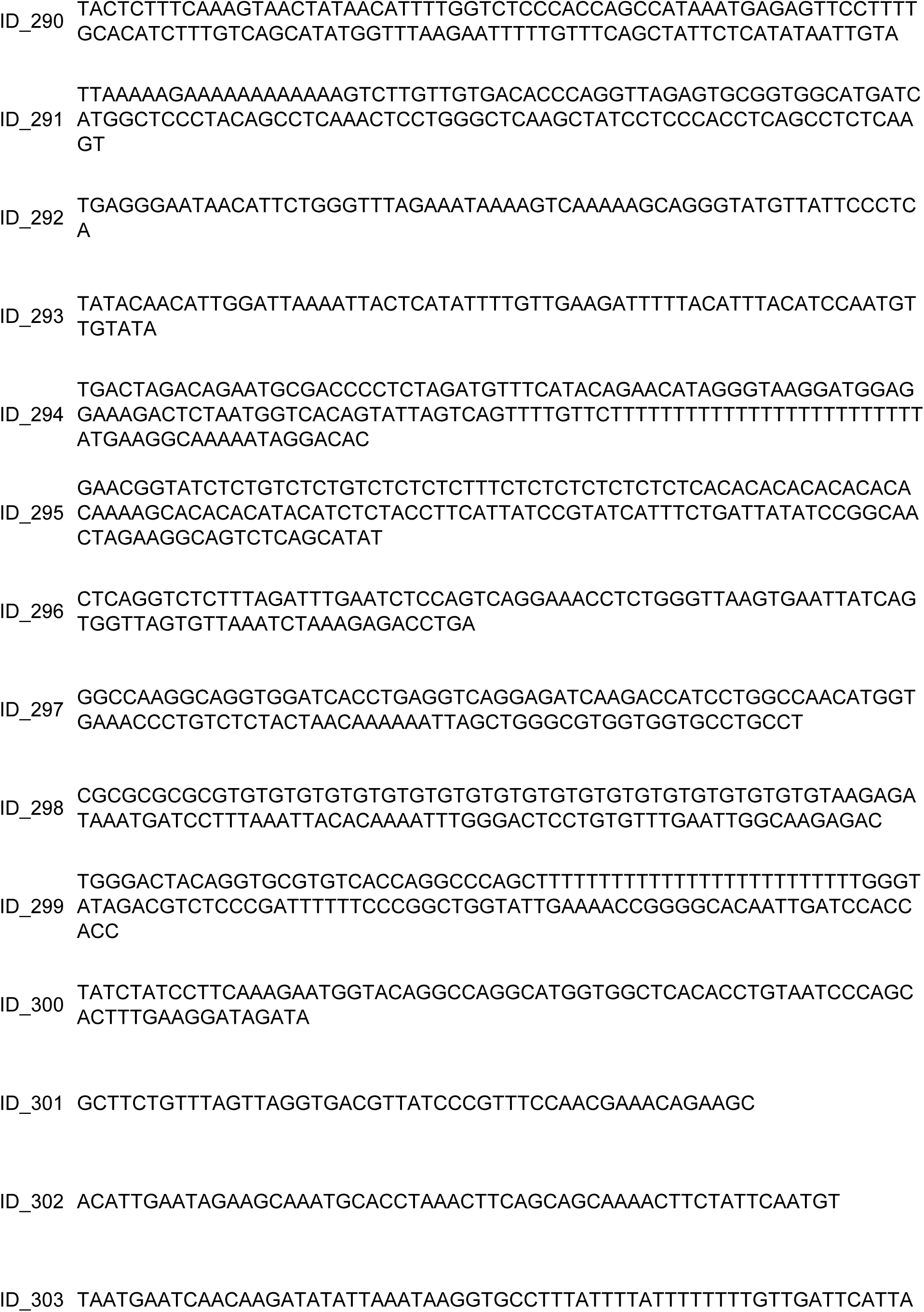
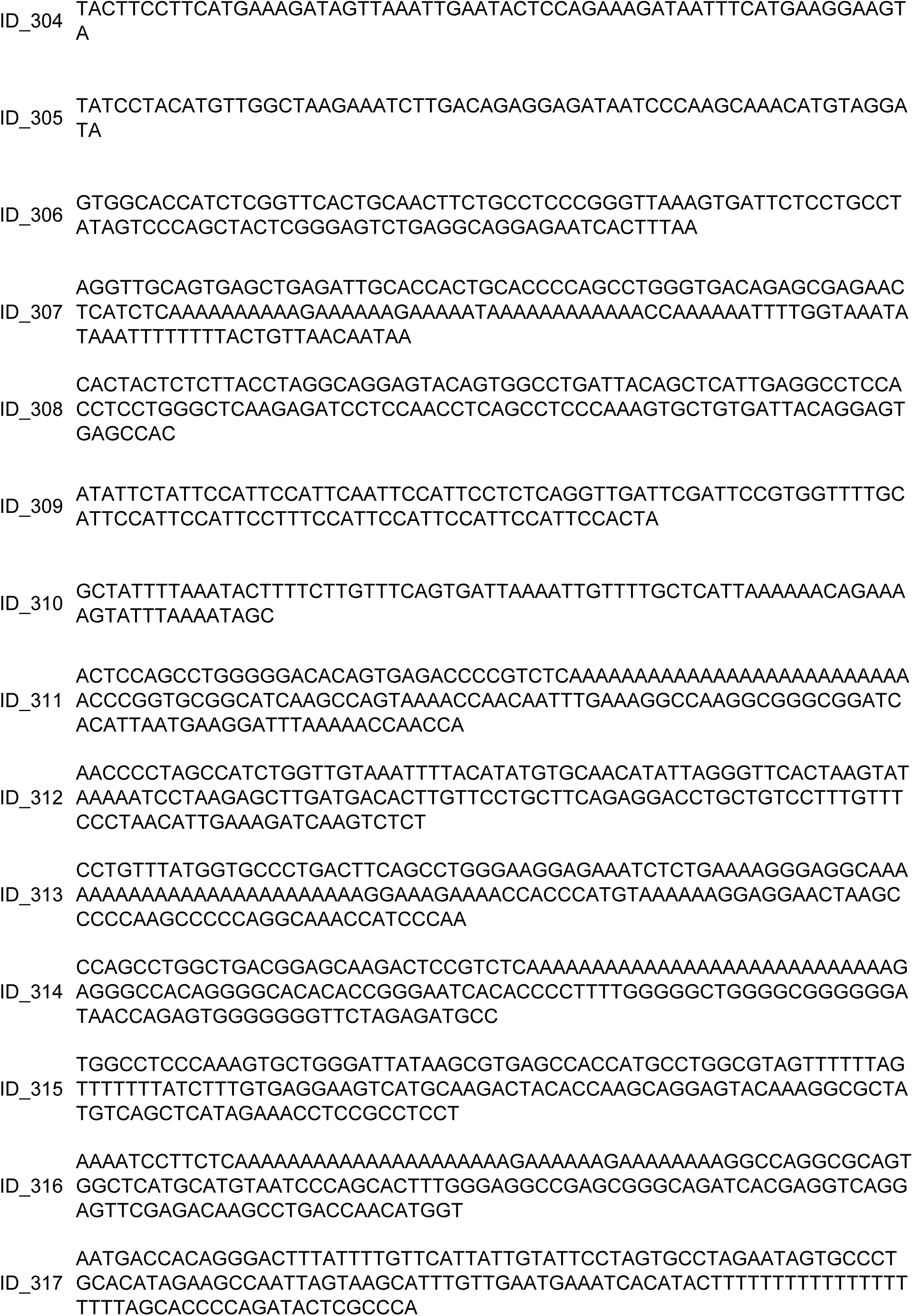
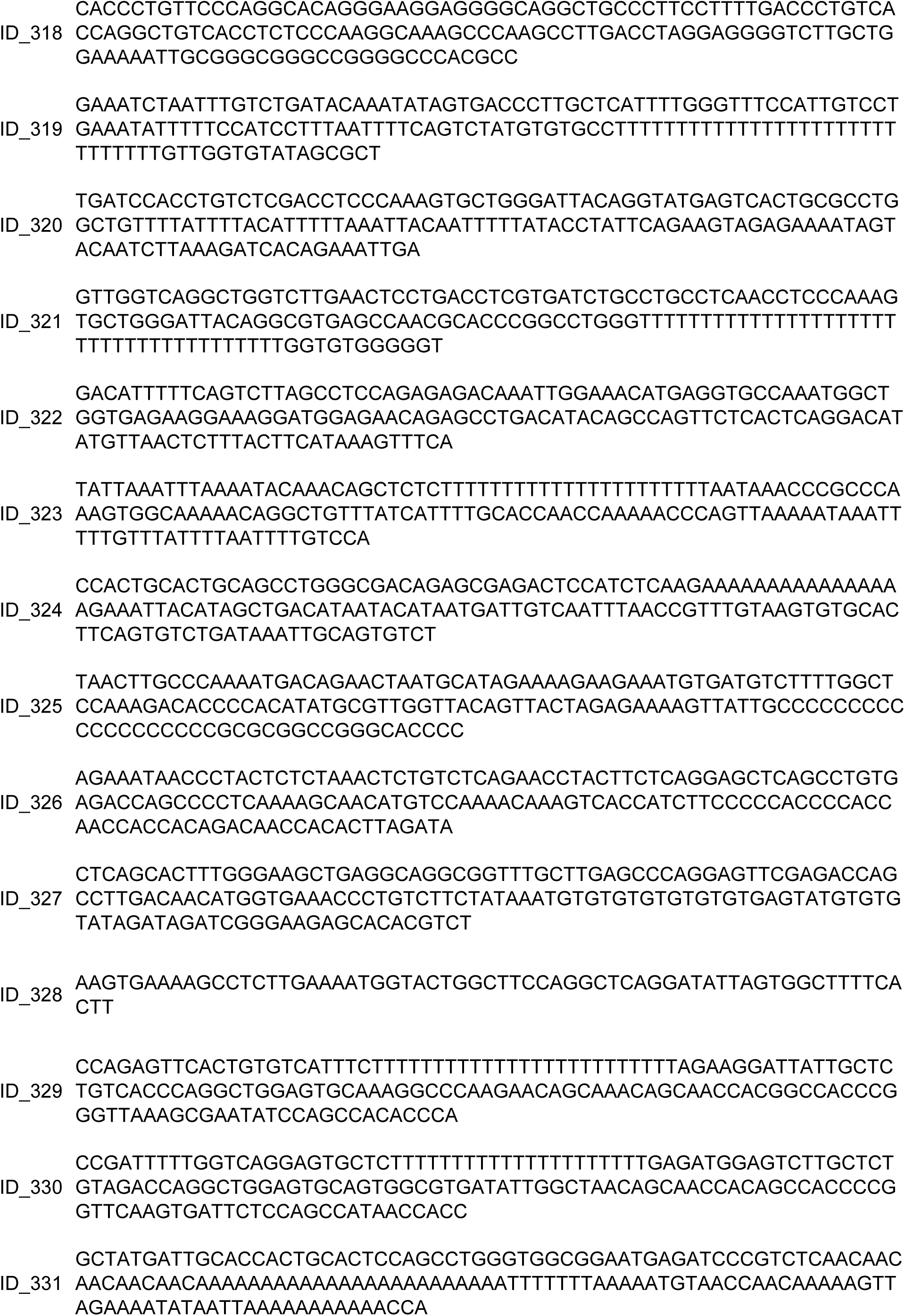
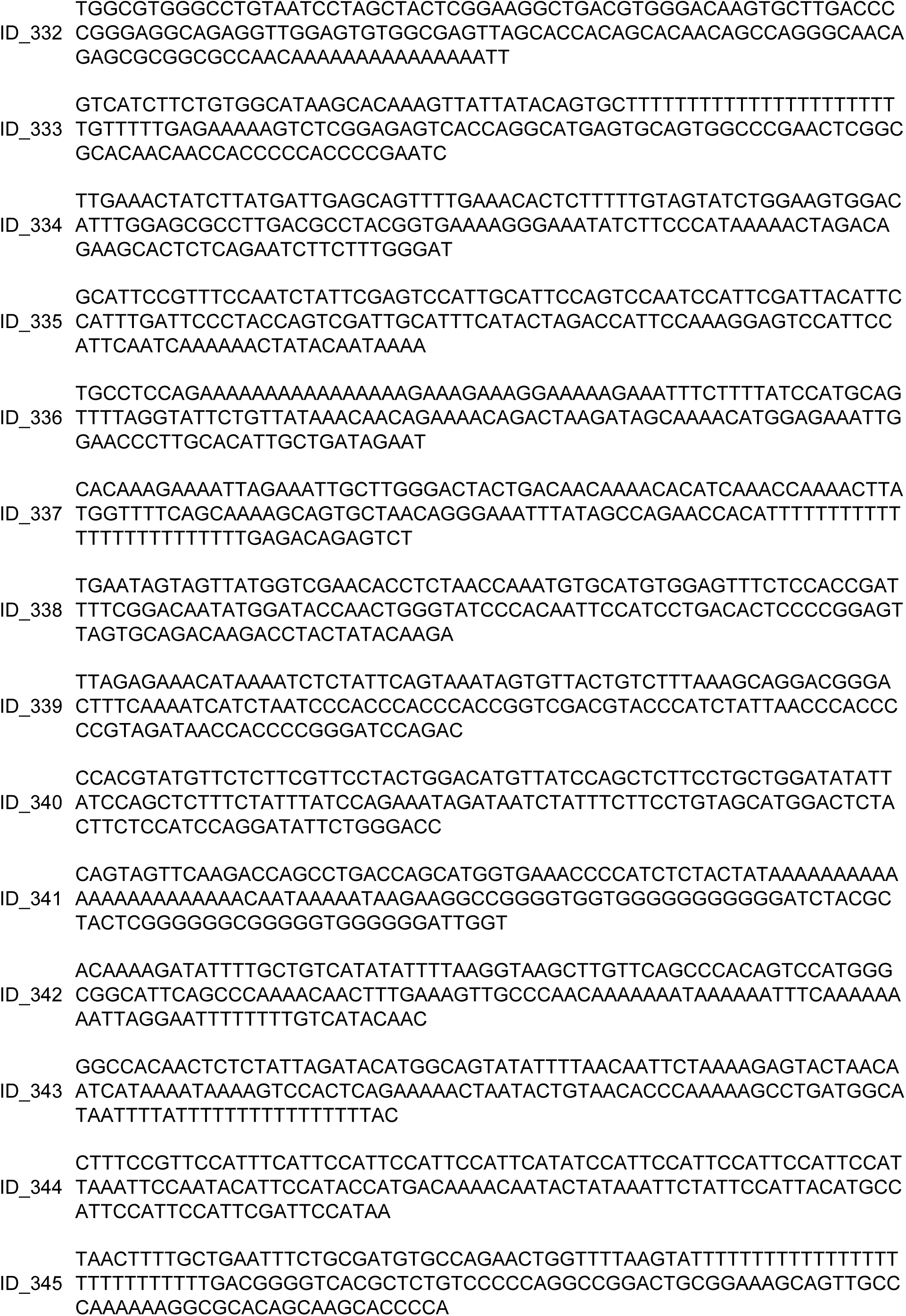
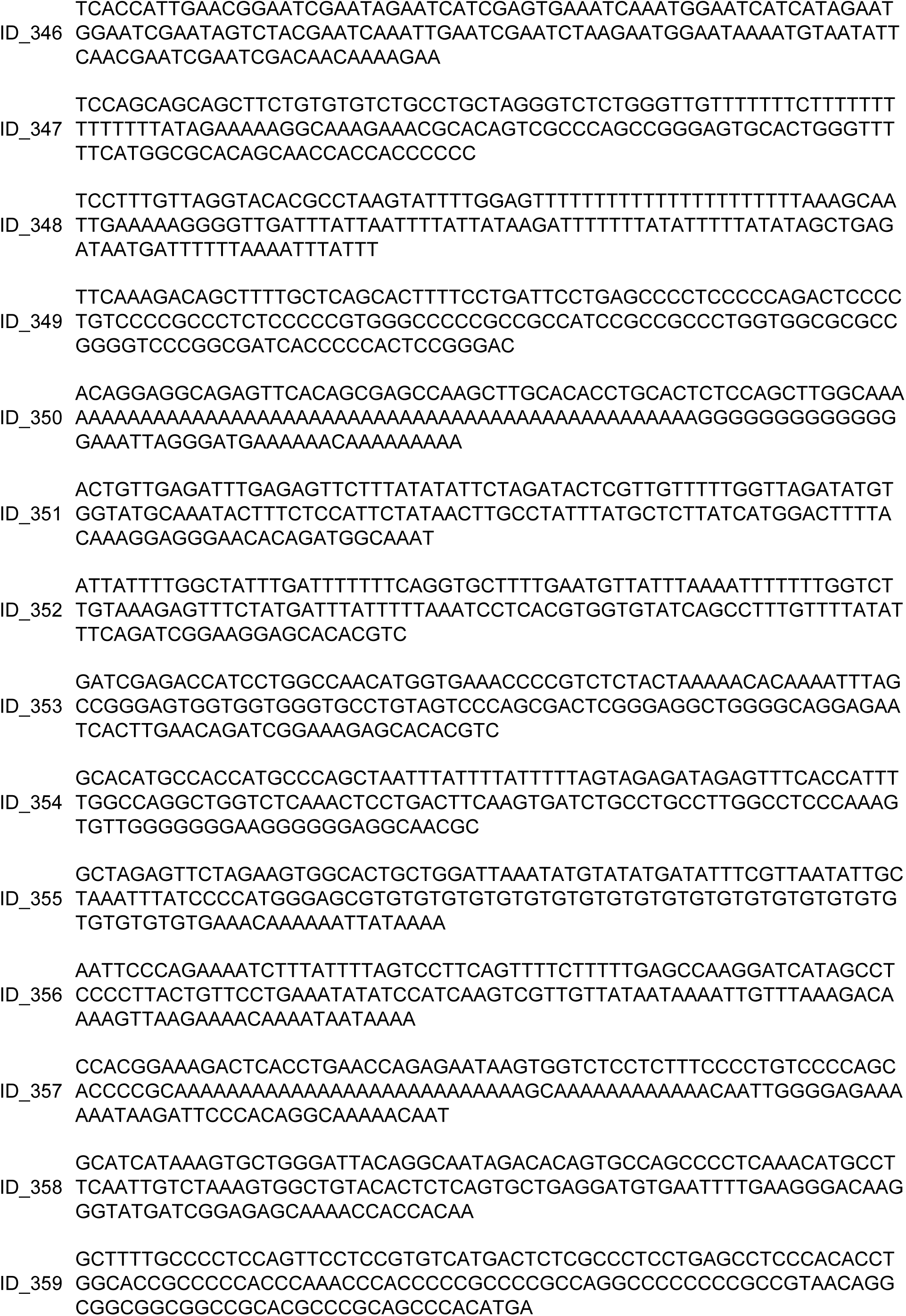
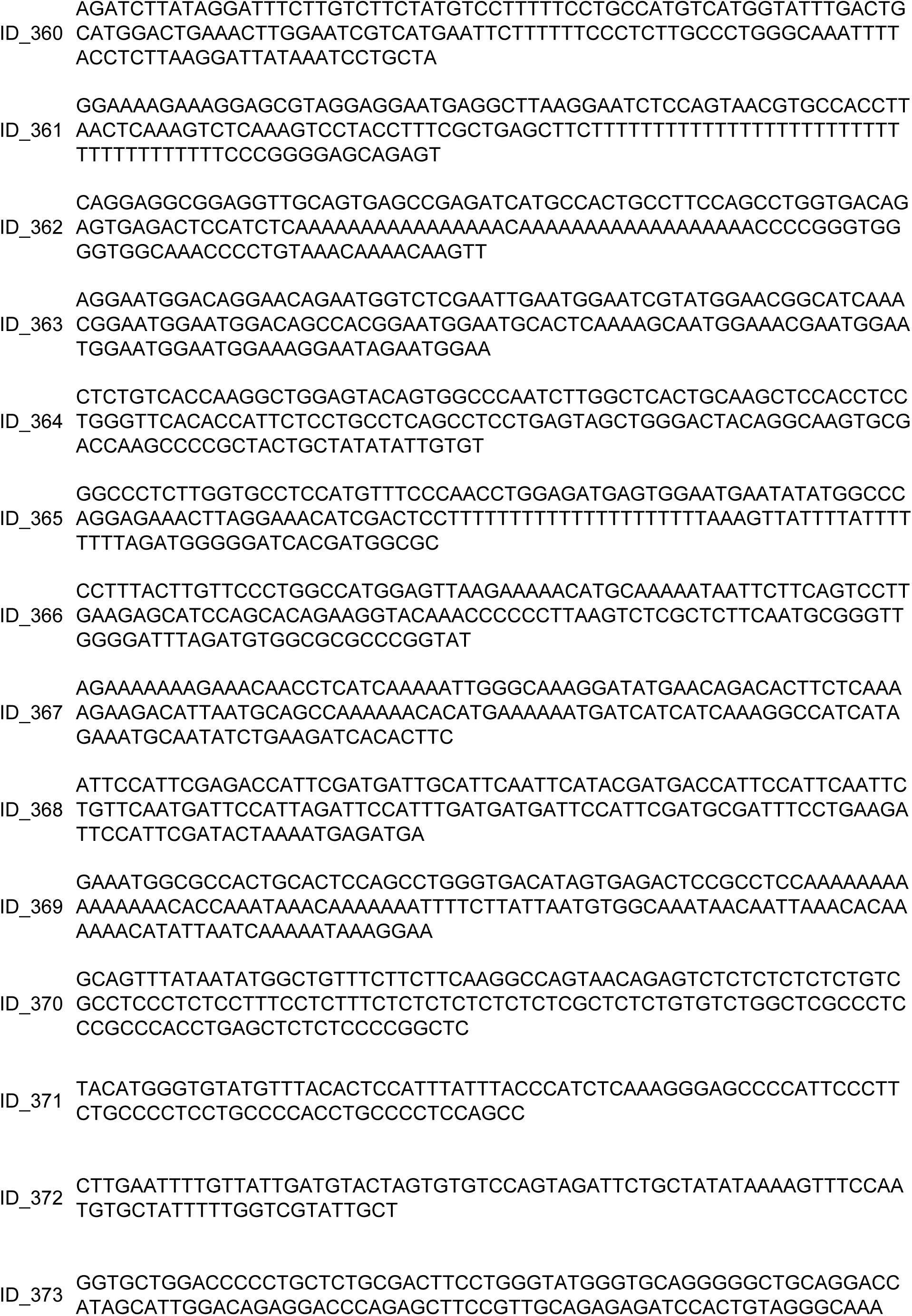
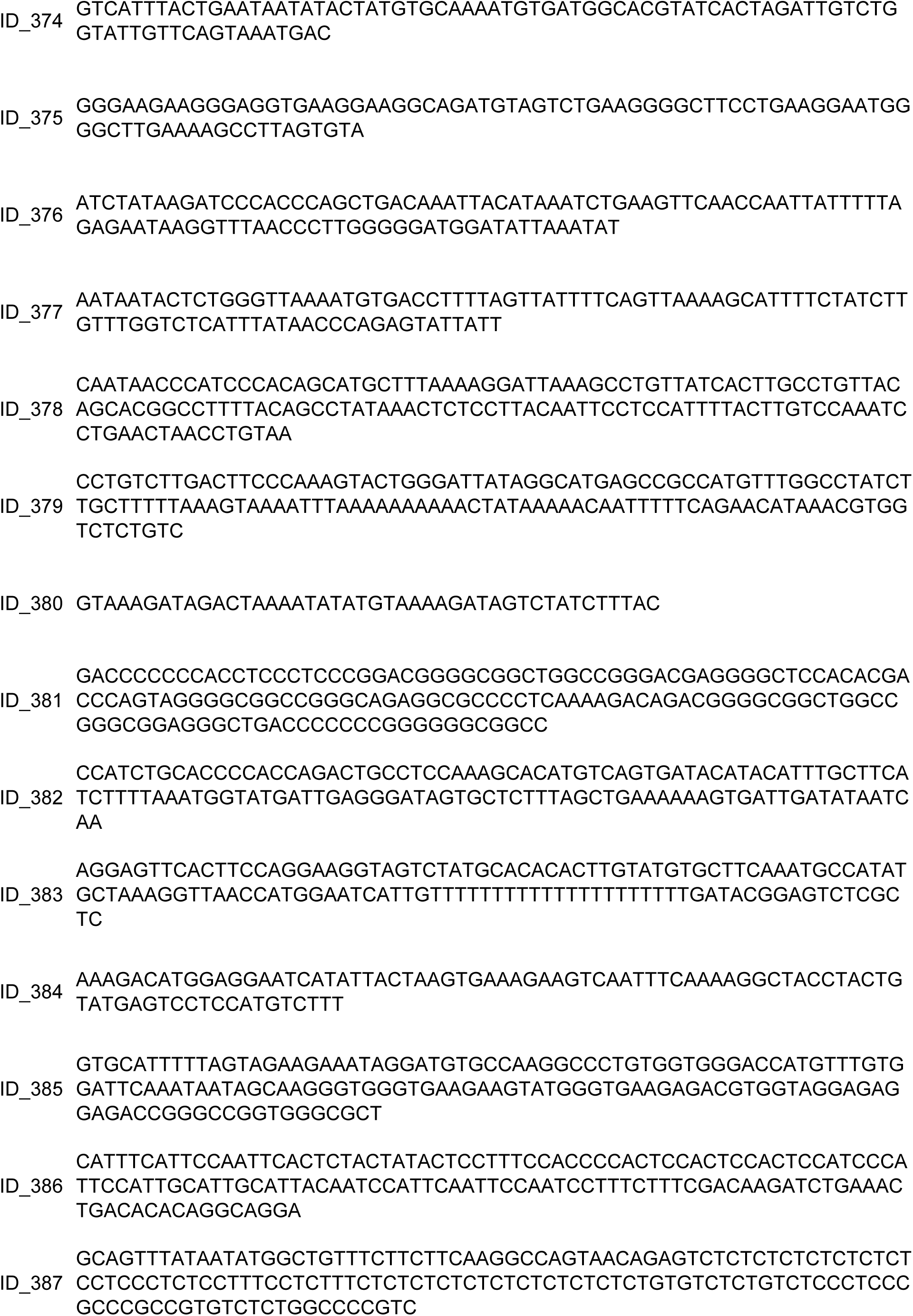
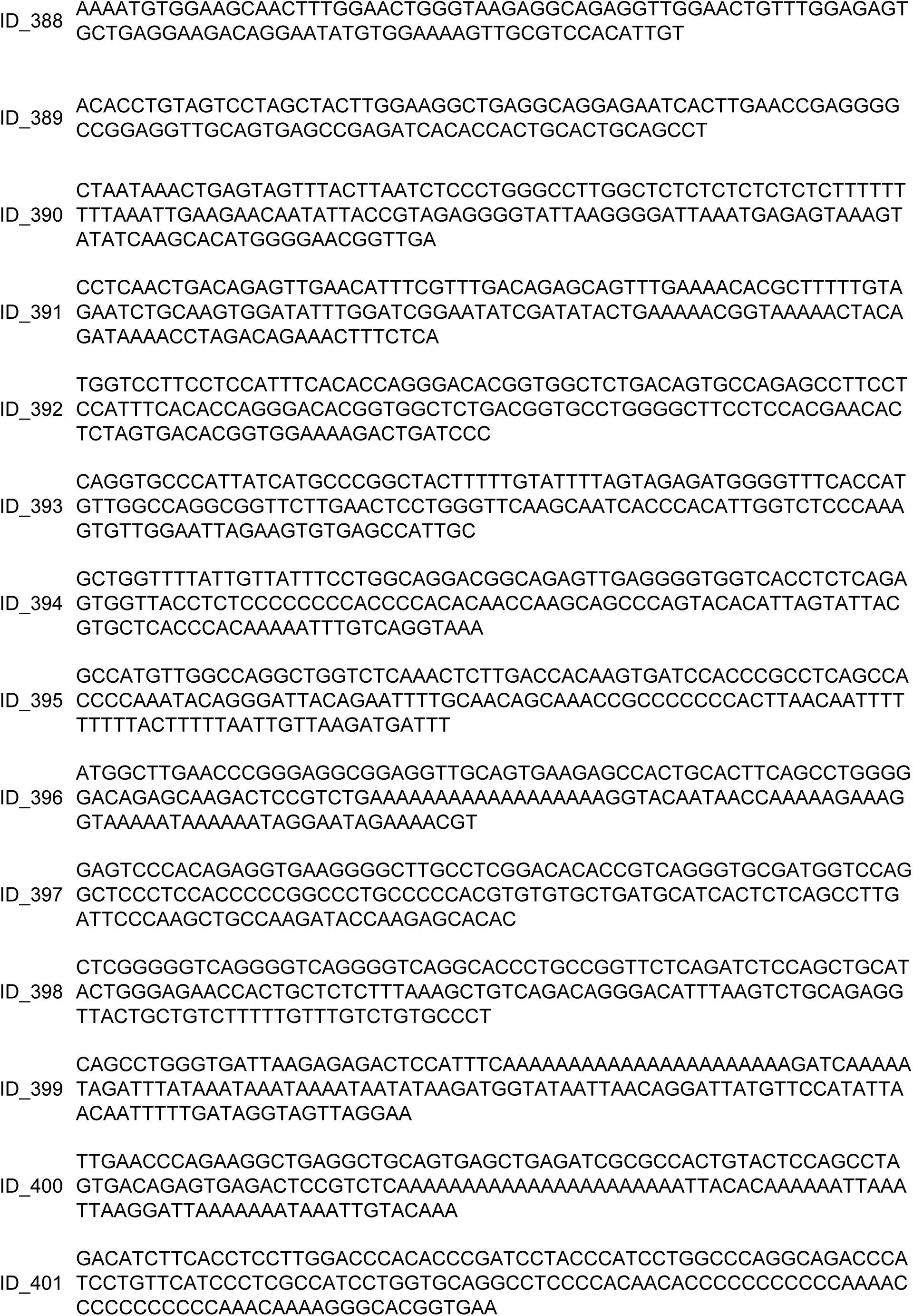
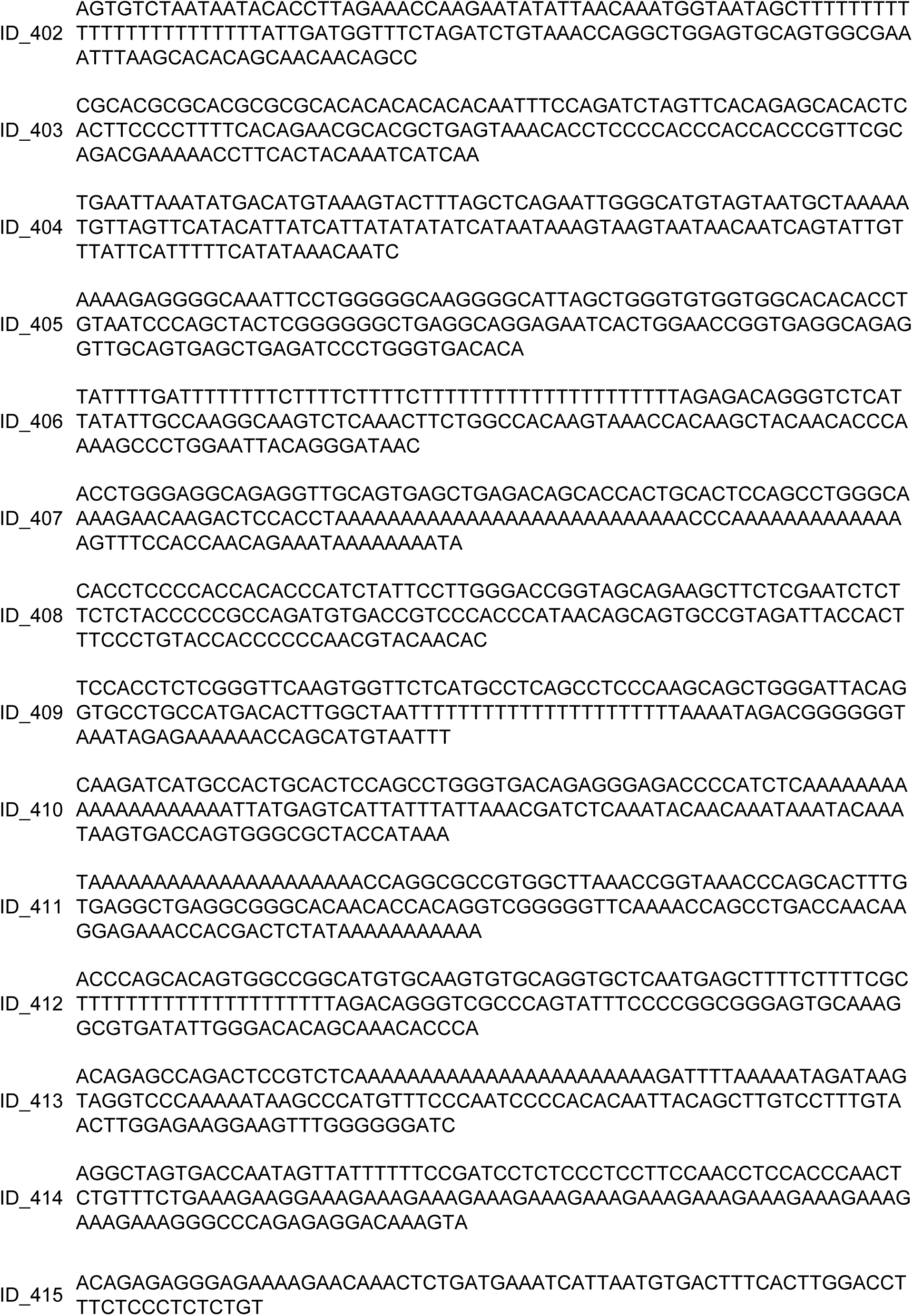
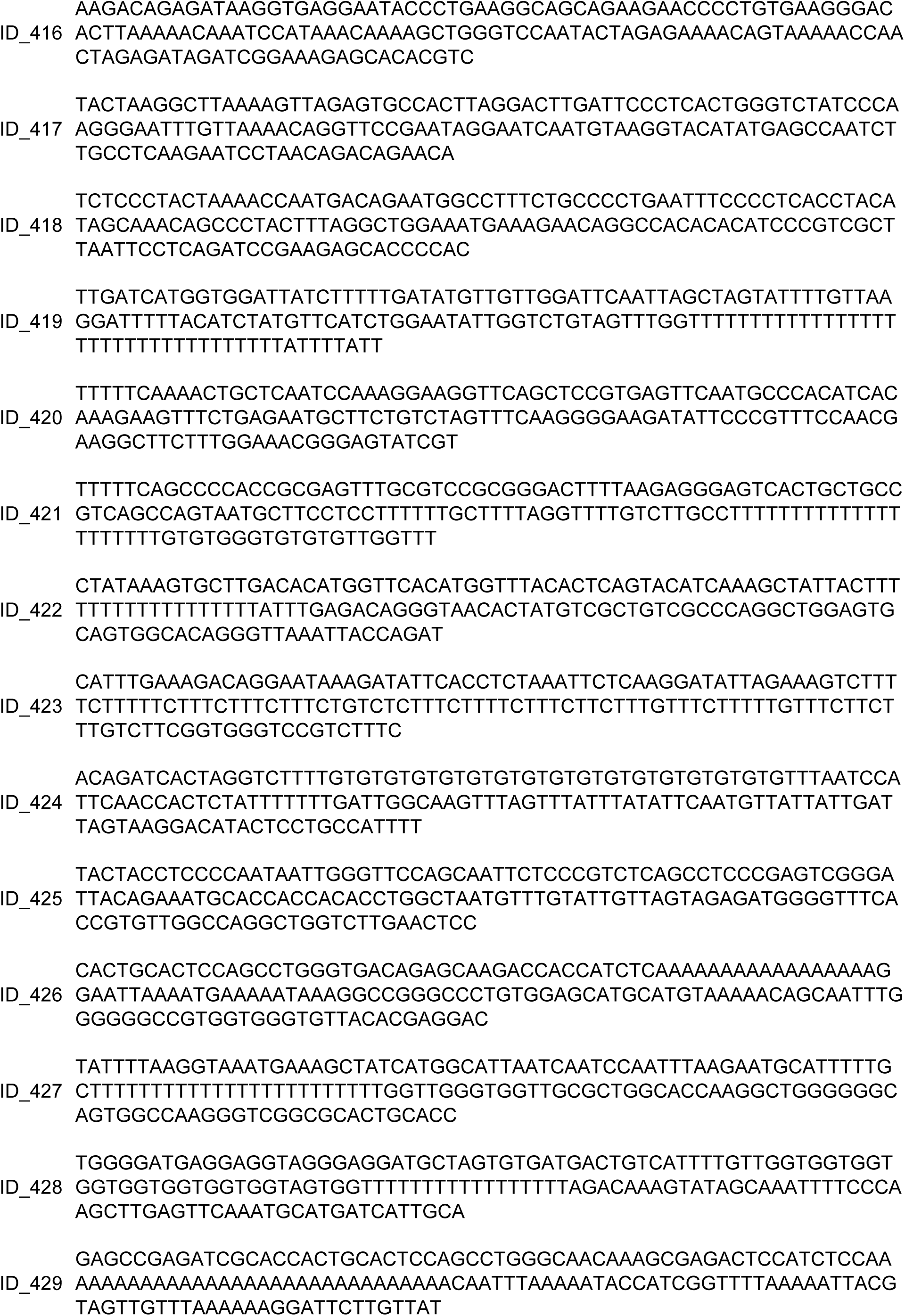
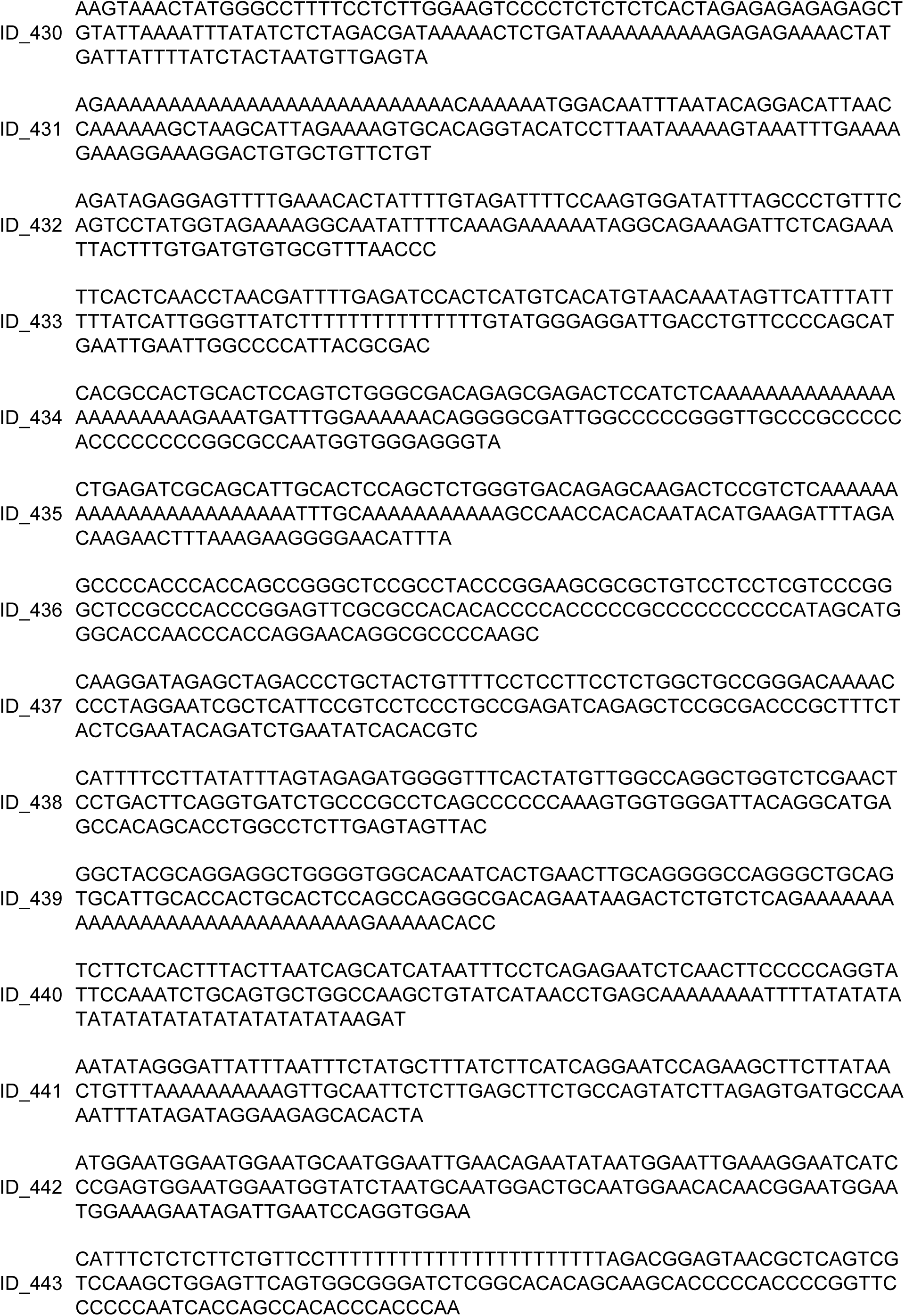
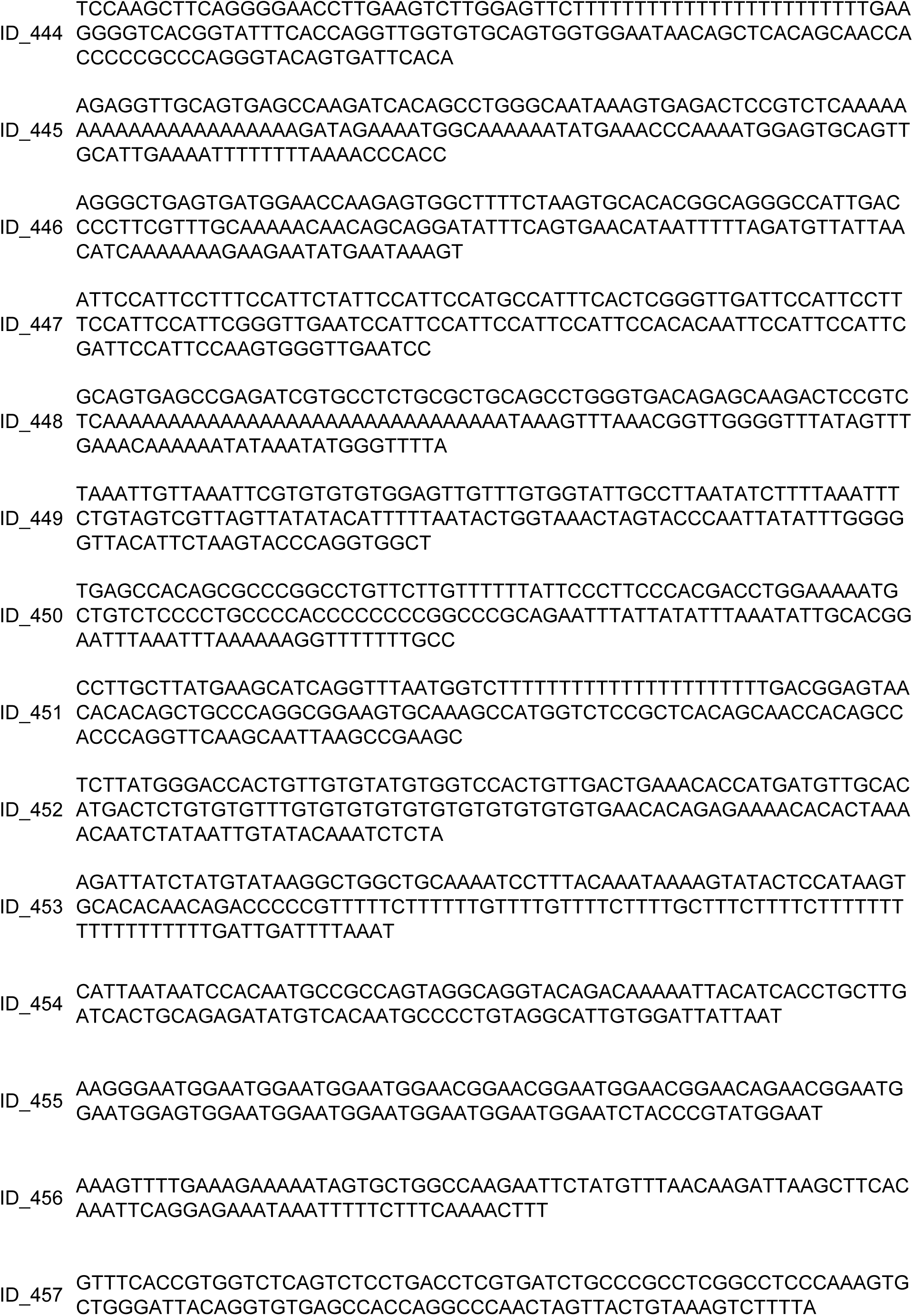
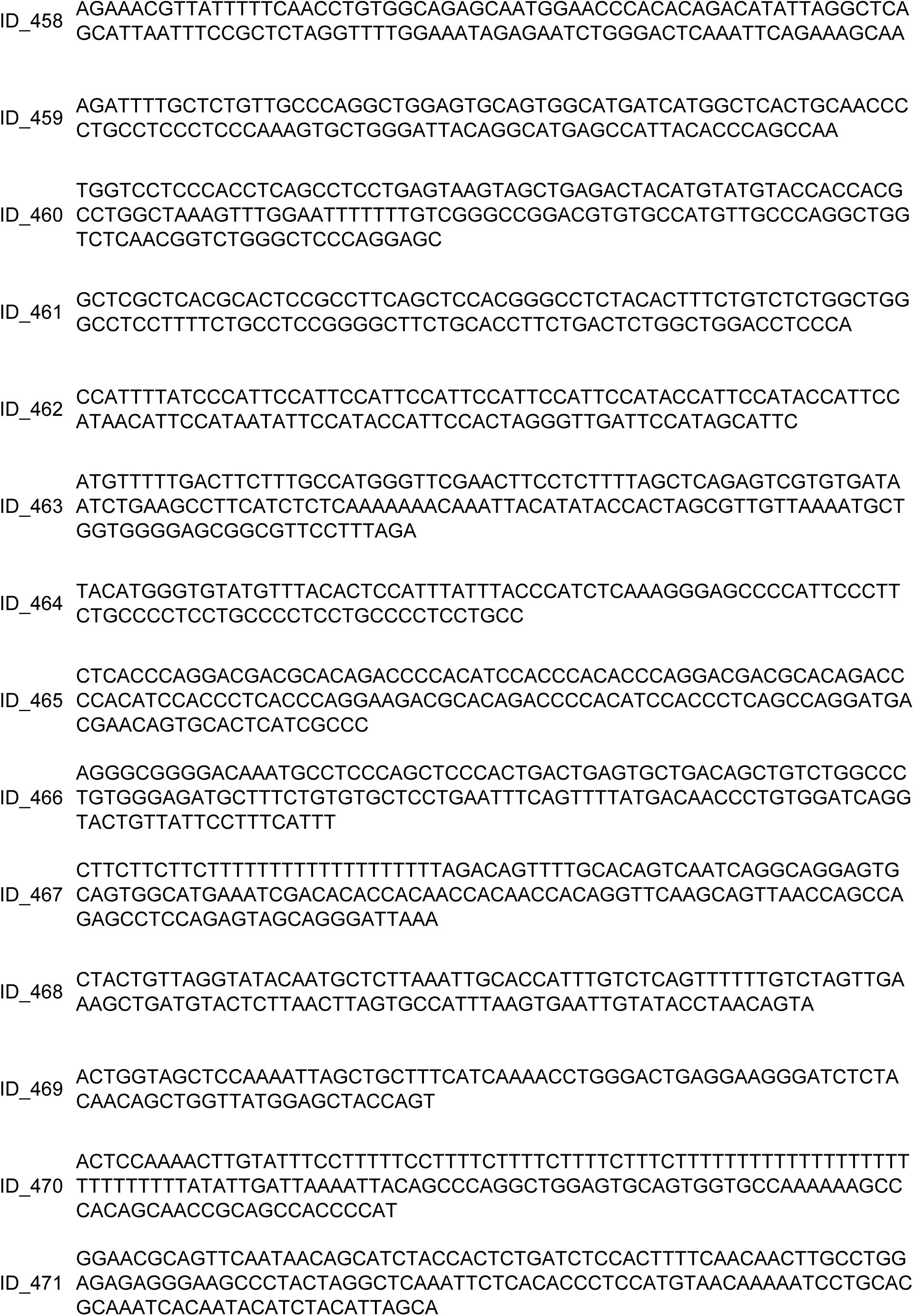
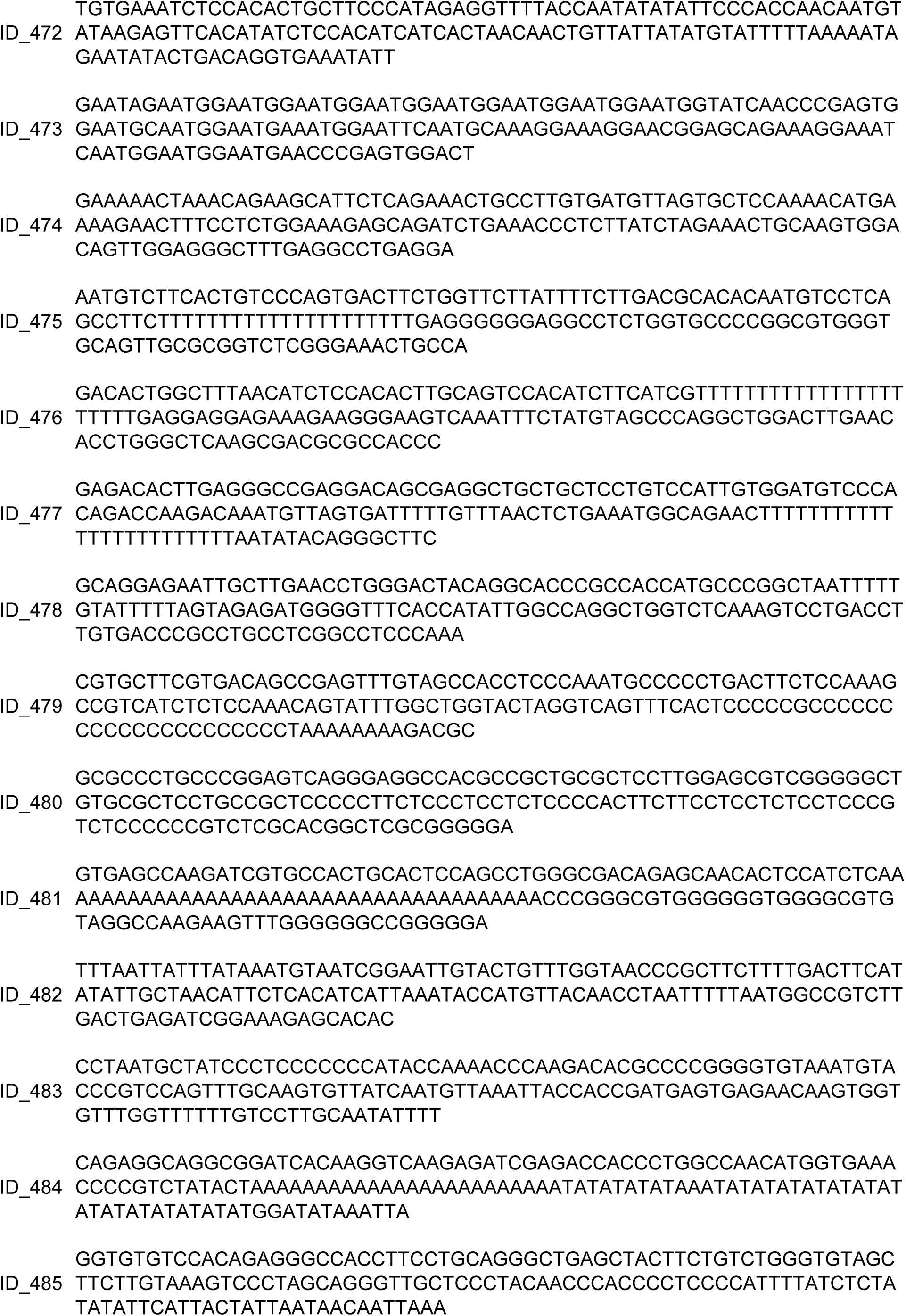
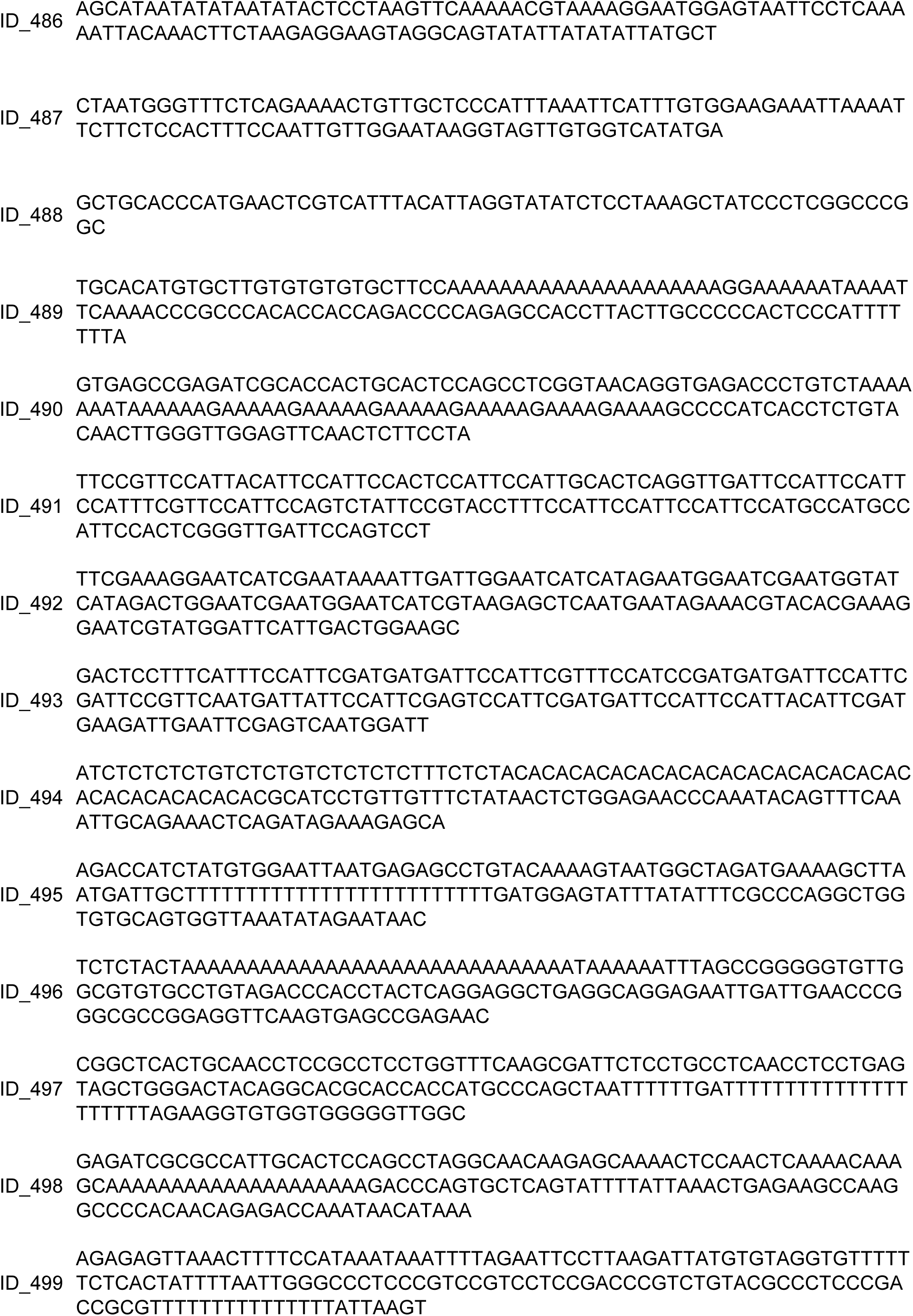
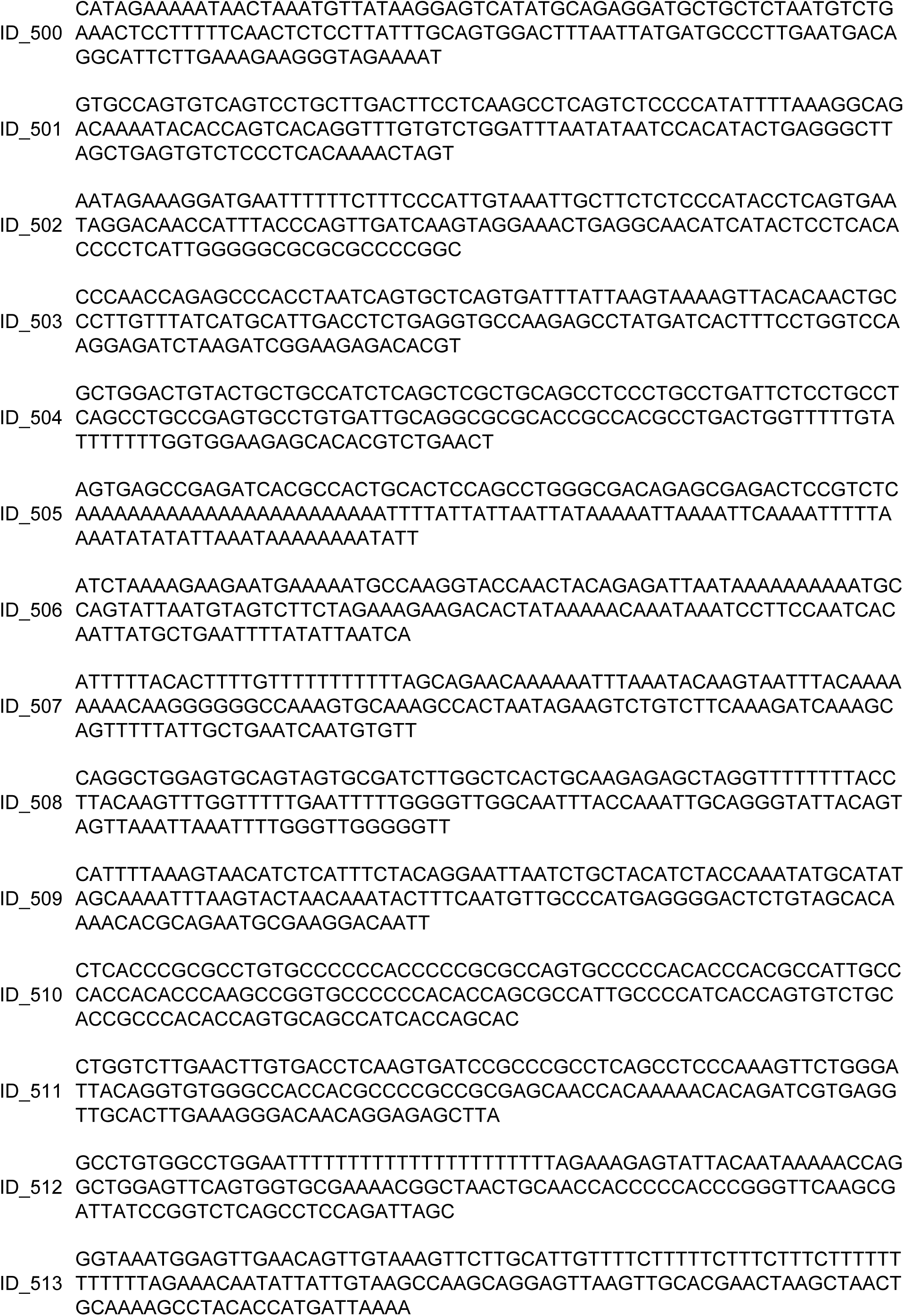
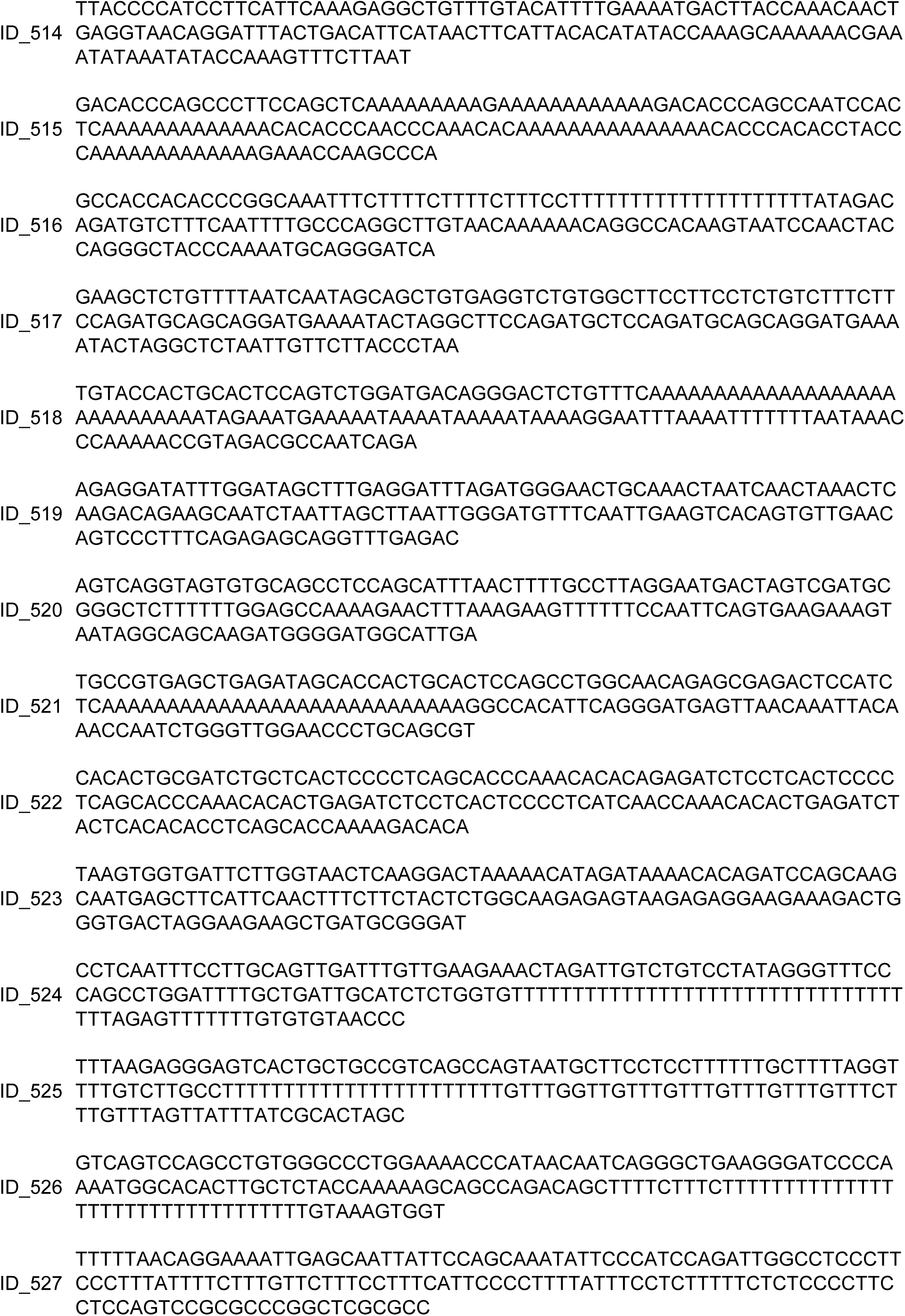
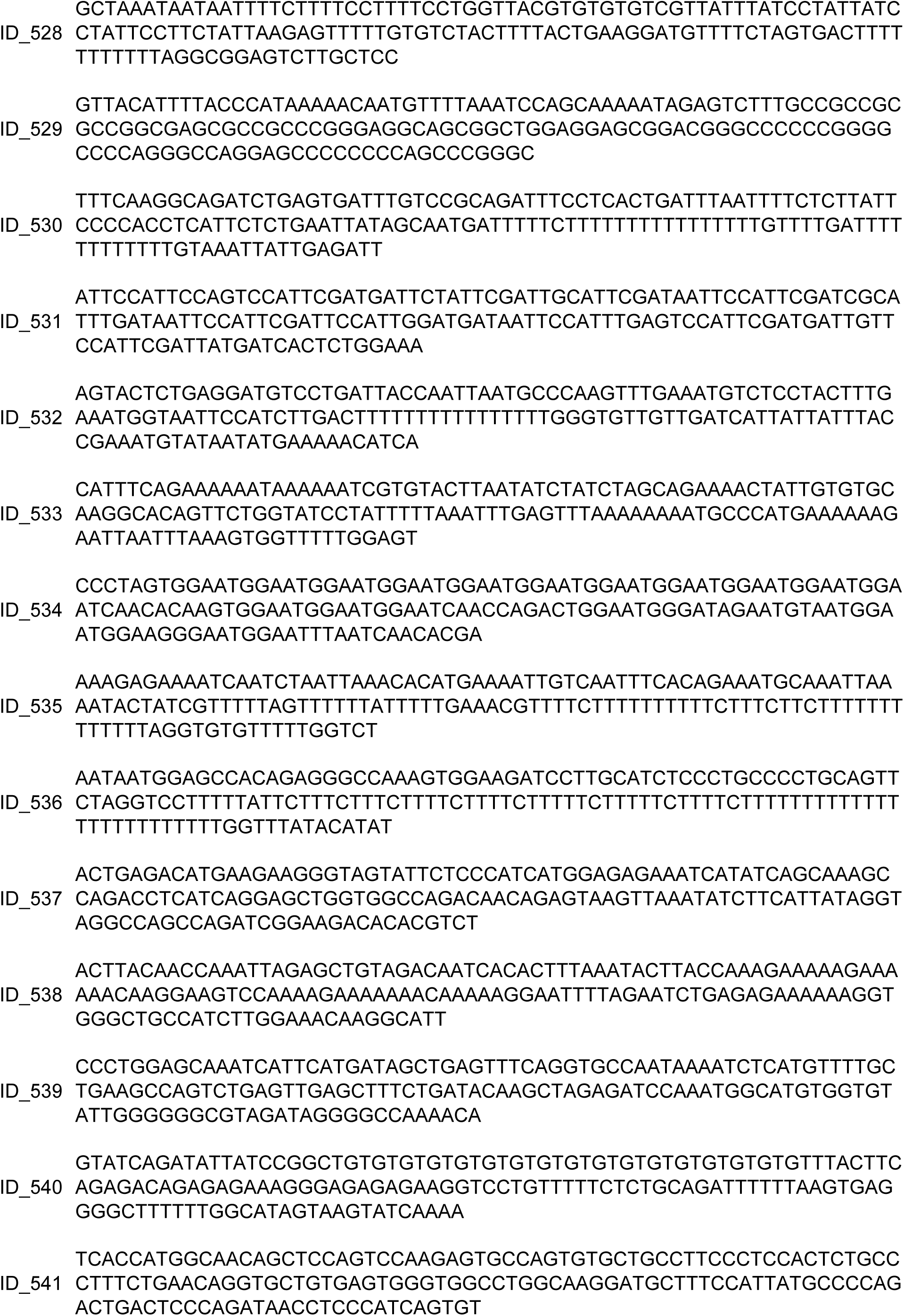
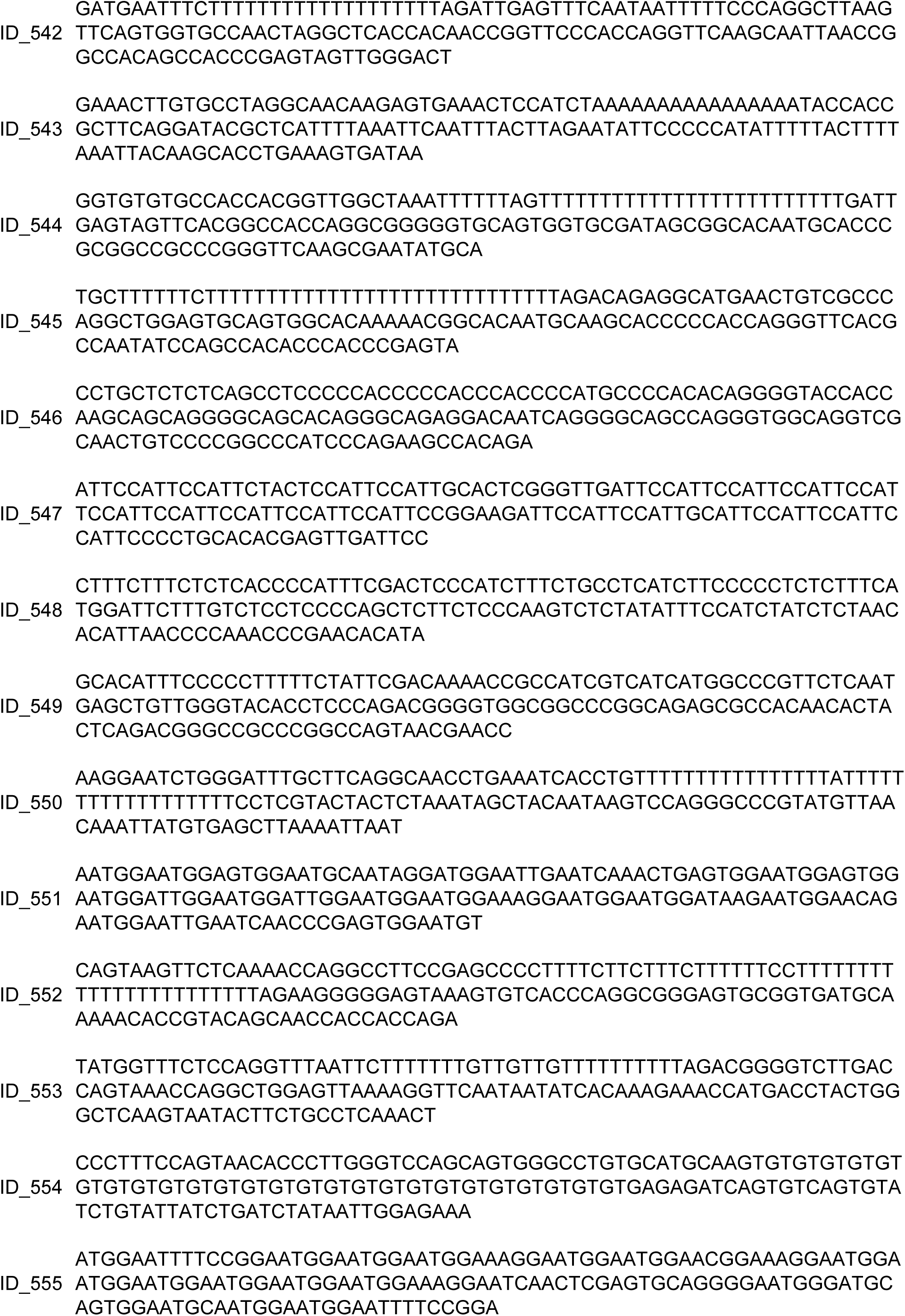
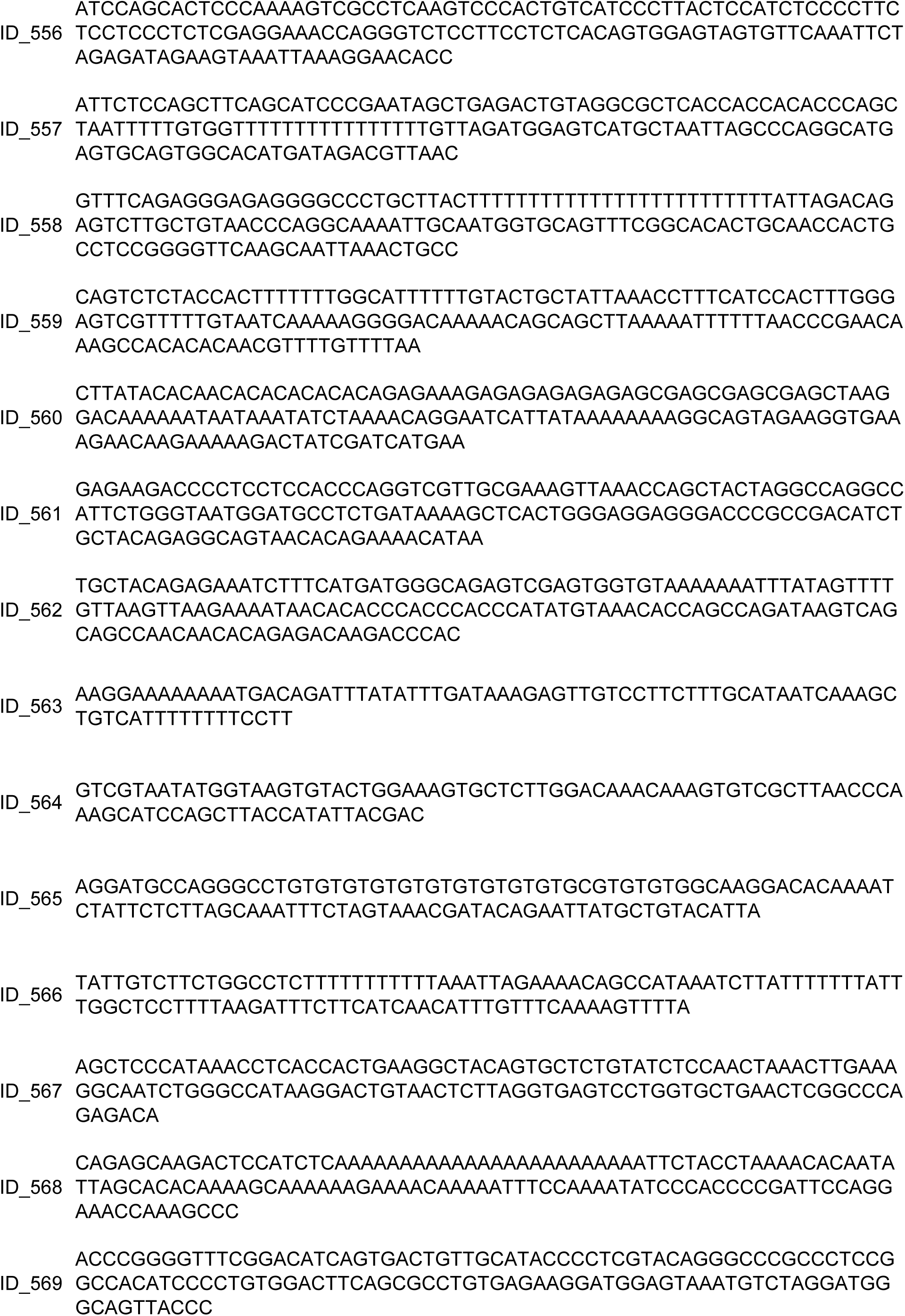
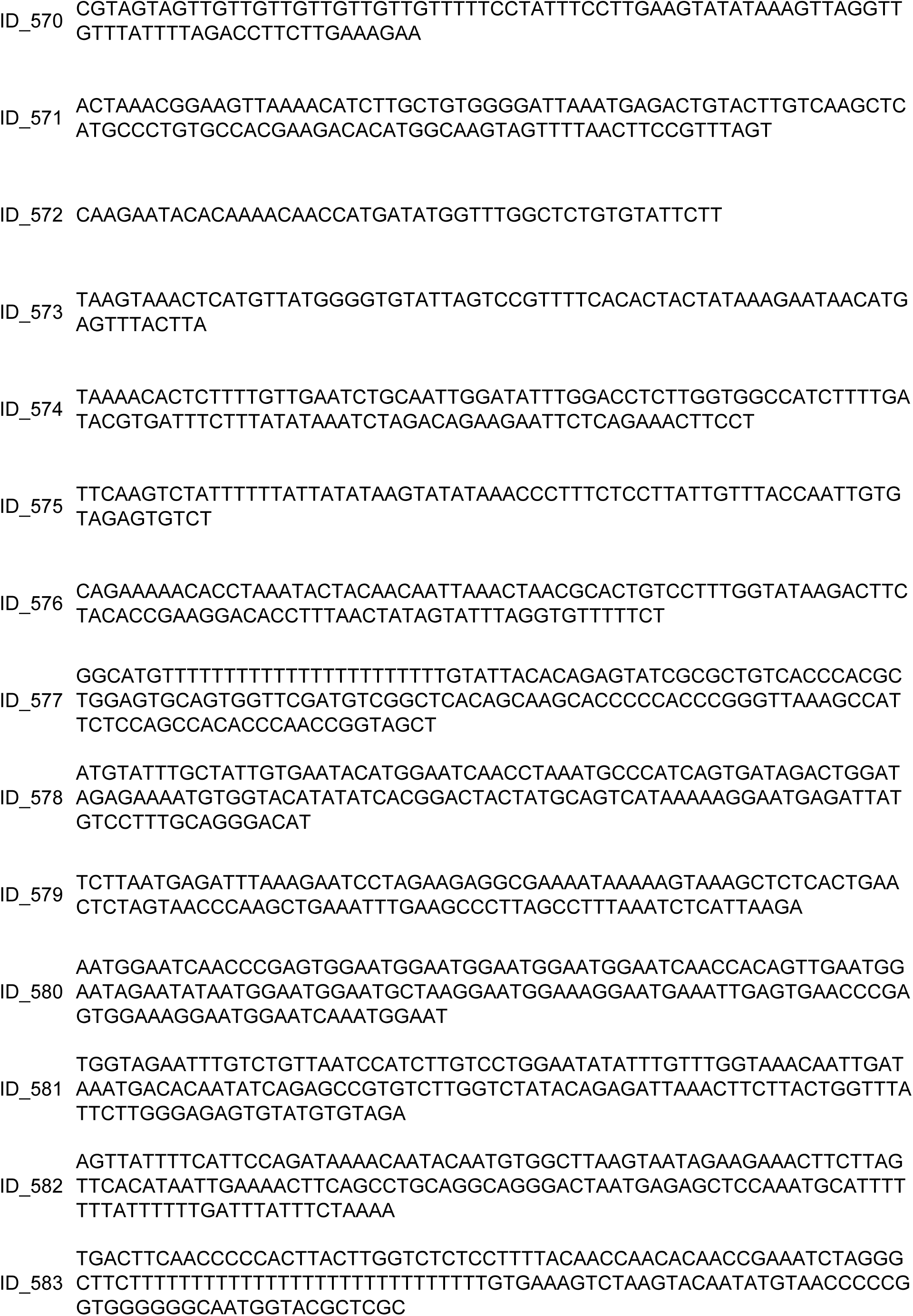
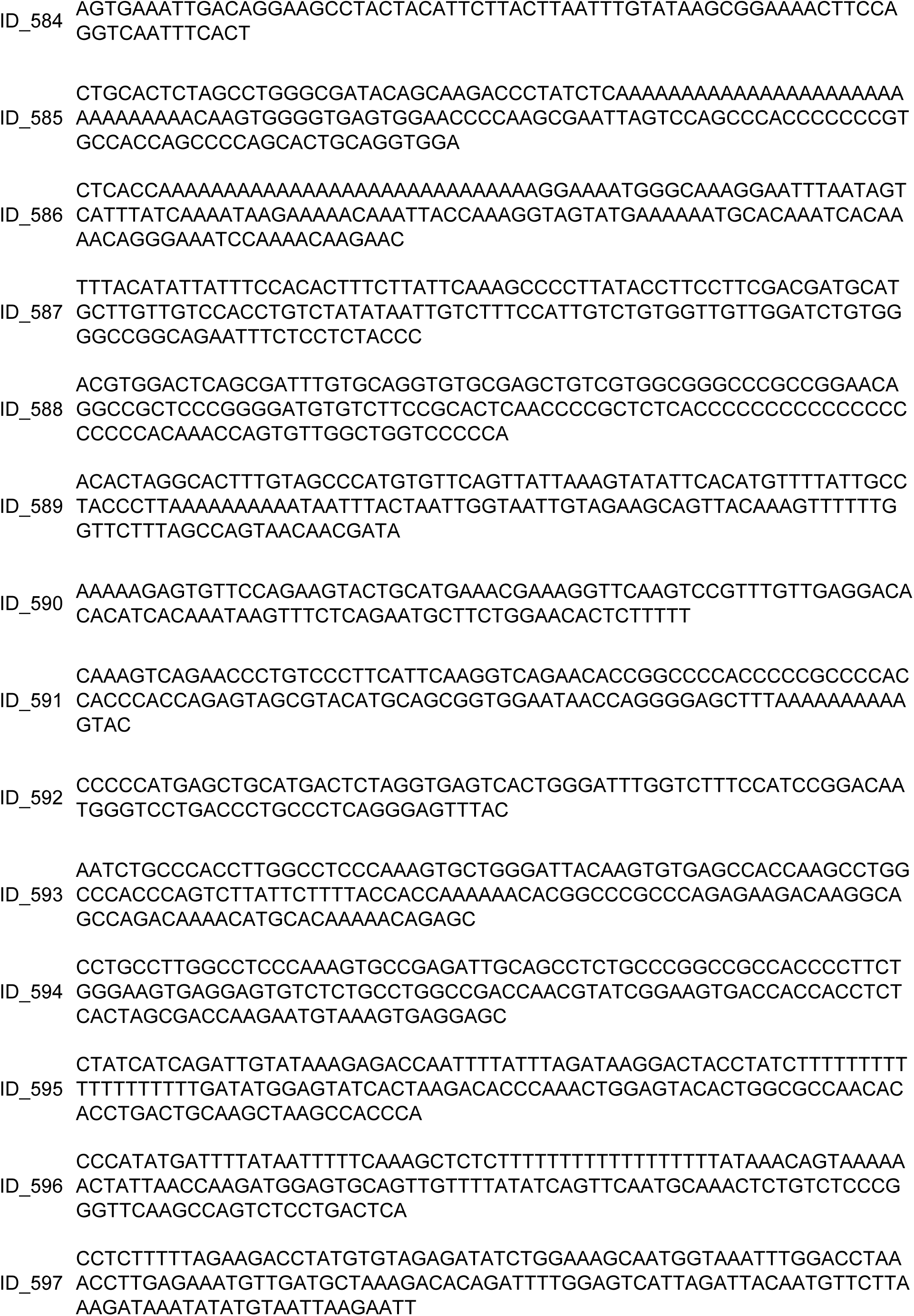
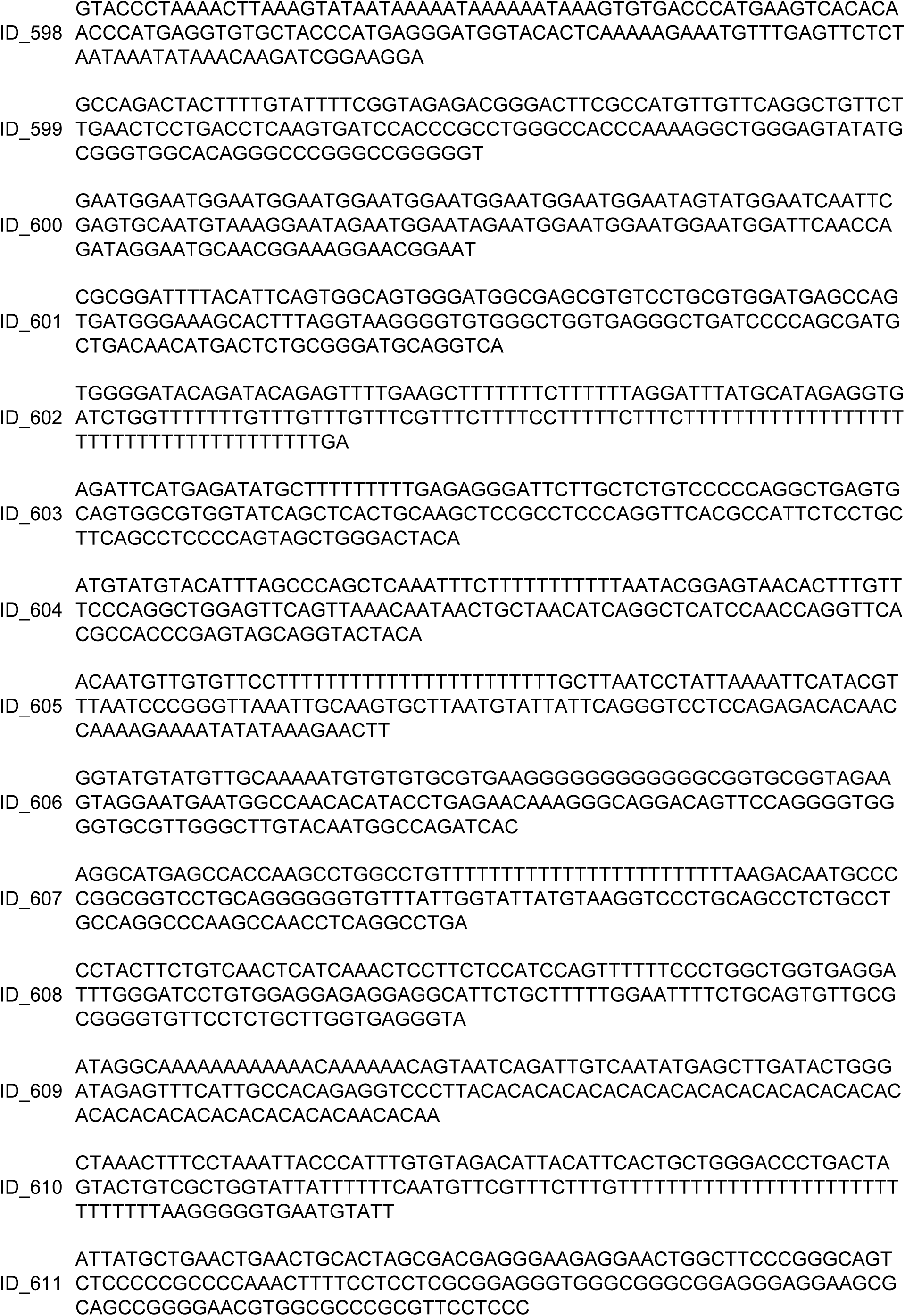
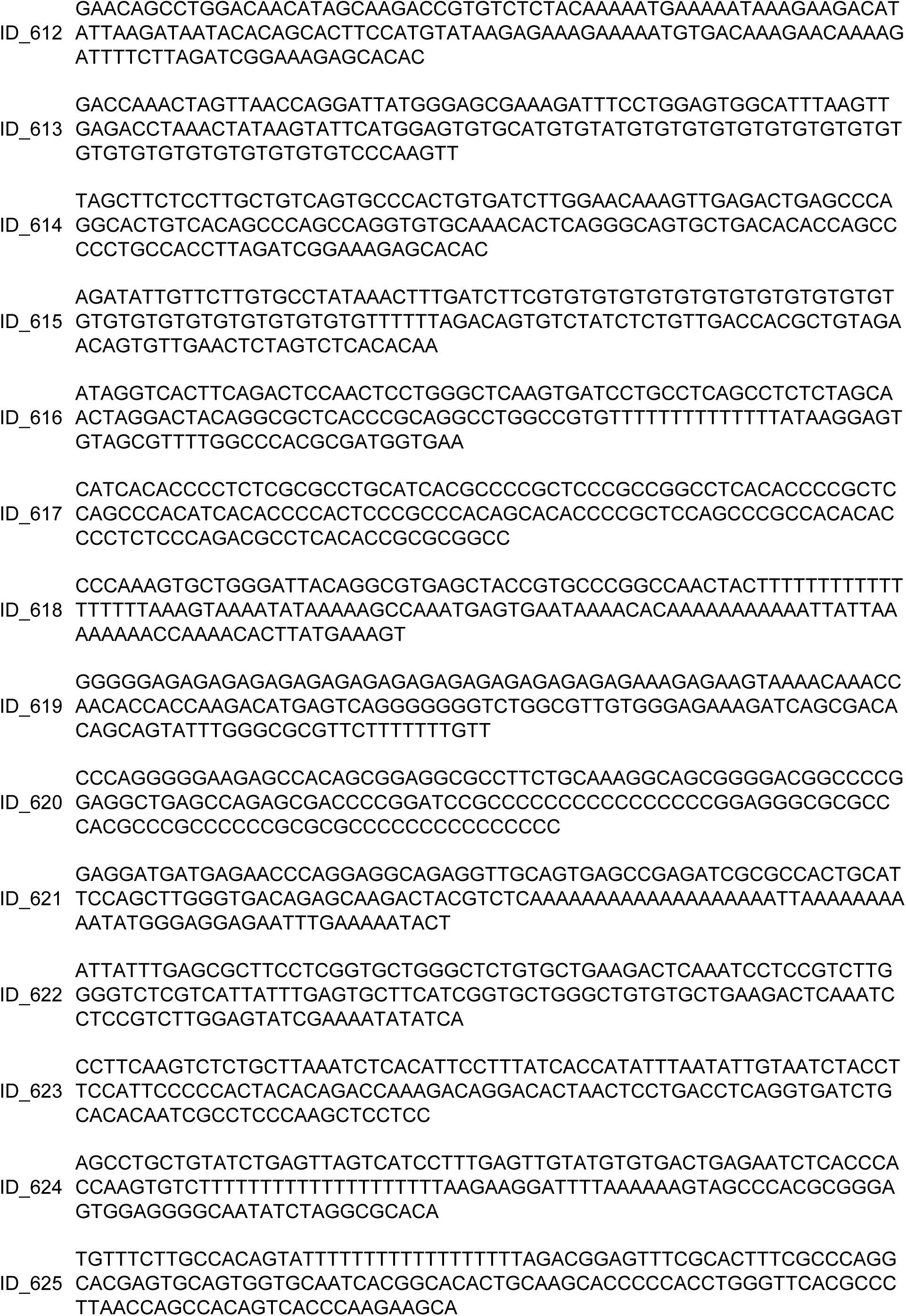
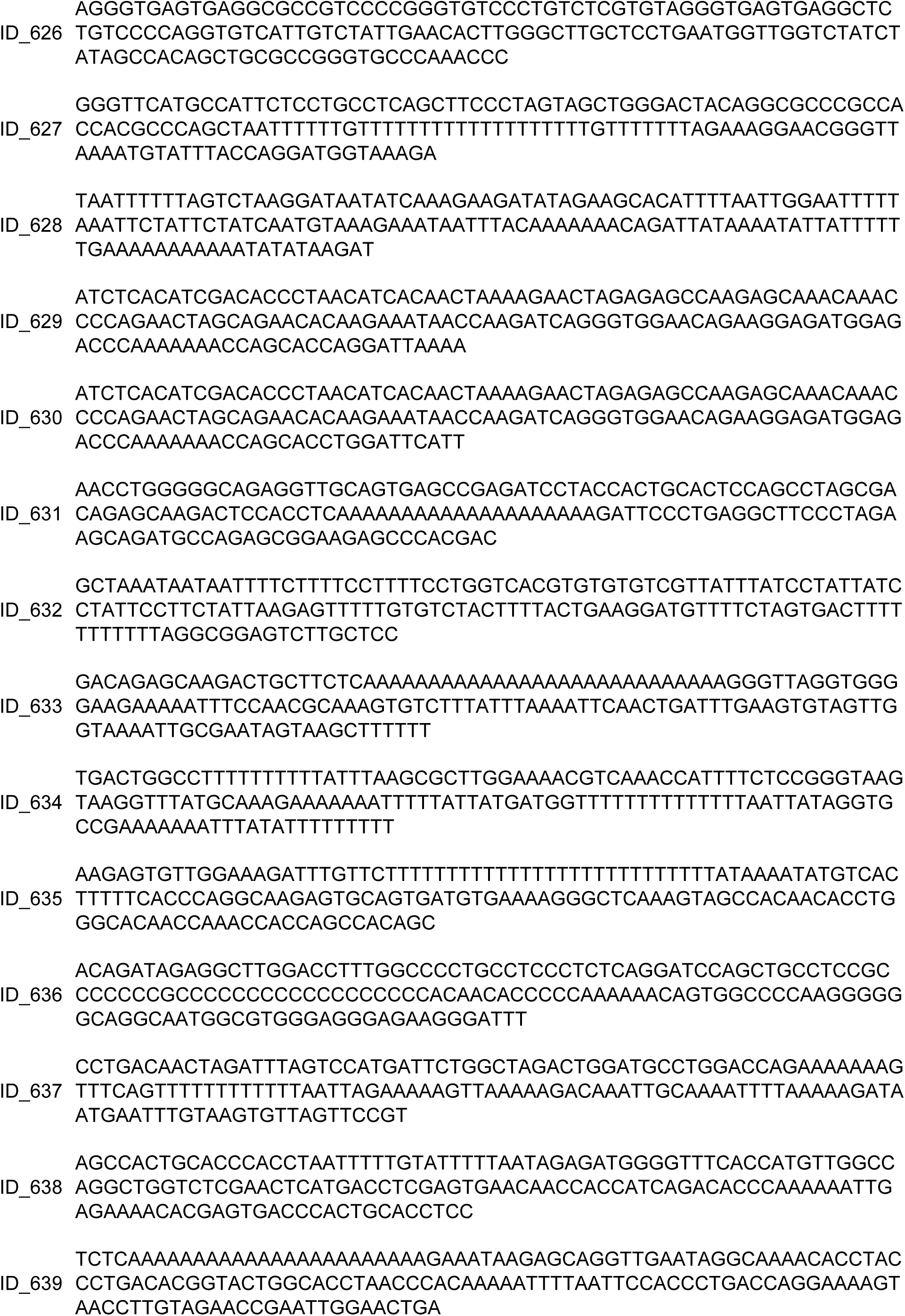
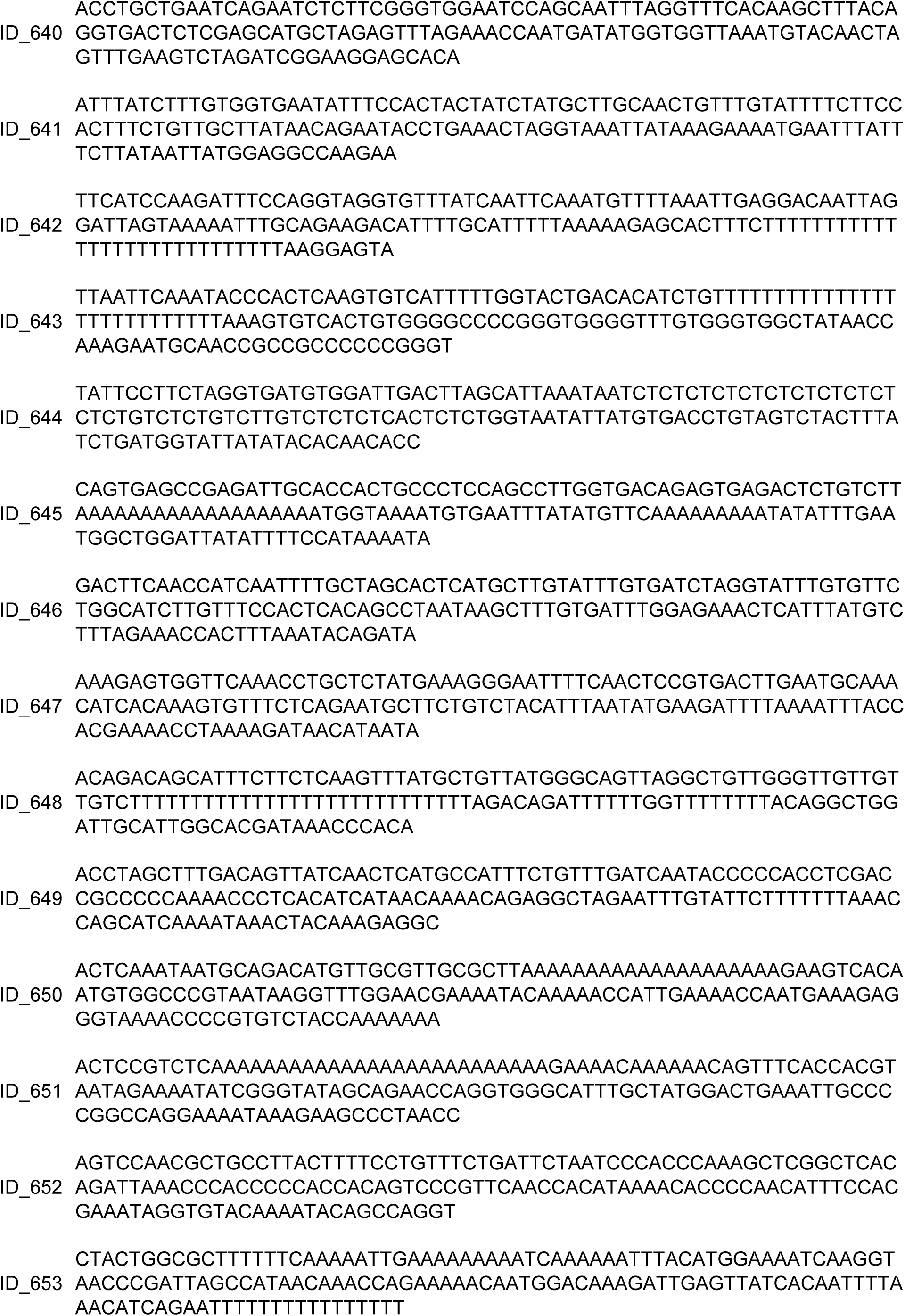
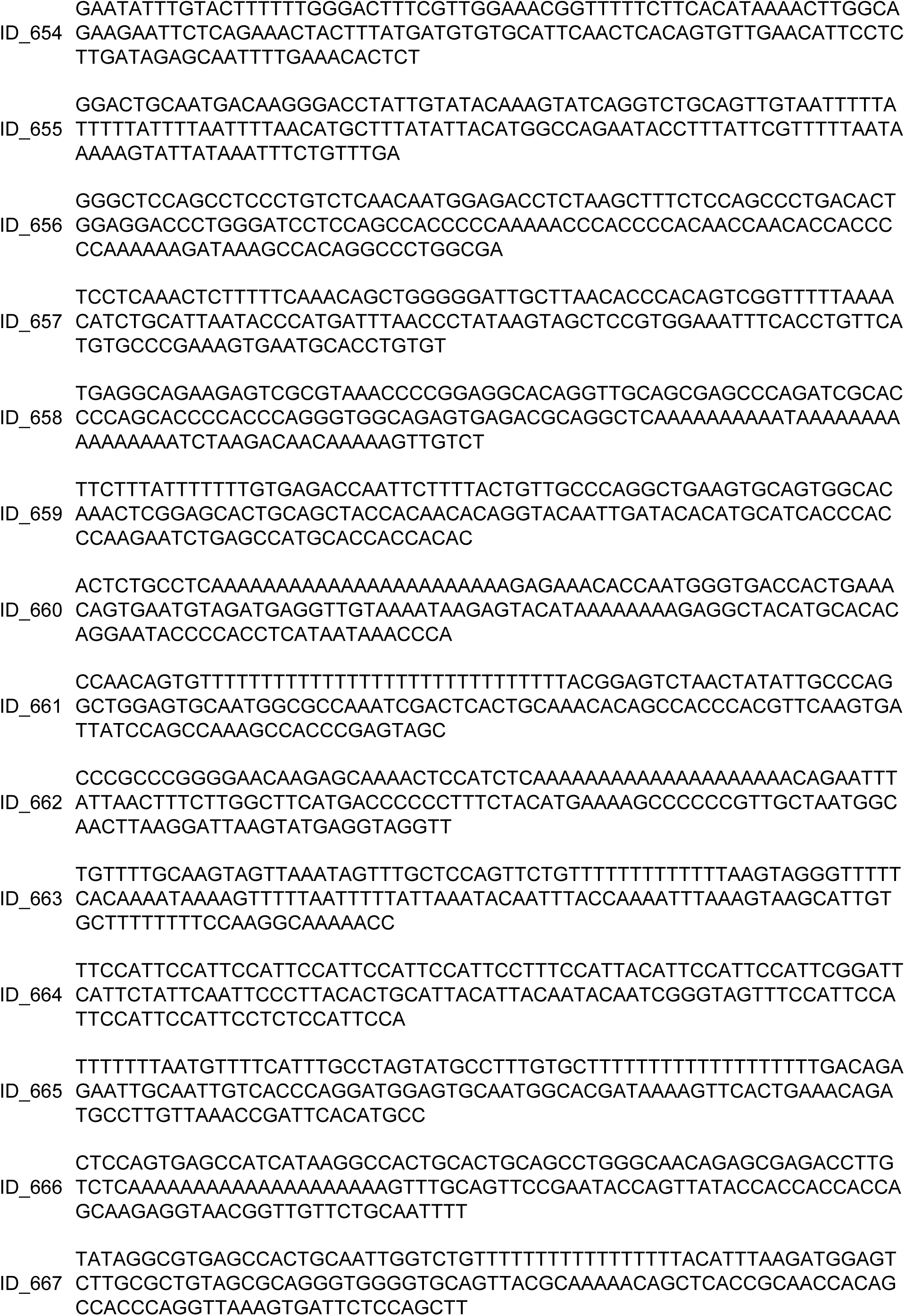
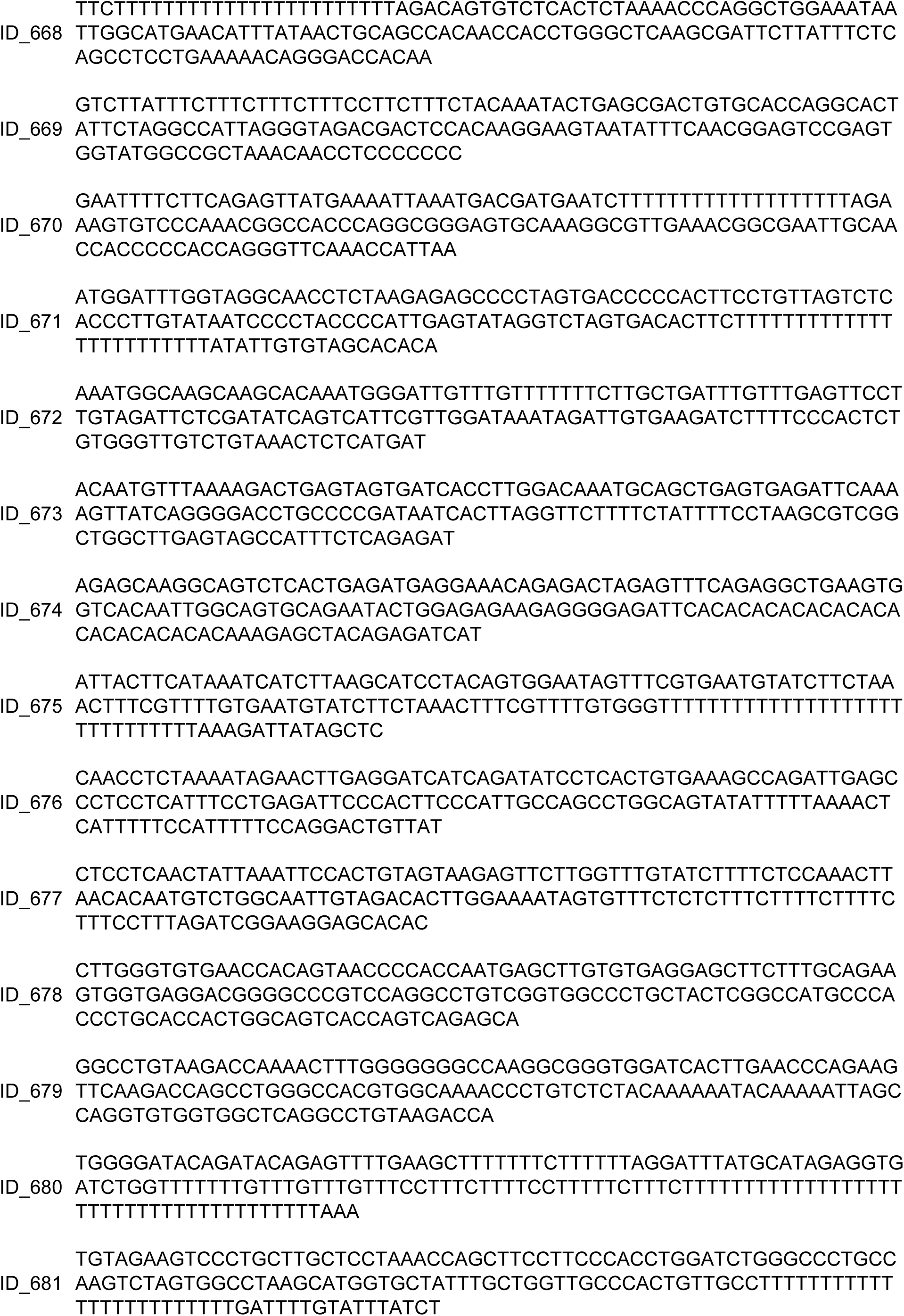
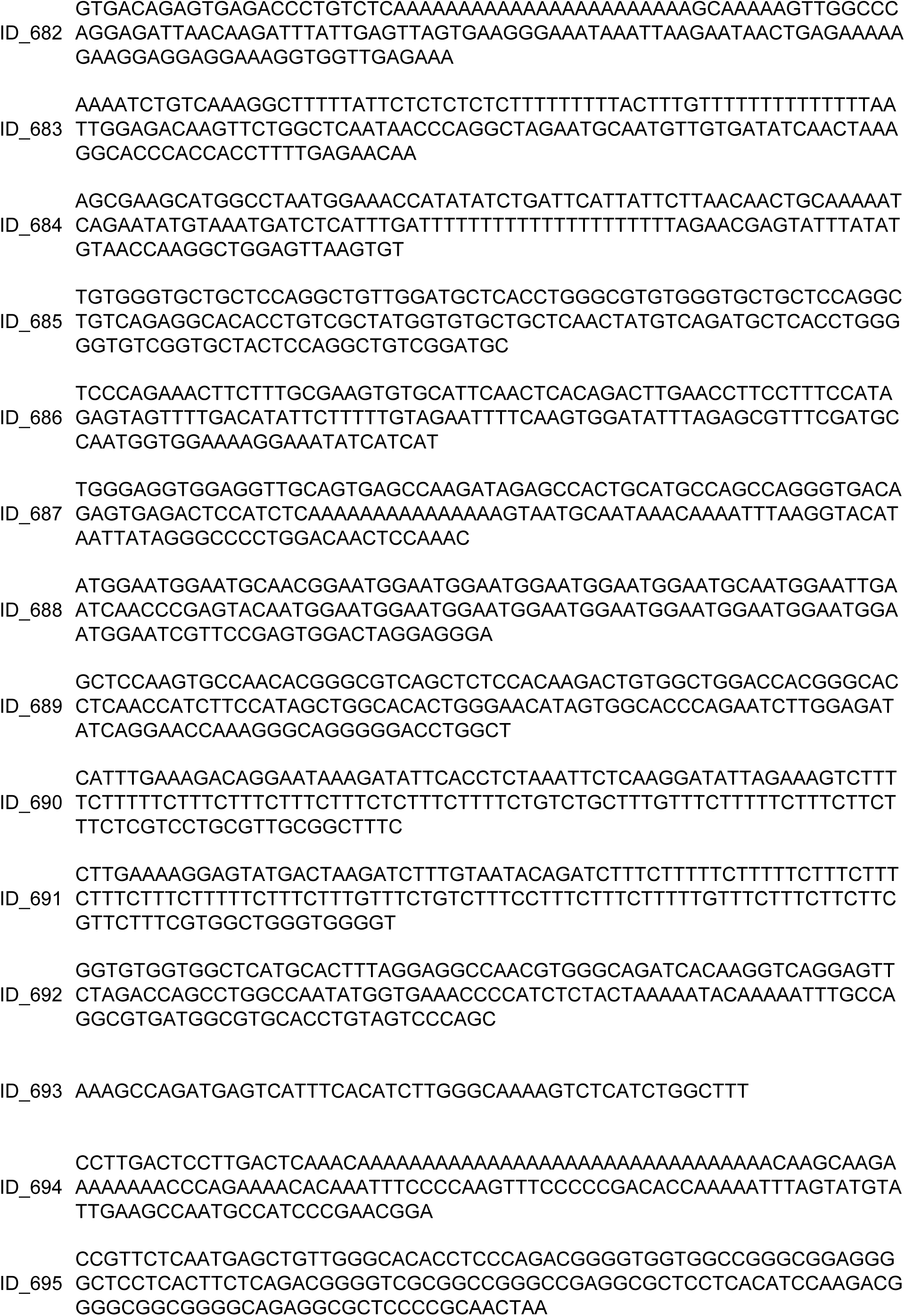
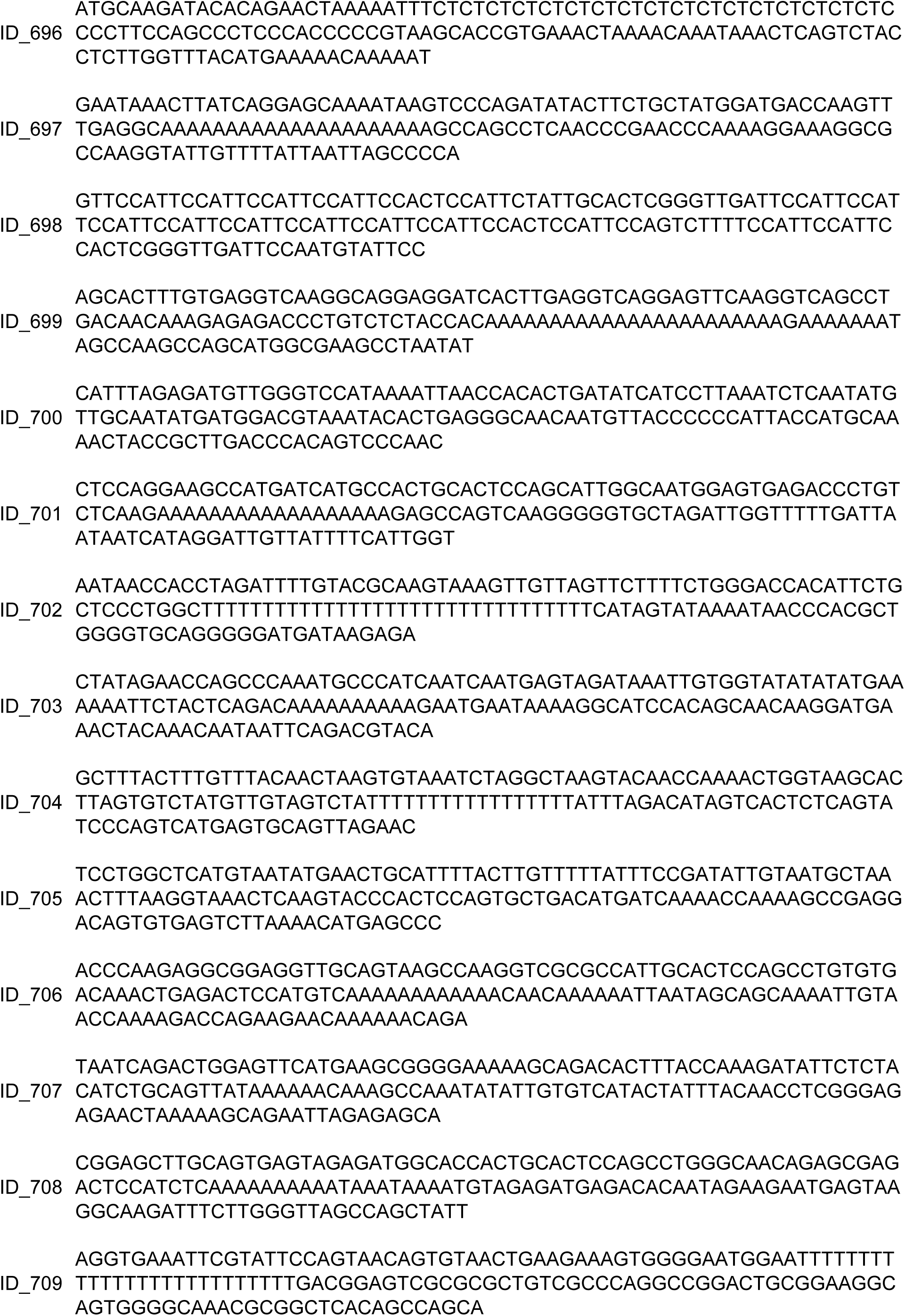
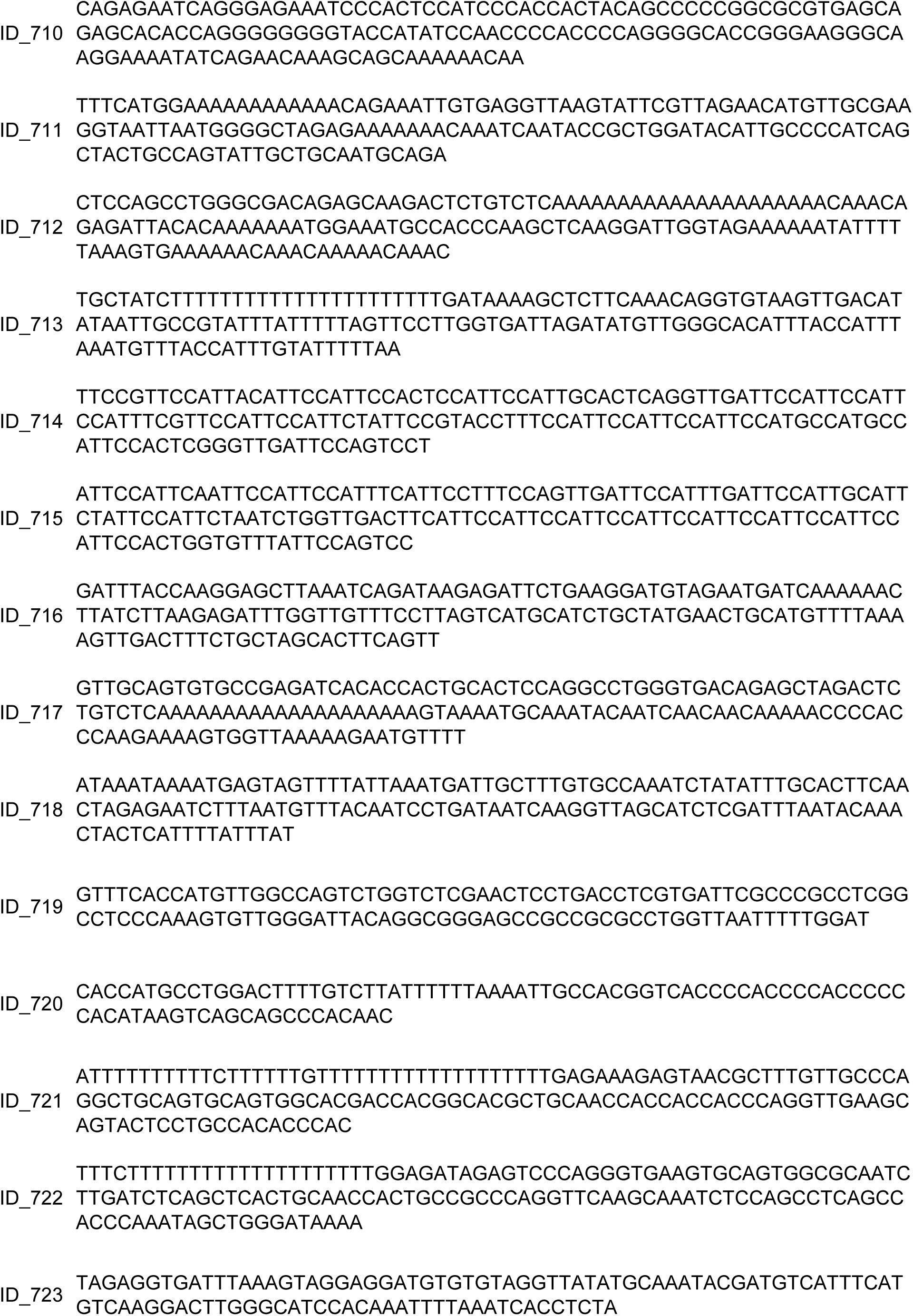
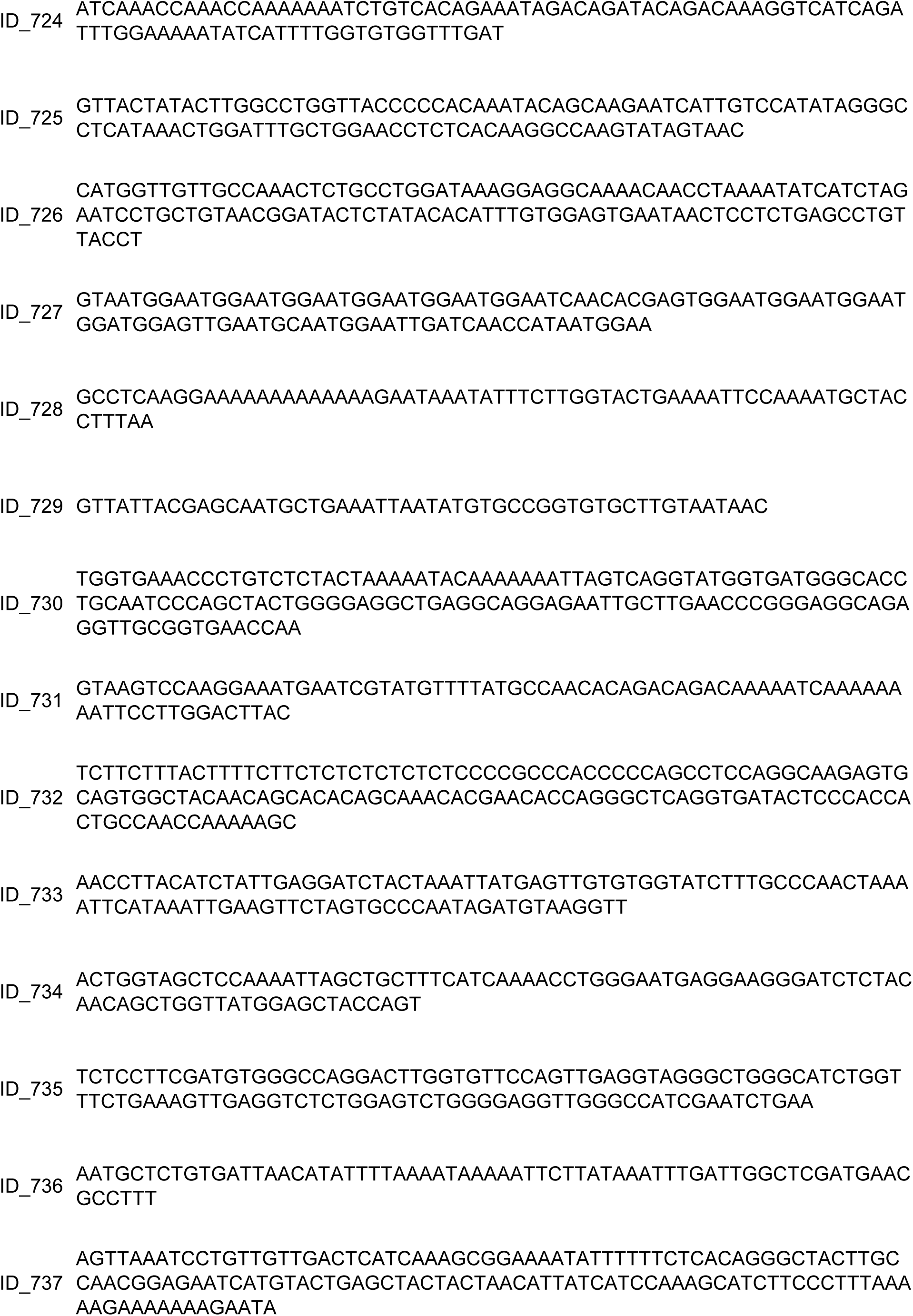
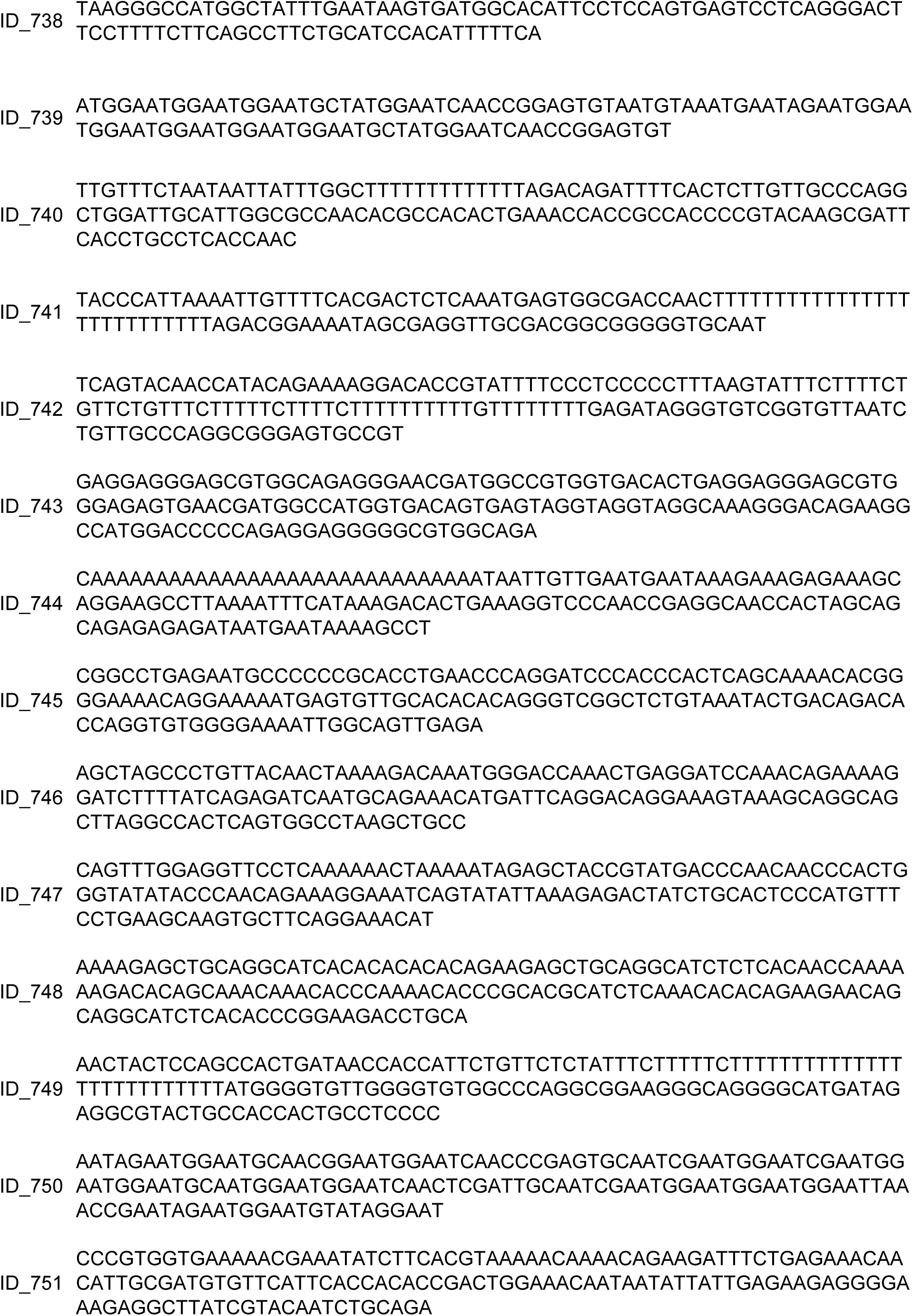
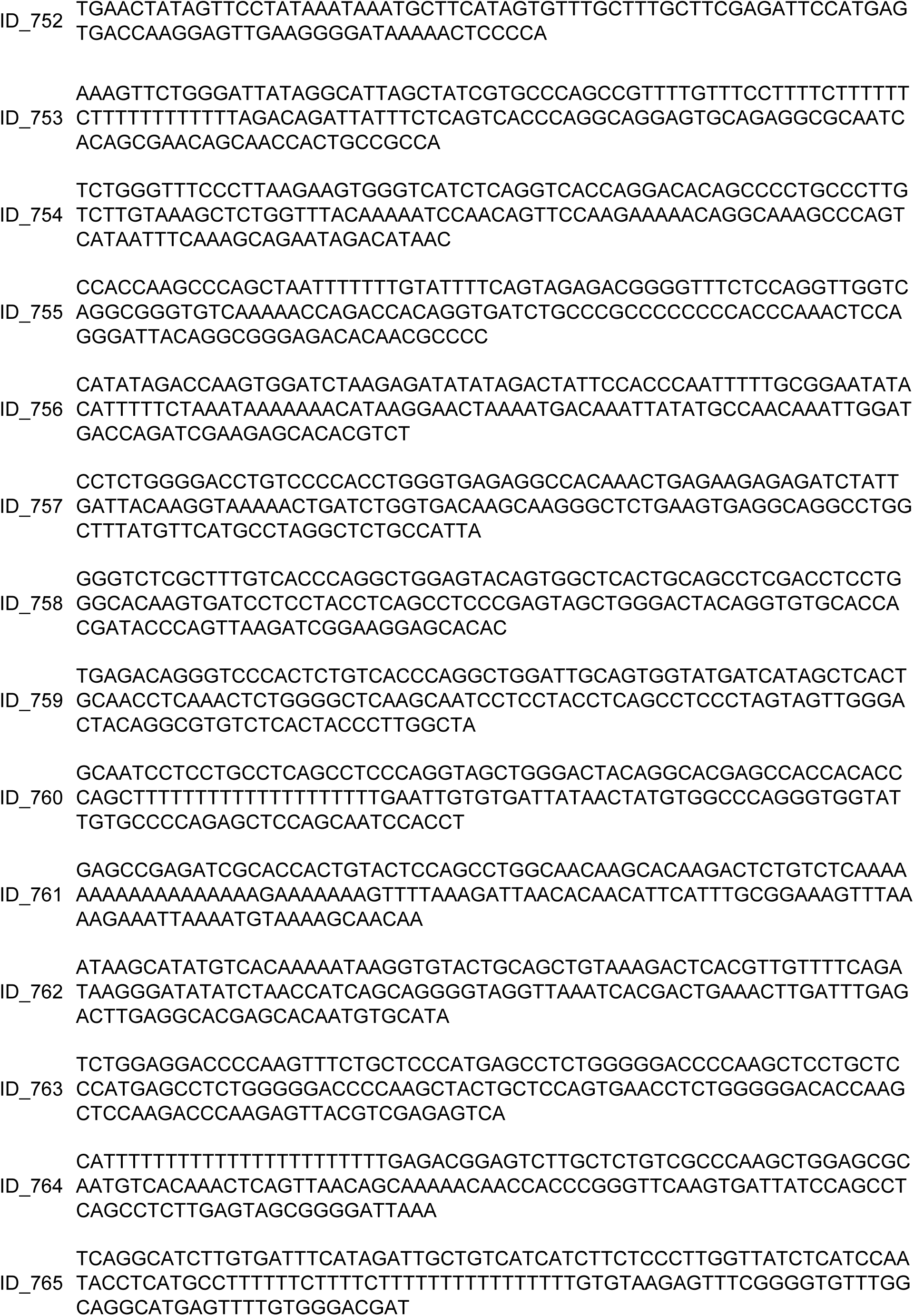
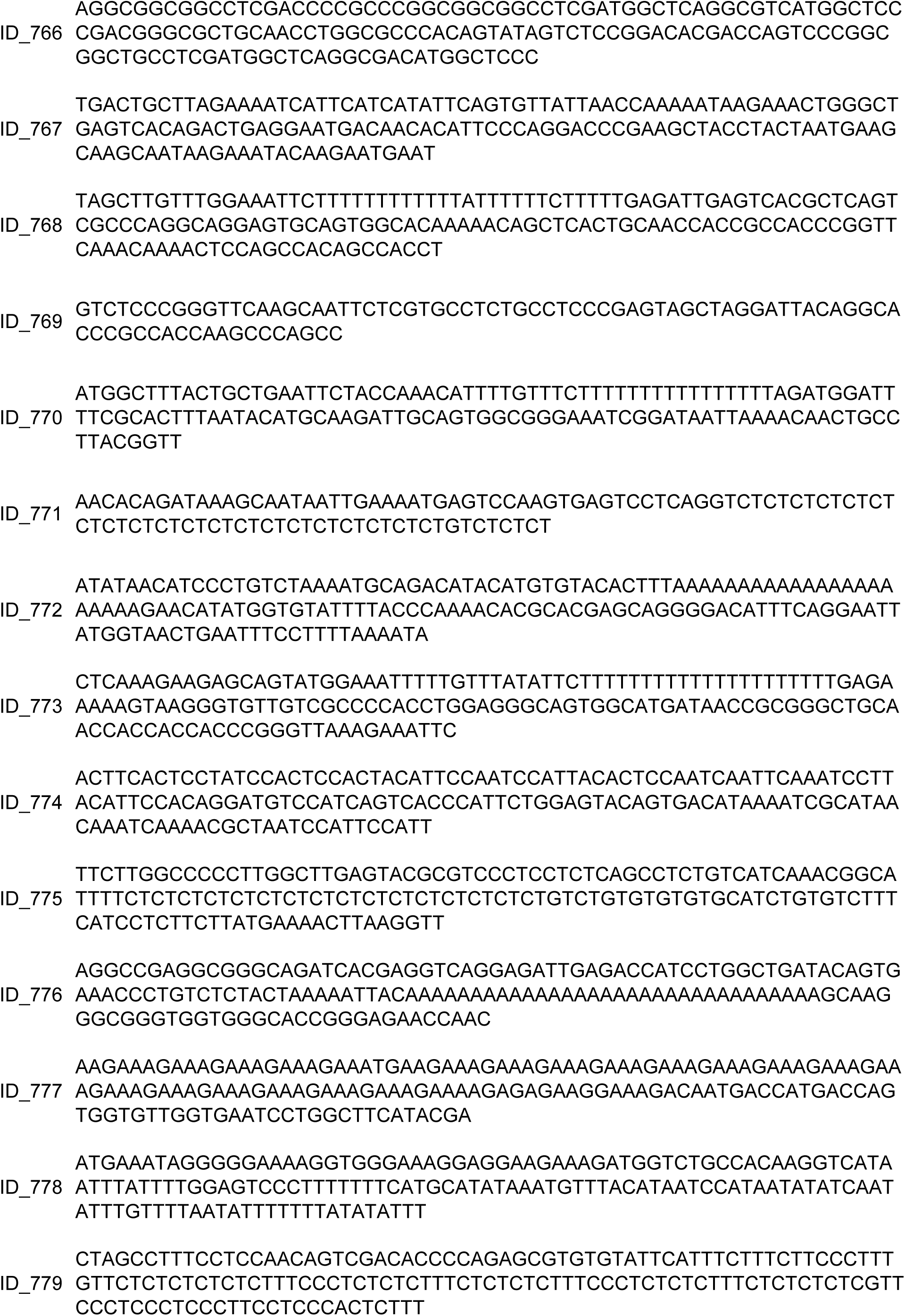
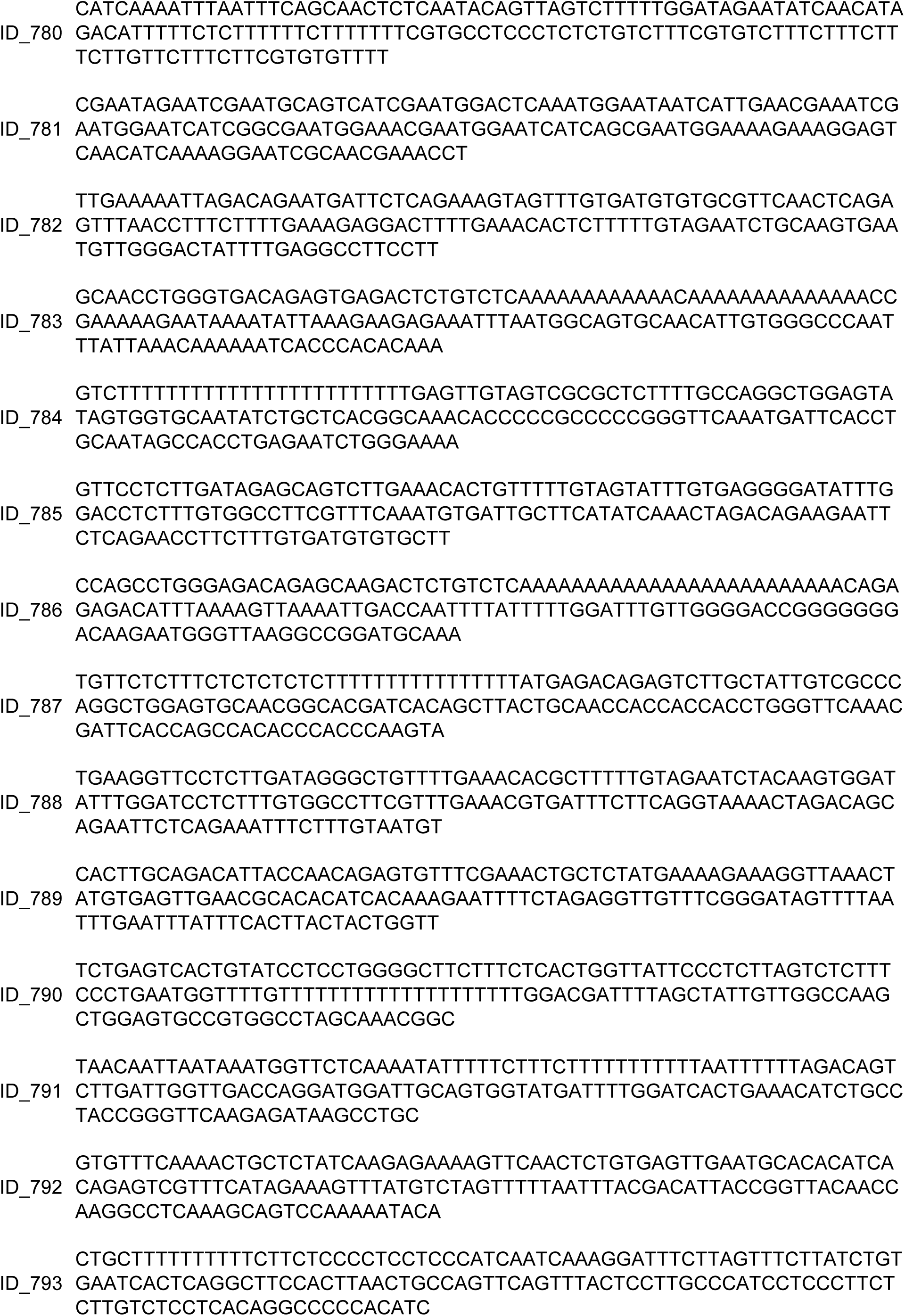
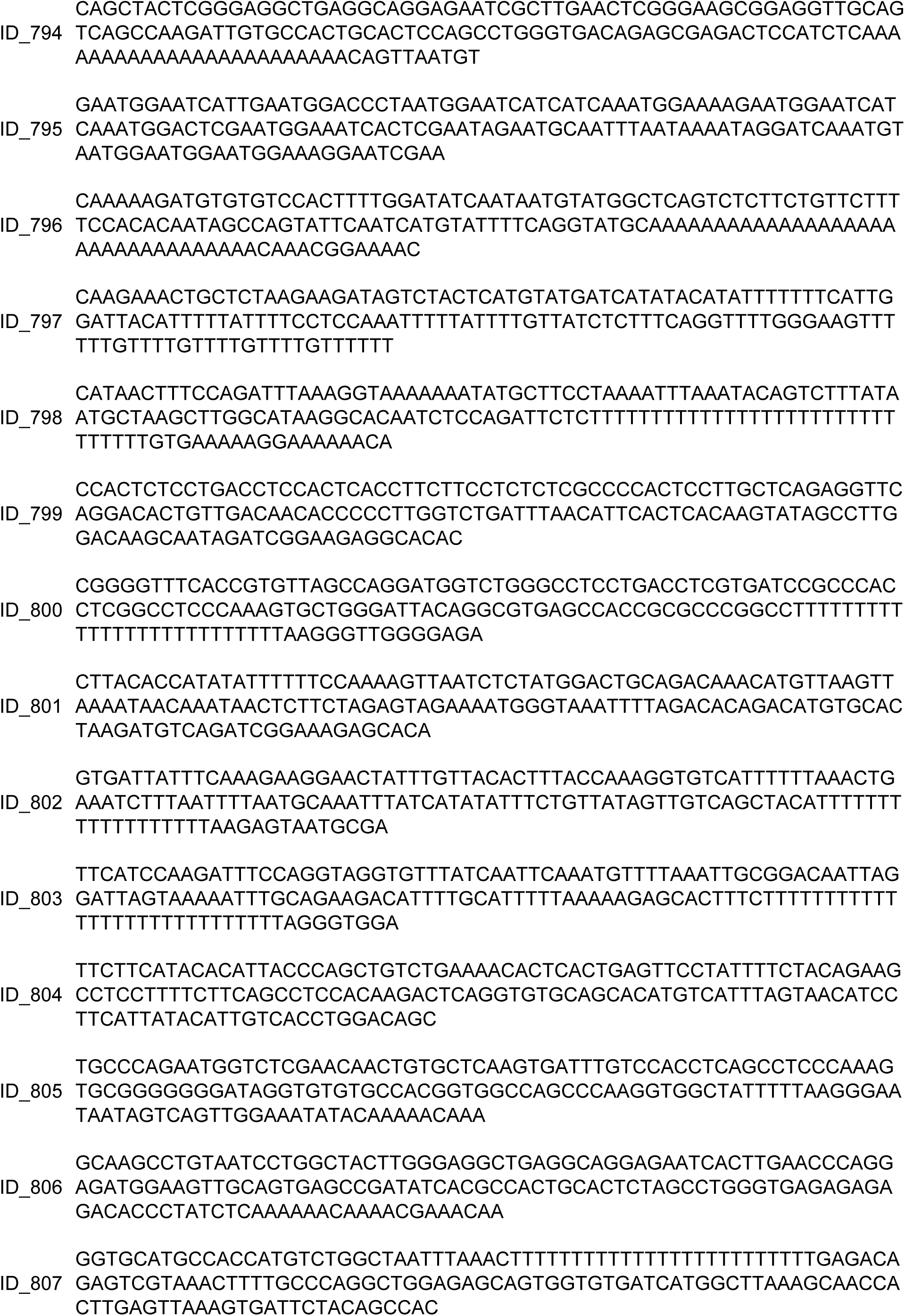
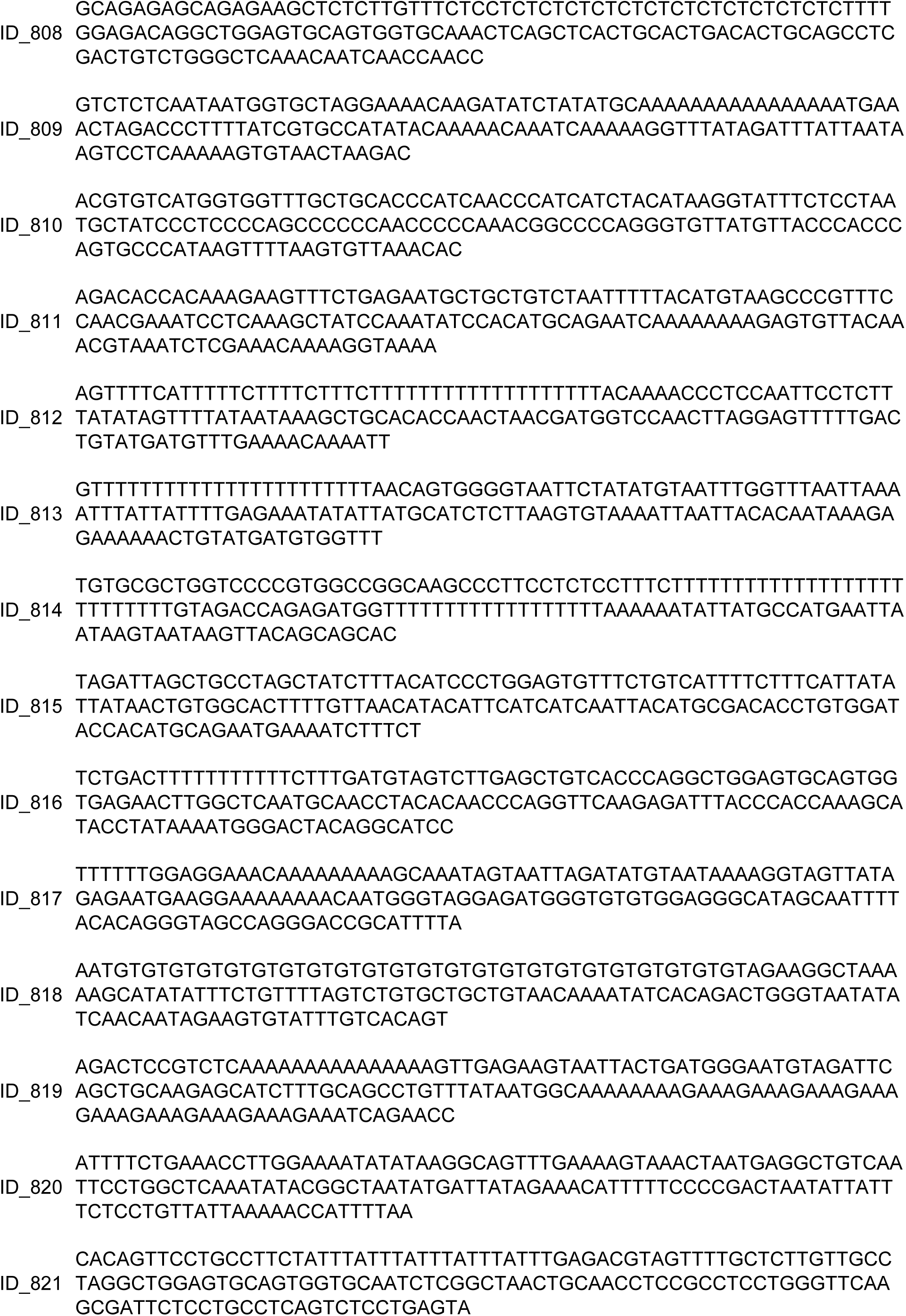
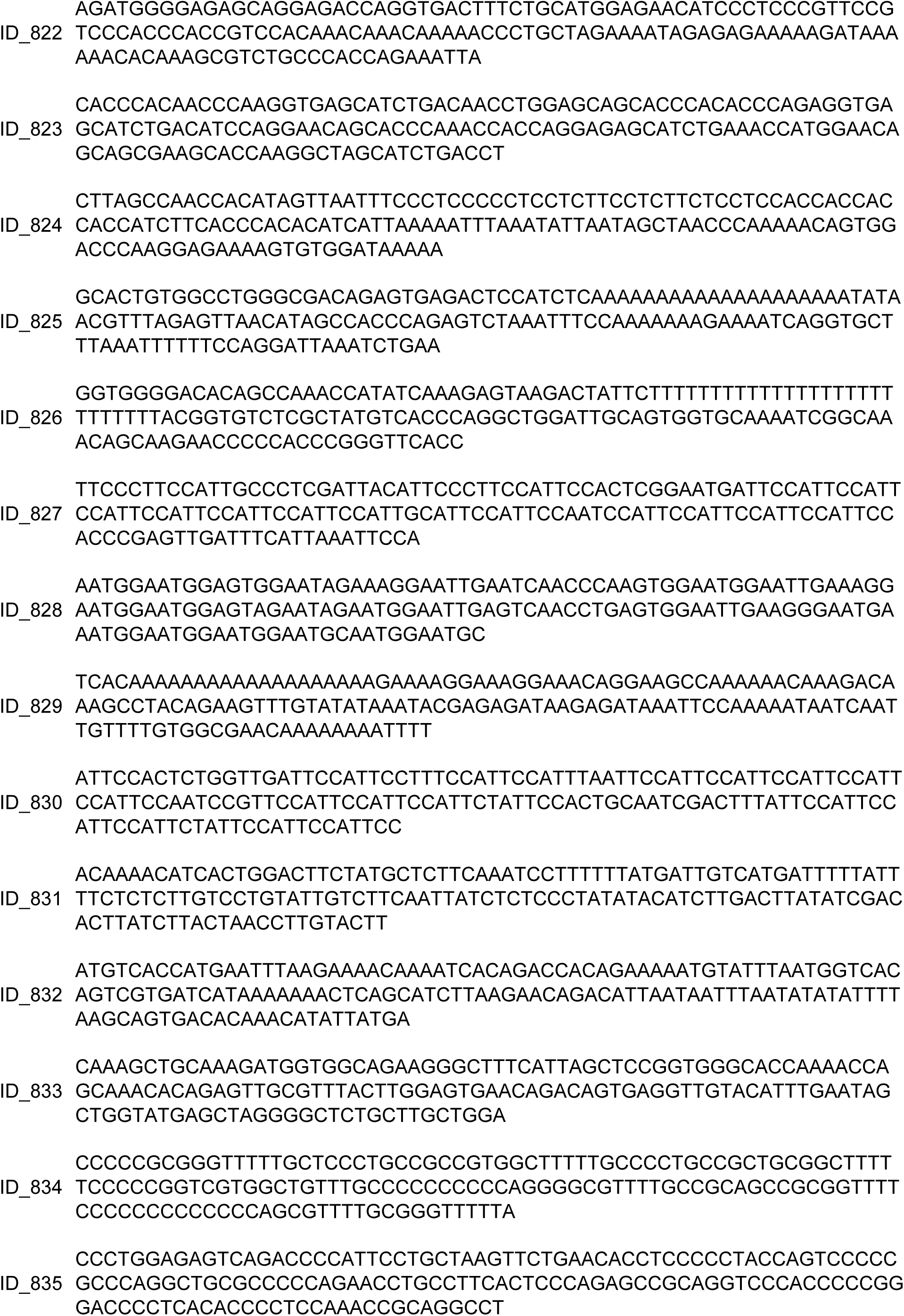
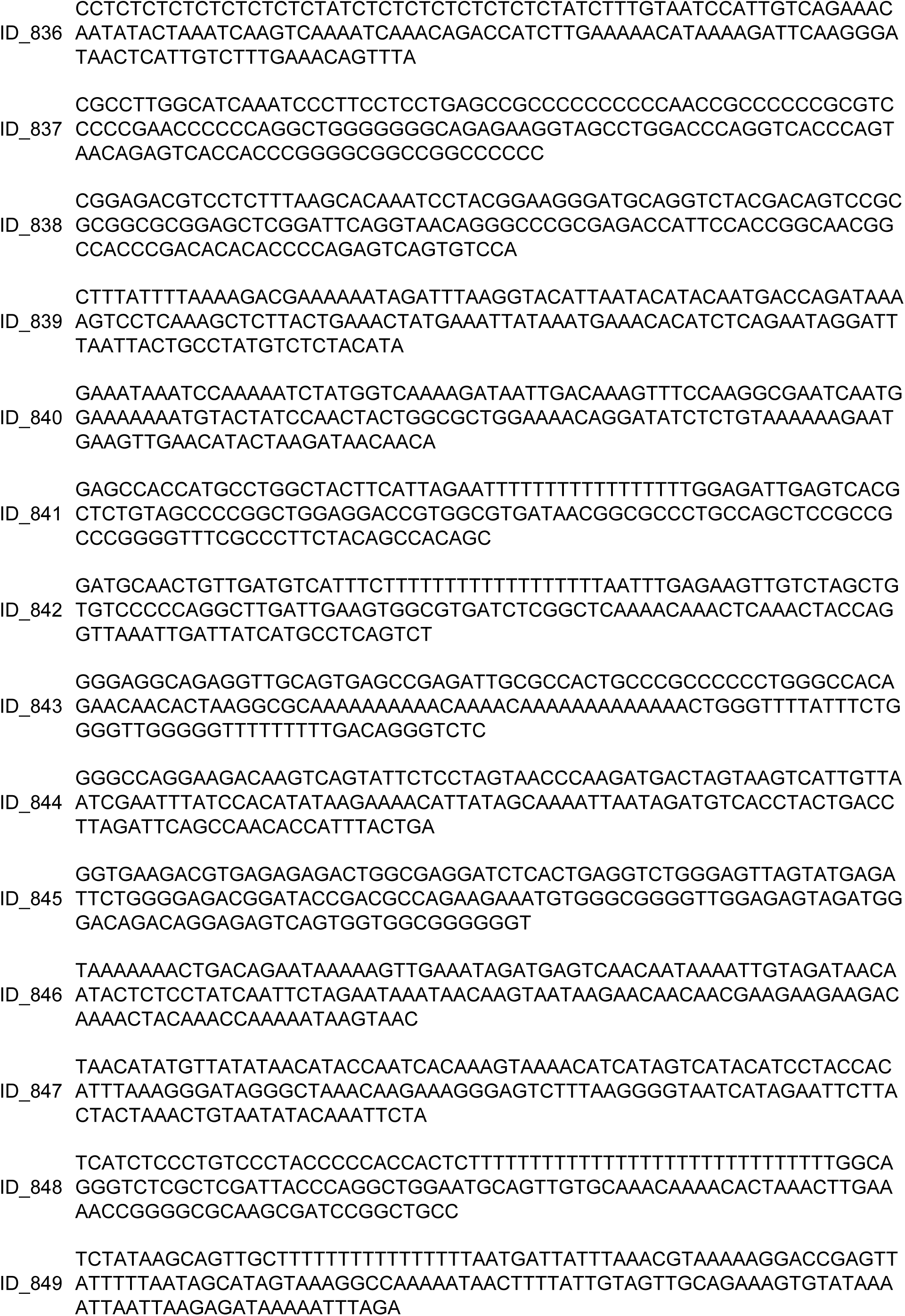
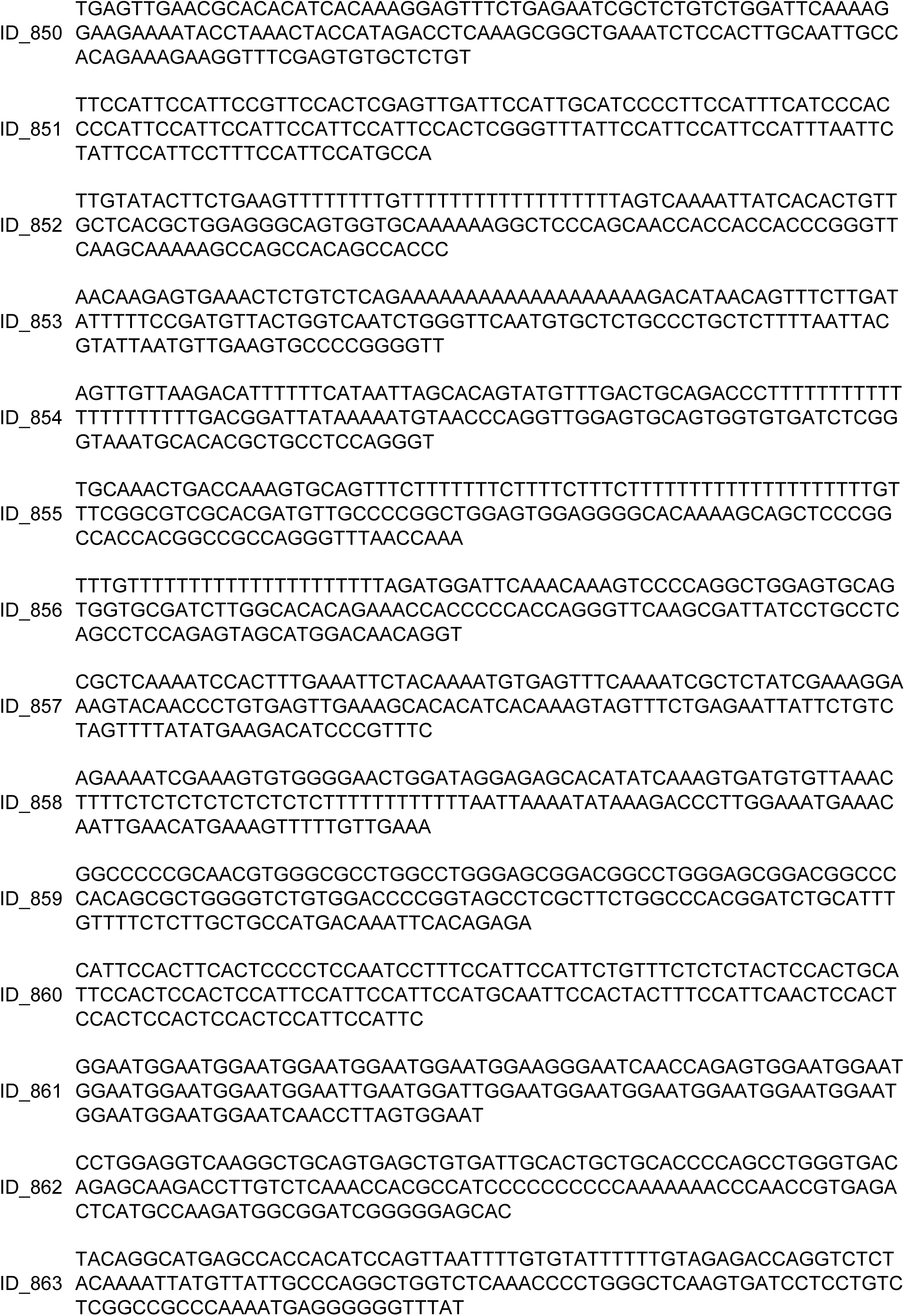
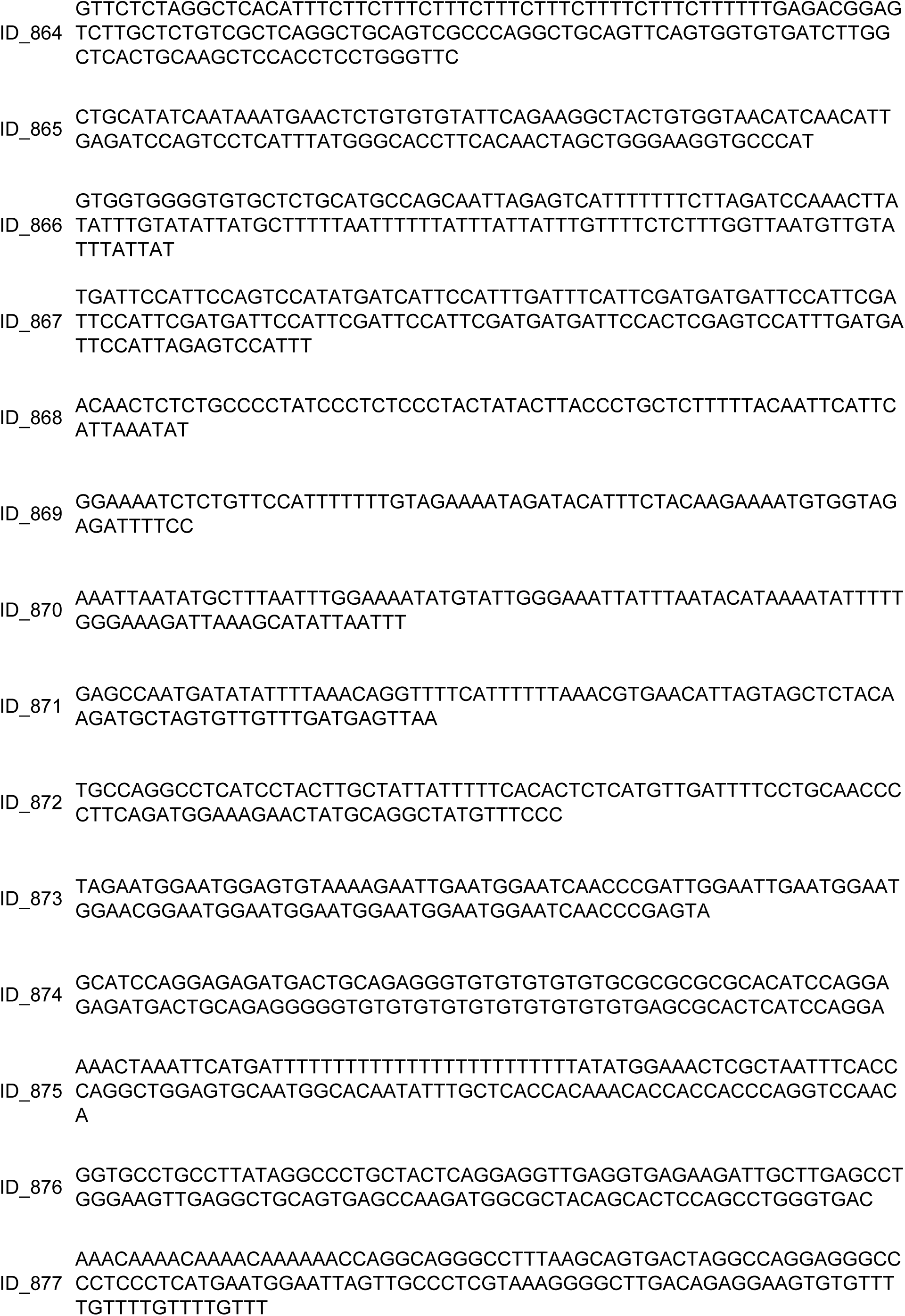
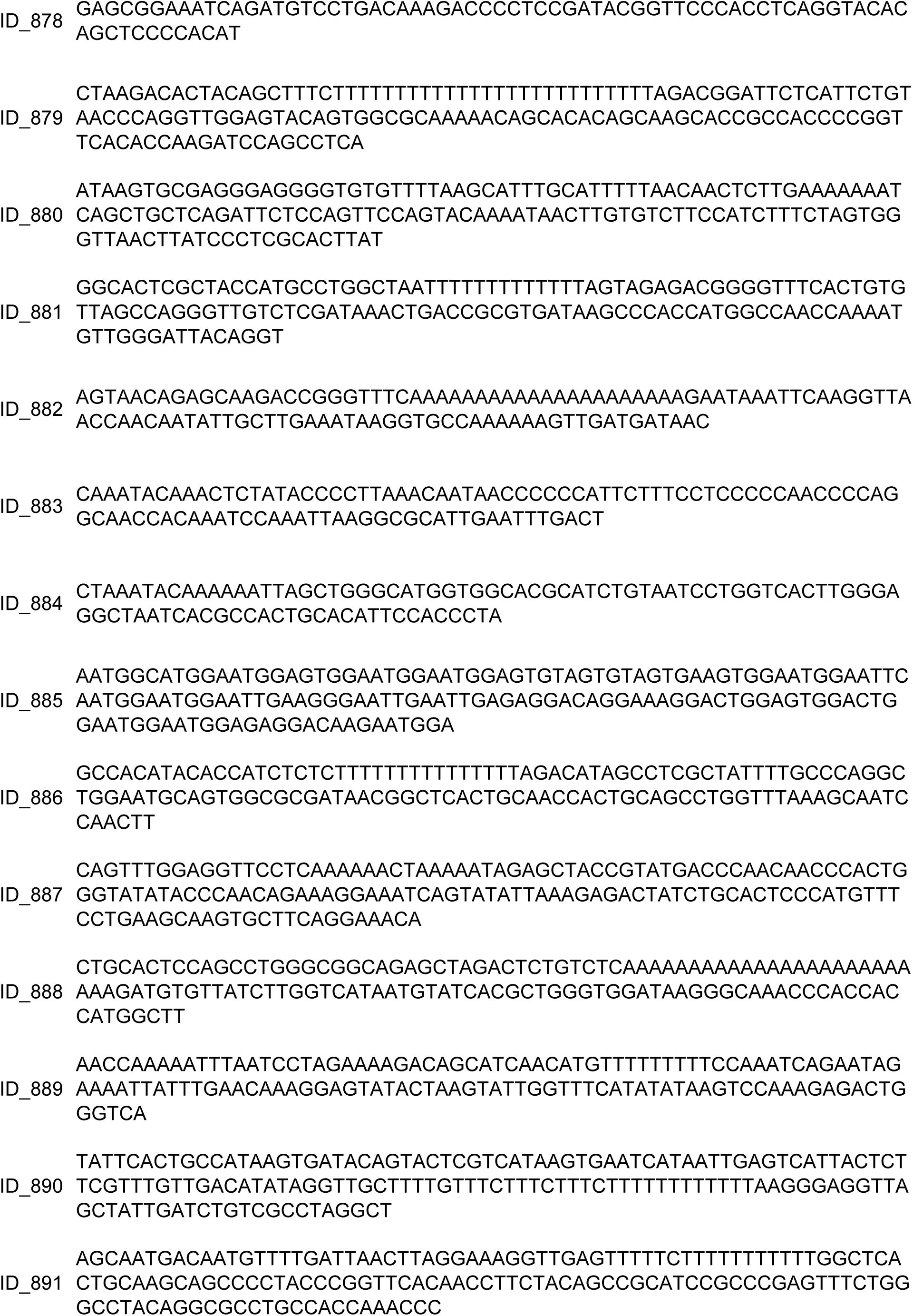
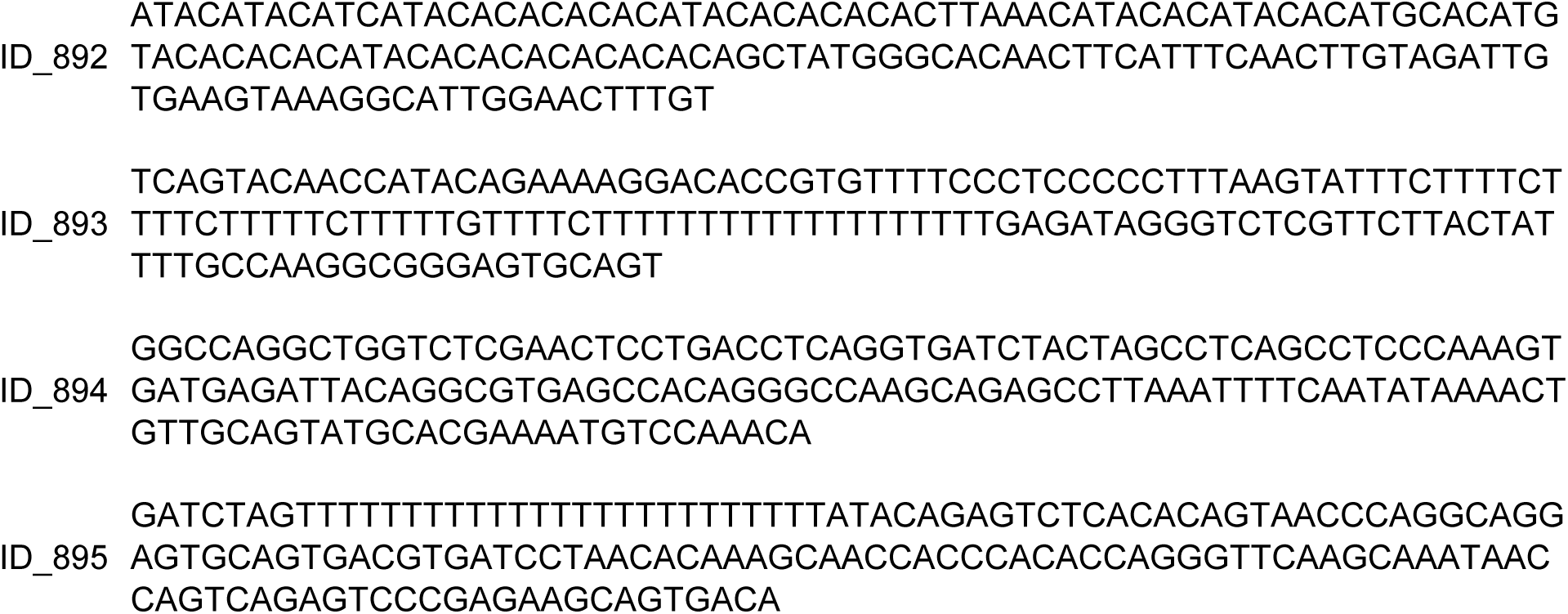
IDS Fragments in *Lycoptera* Fossil Records.

**Table S2A.**
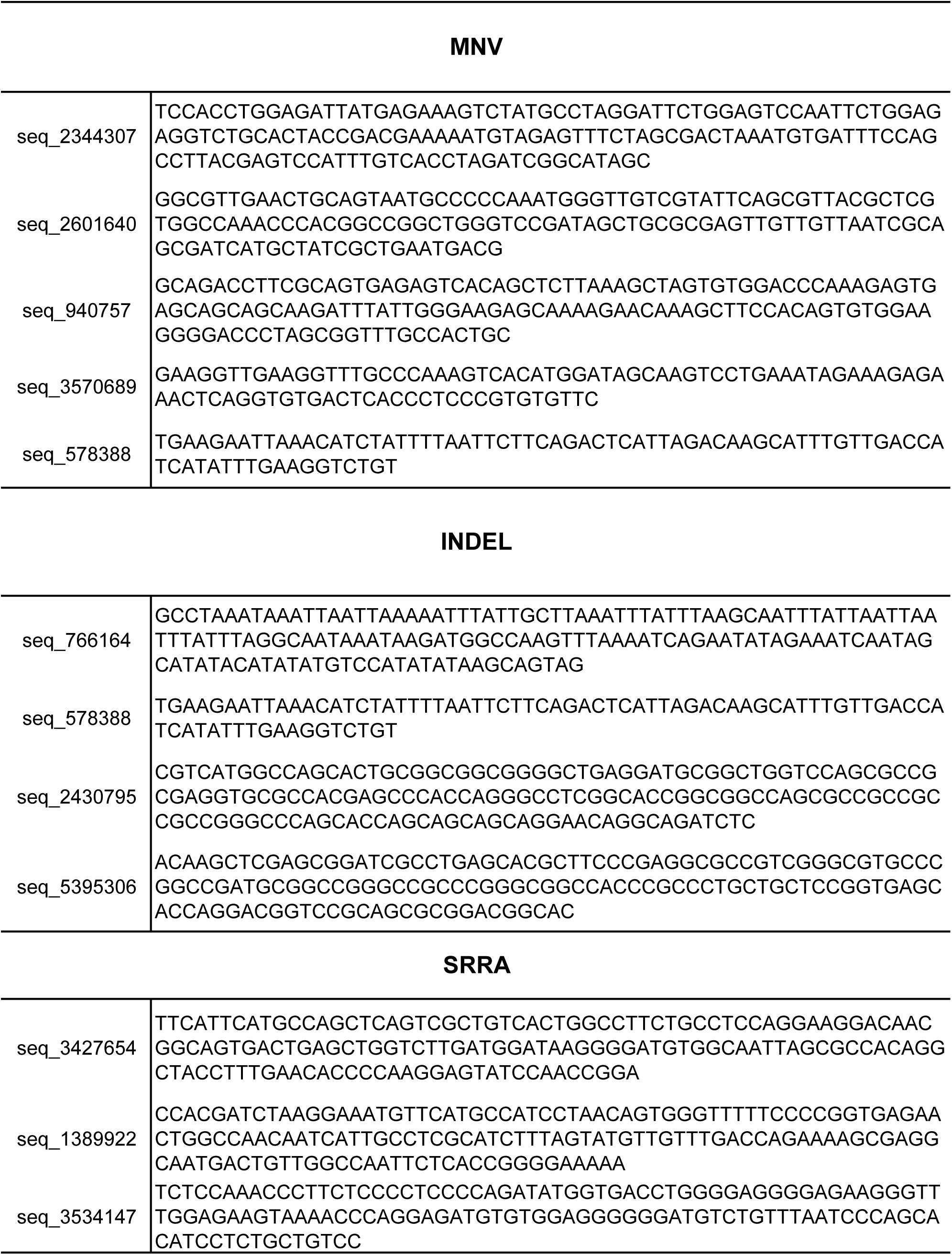
IDS Fragments in Round-Bellied Jars.

**Table S2B.**
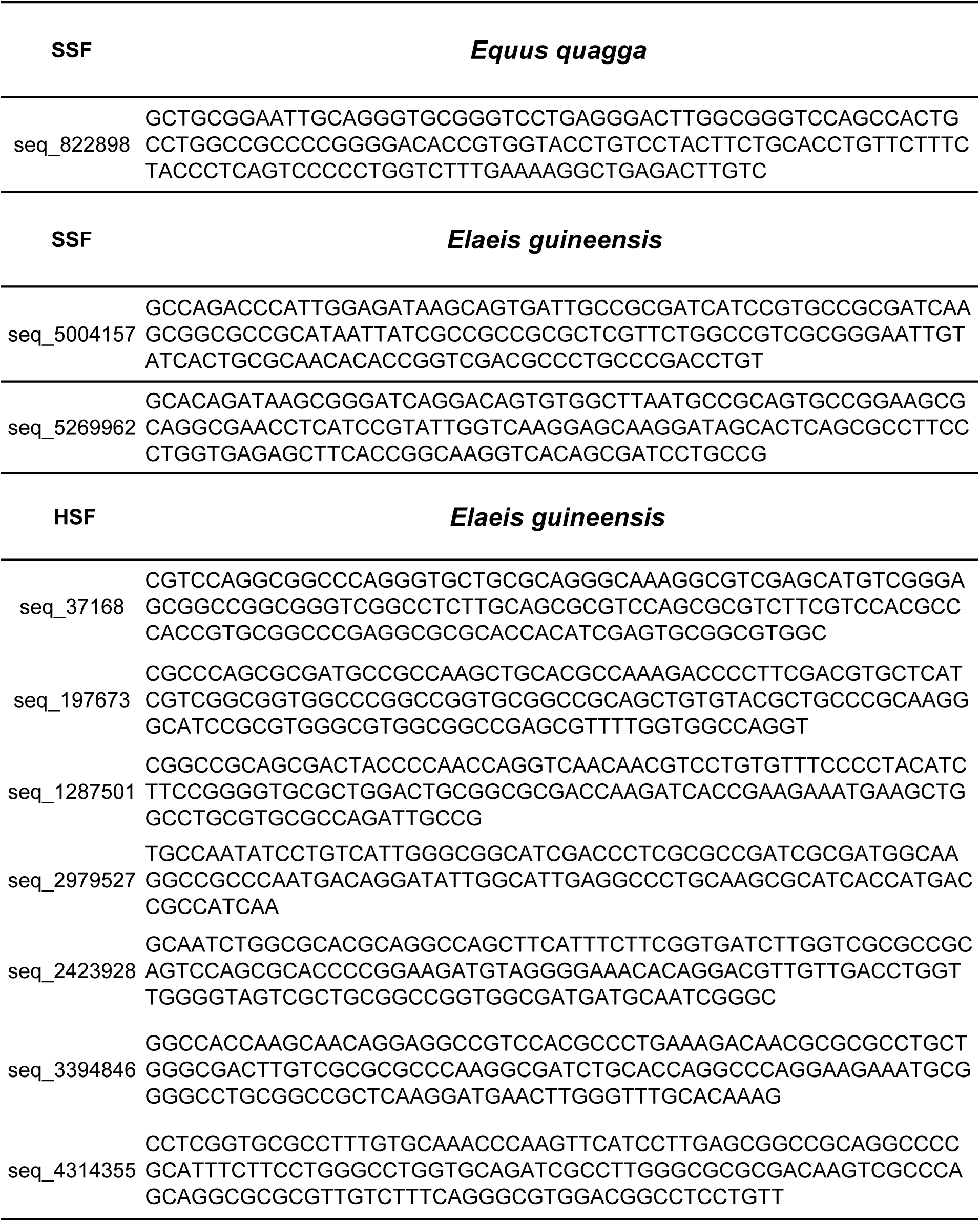
SSF Sequences in Round-Bellied Jars.

## References

01. Zhao, W. Q., et al. Ancient DNA in *Lycoptera* Fossils: Key Discoveries for Evolution and Methodological Reassessment (2023).

02. Gould, S. J. & Eldredge, N. Punctuated equilibria: The tempo and mode of evolution reconsidered. Paleobiology, 3(2), 115–151 (1977).

03. Vacca, F. Yalcin, B., Ansar, M. Exploring the pathological mechanisms underlying Cohen syndrome. Front Neurosci. 18:1431400 (2024).

04. Anholt, R. R. H. Olfactomedin proteins: central players in development and disease. Front Cell Dev Biol. 2:6 (2014).

05. Higuchi, R. et al. DNA sequences from the quagga, an extinct member of the horse family. Nature 312, 282–284 (1984).

06. Jónsson, H. et al. Speciation with gene flow in equids despite extensive chromosomal plasticity, Proc. Natl. Acad. Sci. U.S.A. 111(52),18655–18660 (2014).

07. Wang, X. G. Exploration on Ecological Environment Changes and the Rise of Xia Dynasty [In Chinese]. Science Press (2014).

08. Rightmire, G. P. Human evolution in the Middle Pleistocene: The role of *Homo heidelbergensis*. Evolutionary Anthropology: 6(6), 218–227 (1998).

09. Antón, S. C. Evolutionary significance of cranial variation in Asian Homo erectus. American Journal of Physical Anthropology 118(4), 301–323 (2002).

10. 10. Etler, D. A. Homo erectus in East Asia: Human Ancestor or Evolutionary Dead-End? Retrieved from https://www.academia.edu/ (2014).

11. Bae, C. J., Douka, K., Petraglia, M.D. On the origin of modern humans: Asian perspectives. Science. 358(6368), eaai9067 (2017).

12. Fu, Q. et al., DNA analysis of an early modern human from Tianyuan Cave, China. Proceedings of the National Academy of Sciences 110, 2223–2227 (2013).

13. Massilani, D., et al. Denisovan ancestry and population history of early East Asians. Science. 370 (6516), 579–583 (2020).

14. Zhao, W. Q., et al. Emerging Challenges in Molecular Paleontology: Misapplication of Environmental DNA Fragments and Misconception of Deamination as a Key Criterion for *In Situ* DNA Identification. arXiv. 2412.06378 (2024).

15. Xue, J. R. et al. The functional and evolutionary impacts of human-specific deletions in conserved elements. Science 380 (6643), eabn2253 (2023).

16. Lefébure, T. et al. Less effective selection leads to larger genomes. Genome Res. 27(6), 1016–1028 (2017).

17. Hedges, S. B., et al. Detecting Dinosaur DNA. Science. 268(5214), 1191 -1194 (1995).

